# Detection of five viruses commonly implicated with Bovine Respiratory Disease using loop-mediated isothermal amplification

**DOI:** 10.1101/2025.11.06.687045

**Authors:** Josiah Levi Davidson, Murali Kannan Maruthumuthu, Mohamed Kamel, Suraj Mohan, Ana Pascual-Garrigos, Andres Dextre, Ruth Eunice Centeno-Delphia, Jacquelyn P. Boerman, Deepti Pillai, Jennifer Koziol, Aaron Ault, Jon P. Schoonmaker, Timothy A. Johnson, Mohit S. Verma

**Affiliations:** Department of Agricultural and Biological Engineering, Purdue University, West Lafayette, IN, USA, 47907; Birck Nanotechnology Center, Purdue University, West Lafayette, IN, USA, 47907; Department of Biochemistry, Purdue University, West Lafayette, IN; Department of Medicine and Infectious Diseases, Faculty of Veterinary Medicine, Cairo University, Giza, 11221, Egypt; Department of Animal Science, Purdue University, West Lafayette, IN, USA, 47907; Department of Comparative Pathobiology, Purdue University, West Lafayette, IN, USA, 47907; Department of Veterinary Clinical Science, Purdue University, West Lafayette, IN, USA, 47907; School of Veterinary Medicine, Texas Tech University, Amarillo, TX, USA, 79106; Department of Electrical and Computer Engineering, Purdue University, West Lafayette, IN, USA, 47907; Weldon School of Biomedical Engineering, Purdue University, West Lafayette, IN, USA, 47907

**Keywords:** Bovine Respiratory Disease, LAMP, viral BRD, chute-side diagnostics, BHV-1, BRSV, BVDV-1, BAV-3, bPIV-3, assay development, minimal sample processing, extraction-free detection

## Abstract

Herein, we present novel quantitative loop-mediated isothermal amplification (qLAMP) and reverse-transcription qLAMP (RT-qLAMP) assays for the detection of five viruses commonly implicated with the onset and progression of bovine respiratory disease (BRD): Bovine Alphaherpesvirus Type 1 (BHV-1), Bovine Adenovirus Type 3 (BAV-3), Bovine Respiratory Syncytial Virus (BRSV), Bovine Viral Diarrhea Virus Type 1 (BVDV-1), and Bovine Parainfluenza Virus Type 3 (BPIV-3). Using contrived samples spiked with whole viruses, we found that our extraction-free assays have limits of detection between 30 and 1,057 copies per reaction (1.8% final sample concentration) with minimal sample processing. Using dual-tipped swabs and 1.4 mL resuspension volumes, these limits of detection are on the order of 2 × 10^5^ copies per swab for BAV-3 and BHV-1 and between 6.31 × 10^6^ to 8.22 × 10^6^ copies per swab in the case of BPIV-3, BRSV, and BVDV-1. Analytical sensitivities ranged from 73 – 100% and analytical specificities ranged from 90 – 100%. Additionally, we introduced a streamlined pipeline to minimize the experimental workload to design, screen, select, and characterize LAMP performance for developing assays. The assays targeting these BRD viruses can be utilized to develop colorimetric LAMP assays that enable the sensitive and specific detection of these viruses’ chute side to aid in diagnosing and treating BRD. The associated development pipeline enables more rapid development of LAMP-based diagnostic tools targeting emerging pathogens.

## 1. Introduction

Bovine respiratory disease (BRD) is a multifactorial disease complex that has substantial economic impacts on the beef and dairy industries worldwide. Several viral and bacterial pathogens cause BRD but many of those pathogens do not have targeted diagnostics available for field-based usage (Blakebrough-Hall et al. 2020; Gaudino et al. 2022). As a result, caretakers frequently use blanket antimicrobial treatments to treat cattle due to the inability to treat these potential pathogens in a targeted manner, thus giving rise to antimicrobial resistance infections (Murray et al. 2016; Nickell et al. 2021). Chute-side detection of BRD-associated bacteria using loop-mediated isothermal amplification (LAMP) is emerging as a promising method for targeted treatment in field settings (Pascual-Garrigos et al. 2021). Where specific therapy does not exist even when the cause is known, chute-side LAMP assays can help sequester animals and prevent spread to nearby livestock. However, due to a lack of sufficiently developed LAMP assays targeting these viruses, researchers have not employed these field-based detection methods for the diagnosis of BRD.

BRD prevalence varies between 2 – 20% and is affected by geography, climate, diagnostic method, and sampling method (O’Donoghue et al. 2025). Furthermore, viral infection typically precedes bacterial co-infection placing the cattle at a higher risk for development of more dire clinical conditions such as enzootic pneumonia (Sarmiento-Silva et al. 2012; Gaudino et al. 2022). Recent literature investigating the interplay between viral and bacterial infections in BRD epidemiology, however, show that BRD viruses may play a more important role in epidemiological spread in younger calves than in older calves (Calderón Bernal et al. 2023).

BRD is frequently associated with Bovine Adenovirus Type 3 (BAV-3), Bovine Herpesvirus Type 1 (BHV-1), Bovine Parainfluenza Virus Type 3 (bPIV-3), Bovine Respiratory Syncytial Virus (BRSV), and Bovine Viral Diarrhea Virus Type 1 (BVDV-1) infections (Fulton 2020). Buczinski, Calderon Bernal, O’Donoghue, Shen, and Studer, provide four studies of the prevalence of the viruses we study here in various regions (Shen et al. 2020; Studer et al. 2021; Calderón Bernal et al. 2023; Buczinski et al. 2024; O’Donoghue et al. 2025). Additionally, morbidity rates of animals infected with these viruses and subsequently displaying symptoms can be as high as 80% in the case of infection with BRSV (Gaudino et al. 2022). Similarly, the frequency of detection of viruses studied herein also varied by pathogen and region. BRSV prevalence varied across 2%, 18%, and 20% in Switzerland, Canada, and Spain, respectively, when testing samples using quantitative reverse transcription polymerase chain reaction (qRT-PCR; (Studer et al. 2021; Calderón Bernal et al. 2023; Buczinski et al. 2024). In Canada, BSRV prevalence increased to as high as 21% when isolating the virus from samples (Buczinski et al. 2024). bPIV-3 varied across 3%, 4%, and 26% in Switzerland, Canada, and Spain, respectively (Studer et al. 2021; Calderón Bernal et al. 2023; Buczinski et al. 2024). BVDV-1 varied between 1% in Canada and 17% in Spain, while BHV-1 had a frequency of detection of 3% in both Canada and Spain (Studer et al. 2021; Calderón Bernal et al. 2023). BAV-3 is widespread in many regions of the world, and in one study conducted in China, BAV-3 was detected at a rate of 79% (Shen et al. 2020). As seen, the frequency of detection for a single virus has a large variance depending on the geographic region and a somewhat smaller variance depending on the diagnostic method.

Trained personnel typically conduct daily visual inspections of cattle to diagnose BRD (Buczinski et al. 2014; Timsit et al. 2016). Due to the complex nature of BRD, no gold standard for the antemortem diagnosis of BRD exists complicating the detection of these viruses (Churchill et al. 2025). Rather, caretakers identify at-risk cattle using visual methods such as DART to evaluate a cattle’s behavior, feeding patterns, and potential signs of respiratory distress (Juge et al. 2022; Kamel et al. 2024). Often, these methods vary from animal to animal and do not present consistently. Detection of BRD-associated viruses is further complicated by the differing structures of the viral nucleic acids and may consist of double-stranded DNA (dsDNA) for BAV-3 and BHV-1 or single-stranded RNA (ssRNA) for bPIV-3, BRSV, and BVDV-1. Accordingly, assays targeting these viruses must consider the possibility of a ssRNA genome and include reverse transcription. Traditional laboratory tests such as quantitative polymerase chain reaction (qPCR) and Reverse-Transcription qPCR (RT-qPCR) can improve the diagnosis of BRD and subclinical BRD but are typically costly (Griffin 2014; Pratelli et al. 2021). Furthermore, traditional laboratory tests require transportation to a laboratory and specialized equipment practically doubling the time required to get a result in ideal scenarios (Figure 1). As a result, caretakers need methods with diagnostic performance comparable to qPCR/RT-qPCR that can be conducted in field settings to effectively guide treatment of infections and BRD outbreaks in a targeted manner.

**Figure 1:**
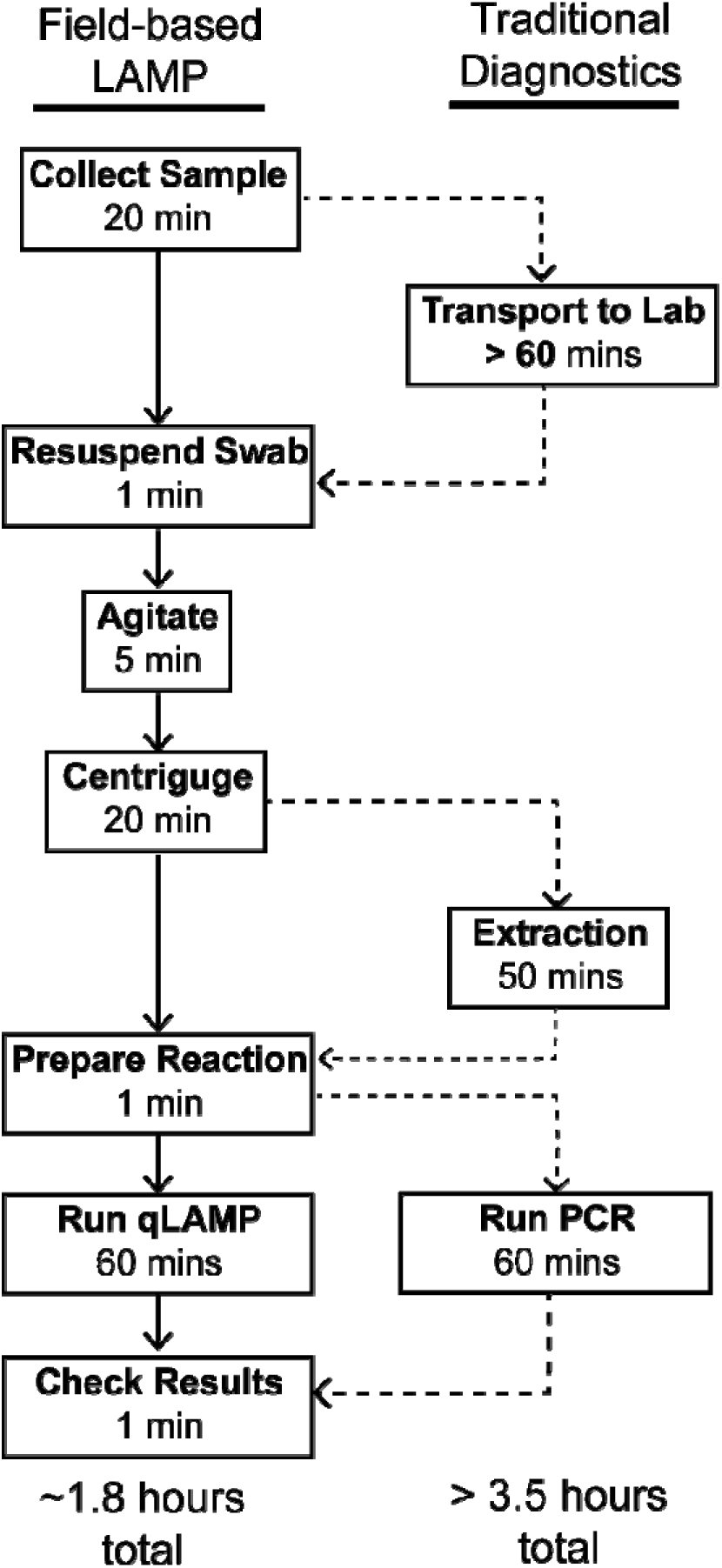
Comparison between field-based LAMP workflow presented in this study and the workflow required when using traditional diagnostics. Dashed connectors indicate additional processes that must be completed for traditional diagnostics.

LAMP is one such technique that allows for the sensitive detection of pathogens chute-side and in field settings (Pascual-Garrigos et al. 2021; Ahmed et al. 2025; Rafiq and Verma). We have previously compared isothermal methods for field-based applications in a review paper (Wang et al. 2022). Compared to other isothermal assays, LAMP has a higher resistance to inhibitors found in complex samples, making it a desired reaction for running in rudimentary settings outside of the lab (Wang et al. 2022). Furthermore, LAMP is amenable to a “single pot” reaction since it typically does not require preparation of the nucleic acid template to initiate amplification. This contrasts with other isothermal amplification assays, such as rolling circle amplification (RCA), which require pre-ligating sequences to form a circular template before amplification begins, thus increasing the complexity of the assay (Wang et al. 2022). Assays employing LAMP for the detection of bacterial pathogens associated with BRD show promising results when used in field-based settings with complex samples (Peltzer et al. 2021; Mohan et al. 2021; Pascual-Garrigos et al. 2021; Davidson et al. 2021; Centeno-Martinez et al. 2022; Ayaz Kök et al. 2023). Additionally, LAMP has comparable results to traditional lab-based diagnostic techniques, but without the added expense of costly equipment and trained personnel.

The primary drawback of LAMP, however, is its primer design, which is markedly more complex than other isothermal amplification schemes generally requiring only 2 – 4 primers (Wang et al. 2022). Due to the complexity of the LAMP reaction requiring a total of six primers targeting eight regions, the development of LAMP primer sets typically requires the iterative design and experimental screening of multiple primer sets (Huang et al. 2022). This screening can be both time- and resource-intensive, and to our knowledge, the literature does not substantially address guidelines for a LAMP-specific experimental screening and selection process.

We have previously developed LAMP assays to detect bacterial pathogens associated with BRD, but similar LAMP assays are not sufficiently developed in the literature to detect BRD-associated viruses in a timely and robust manner (Mohan et al. 2021). When assays are available for certain viruses associated with BRD, users generally need to extract nucleic acids from samples in a laboratory environment rendering the assay unsuitable for field-based usage. Table 1 summarizes assays found in the literature that target the viruses we investigate in this study. To the best of our knowledge, these assays are the best-developed LAMP assays available for the indicated virus, and all require extraction prior to testing with LAMP. The LAMP assay targeting BHV-1 has a reported reaction time of 8 – 30 mins; however, using the assay with crude, unextracted field samples resulted in false positives. Consequently, users had to perform rudimentary extraction and pre-treatment to sufficiently detect BHV-1 (Peltzer et al. 2021). The bPIV-3 LAMP assay required users to extract and reverse transcribe genomic RNA (gRNA) in a separate reaction to produce complementary DNA (cDNA) prior to conducting the LAMP assay. This assay had a limit of detection (LOD) of approximately 416 copies per reaction but this LOD was not confirmed in complex samples (Li Jian-you et al. 2017). Similarly, the LAMP assay targeting BRSV required users to reverse transcribe gRNA to cDNA prior to detection with LAMP, resulting in an LOD of approximately 2.4 × 10^4^ copies per reaction (Li et al. 2019). The BVDV-1 LAMP assay does not include loop primers, which are known to accelerate the reaction time by at least twofold, resulting in a reaction that requires a full 60 mins to return a result. Furthermore, Fan et al. did not validate the BVDV-1 assay with field samples and thus its performance in complex sample is unknown (Fan et al. 2012). At the time of publication, we could not identify any existing LAMP or RT-LAMP assays targeting BAV-3. Therefore, we included a developed qPCR assay targeting BAV-3 in this study for comparison. As with many qPCR assays, users must first extract genomic content from samples prior to using the assay. Moreover, the assay displayed an LOD of 25 copies per reaction when the sample background contained a spike-in target (Wong and Xagoraraki 2010).

**Table 1:** Selection of currently available LAMP or RT-LAMP assays in literature targeting viruses investigated during this study. In the case of BAV-3, no LAMP assays were found resulting in a qPCR assay used as comparison to the LAMP assay from this study. The template used in the reaction, whether that reaction was also tested using a complex sample, whether nucleic acid extraction is needed for the assay, and the limit of detection (LOD) of the assay are listed and compared to the assay developed during this study to target that virus.

| Virus | Reaction Type | Template Type |  | Confirmed in Complex Sample? |  | Extraction Needed? |  | LOD (copies/reaction) |  | Ref. |
| --- | --- | --- | --- | --- | --- | --- | --- | --- | --- | --- |
|  |  | Literature | This Study | Literature | This Study | Literature | This Study | Literature | This Study |  |
| **BAV-3** | qPCR | Plasmid cDNA | Whole Virus from Cell Culture | Yes (cDNA spiked into fecal negative extract) | Yes. Spiked into nasal swabs | Yes | No. Only dilution in water | 25 | 38 | (Wong and Xagorarakis 2010) |
| **BHV-1** | LAMP | Extracted DNA |  | Yes (using DNA precipitation to extract DNA) |  | Yes |  | 14 | 21 | (Peltzer et al. 2021) |
| **bPIV-3** | RT-LAMP | cDNA |  | No |  | Yes |  | 416 | 1,057 | (Li Jian-you et al. 2017) |
| **BRSV** | RT-LAMP | cDNA | | No | | Yes | | $\sim 2.4 \times 10^4$ | 811 | (Li et al. 2019) |
| **BVDV-1** | RT-LAMP | Extracted RNA |  | No (all samples were Extracted) |  | Yes |  | 4.67 | 1,004 | (Fan et al. 2012) |

Herein, we report five novel LAMP assays for the detection of BAV-3, BHV-1, bPIV-3, BRSV, and BVDV-1 in contrived minimally processed field samples (shallow nasal swabs from cattle, resuspended in water) using a comprehensive process to design, screen, select and characterize LAMP assay performance. We then characterize these LAMP assays and report the LOD, analytical specificity, and analytical sensitivity are reported for each assay. Furthermore, to ensure confidence in the characterization and performance of our assay, we quantify target viral concentration in parallel with digital PCR (dPCR). Finally, we report the cross-reactivity of each LAMP assay with the other viruses investigated in this study.

The LAMP assays designed and presented here form the foundation for a field-based detection tool for determining the presence of five viruses commonly implicated with BRD. Researchers can further develop these LAMP assays for use in a field-based colorimetric LAMP detection assay chute, as Pascual-Garrigos et al. previously demonstrated for the detection of BRD bacteria chute-side (Pascual-Garrigos et al. 2021). Additionally, assay developers can utilize the process we developed and documented herein to accelerate the design and characterization of novel LAMP assays for other pathogens or targets of interest in animal health.

## 2. Materials and Methods

### 2.1. Host Cell Propagation

Mavin Darby bovine kidney (MDBK) cells (NBL-1 Strain; ATCC CCL-22; Manassas, VA, USA) were cultured in DMEM or DMEM/F12 with 4 g/L glucose and 2 mM L-glutamine supplemented with 10% Fetal Bovine Serum (FBS) which tested negative for anti-BHV-1, anti-BVDV-1, anti-BRSV, anti-BPIV-3, and anti-BAV-3 antibodies. Bovine Turbinate (BT) cells (ATCC CRL-1390; Manassas, VA, USA) were cultured in Dulbecco’s Minimum Essential Media (DMEM) with 4 g/L glucose and 2 mM L-glutamine supplemented with 10% Horse Serum (HS). Cells were incubated at 37.0 °C and 5% CO_2_ and grown to 80 – 90% confluency prior to infection with the virus.

### 2.2. Virus Propagation

Before infection, cells were washed three times with 5 – 10 mL of DPBS without calcium or magnesium. Media for virus propagation was modified from that used for the respective host cell propagation such that the final concentration of FBS was 2%. 10 mL of the resulting 2% serum-supplemented media was added to the monolayer after washing and prior to infection. BAV-3 (WBR-1 Strain; ATCC VR-639; Manassas, VA, USA), BHV-1 (Los Angeles (LA) Strain; ATCC VR-188), and BVDV-1 (NADL Strain; ATCC VR-534) were grown in MDBK cells (NBL-1; ATCC CCL-22). BRSV (A51908 Strain; ATCC VR-1861) and BPIV-3 (SF-4 Strain; BEI Resources NR-3234; Manassas, VA, USA) were grown in BT cells (ATCC CRL-1390; Manassas, VA, USA). MDBK and BT host cells were not permitted to be subcultured more than six times. MDBK cells were infected with 200 – 400 μL of the indicated virus and BT cells were infected with 200 μL of the indicated virus.

BRSV required a blind passage at approximately 10 days post-infection wherein cells were detached from the plate using Accutase® or equivalent (Corning 24-058-CI; Corning, NY, USA), mixed in a 1:1 ratio with uninfected BT cells (detached in the same manner), and centrifuged at 400 × g for seven mins. The supernatant was removed, and the resulting pellet was resuspended in 1 mL of infection media. 500 uL of the resulting cell mixture was used to re-seed a new T-75 plate with 10 mL of media.

Cells were monitored daily for cytopathic effects (CPE). Viruses were harvested when approximately 80% or more of the monolayer was destroyed. Culture supernatant was centrifuged at 14,000 RCF for 20 mins at 10 °C. The resulting supernatant was decanted, aliquoted, and stored at -80 °C until ready for use. BAV-3 and BRSV required two freeze-thaw cycles whereas in cell culture, the media was flash frozen at -80°C and thawed at 37 °C.

### 2.3. Nucleic acid extraction

Viral DNA or RNA was extracted from cell culture supernatant using PureLink™ Viral RNA/DNA Mini Kit (ThermoFisher Scientific 12280050; Waltham, MA, USA) according to the manufacturer’s recommendations using 200-400 µL of viral culture supernatant with the following modifications: i) the second wash step was centrifuged at maximum speed for 3 mins, ii) elution was conducted using room-temperature nuclease-free water and incubated on the column at room temperature for 3-5 mins before eluting. Nucleic acid extracts were stored at -80 °C until use.

### 2.4. Synthesis of primers and probes

All primers were purchased from Integrated DNA Technologies (IDT; Coralville, IA, USA) and were purified using standard desalting. Primers were rehydrated in nuclease-free water to 100 µM by the manufacturer and shipped on dry ice.

All probes were purchased as PrimeTime® 5’ nuclease probes from IDT and were purified using HPLC. Probes were labeled with a 6-FAM™ 5’ fluorophore and a 3’ Iowa Black® Fluorescent quencher with an internal ZEN™ quencher. Probes were shipped dehydrated and, upon receipt, resuspended in nuclease-free water to a concentration of 100 µM. Probes were stored at -20 °C protected until usage. Sequences for the primers and probes used for dPCR are reported in Table S1 and sequences for LAMP/RT-LAMP primers used in this study are provided in Supplemental File 2.

### 2.5. Nucleic acid quantification via digital polymerase chain reaction

Sequences for the primers and probes are reported in Table S1. Digital PCR (dPCR) was used to quantify nucleic acid extracts using primers and probes previously reported in the literature for qPCR. Specifically, for detection of BAV-3, primer set BAV-3.Kishimoto.Hexon was used; for BHV-1, primer set BHV-1.Chandranaik.gB was used; for detection of bPIV-3, primer set BPI3ii.Thonur was used; for detection of BRSV, primer set BRSV.Boxus.N was used; and for detection of BVDV-1, primer set BVDV-n was used (Boxus et al. 2005; Thonur et al. 2012; Chandranaik et al. 2013; Mari et al. 2016; Kishimoto et al. 2017). All primer sets contained a forward and reverse primer (indicated by “F” and “R” appended to the primer set in Table S1) and a probe (indicated by a “P” appended to the primer name in Table S1).

dPCR was conducted on a Qiagen QIAcuity One, 5 Plex System (Qiagen 911022; Germantown, MD, USA) and unless otherwise indicated, was carried out in QIAcuity Nanoplate 26k 24-well (Qiagen 250031). A 10x primer-probe mix comprised 4 µM of both the forward and reverse PCR primers and 2 µM of the probe.

Nucleic acid extracts from BRSV, BPIV-3, and BVDV-1 were quantified using the QIAcuity OneStep Advanced Probe Kit (Qiagen 250132). 44 µL reactions consisting of 11 µL of 4X Probe PCR Master Mix, 0.44 µL of 100x OneStep Advanced RT, 4.4 µL of 10x Primer-probe mix formulated as described above, 5.5 µL of GC Enhancer, 11.66 µL of nuclease-free water, and 11 µL of a 1:1,000 dilution of the desired nucleic acid extract. 42 µL of this reaction mix was then added to a 24-well nanoplate. Reactions were then cycled according to the manufacturer’s protocol (40 mins at 50 °C, 2 mins at 95 °C, and 40 cycles of 5 secs at 95 °C followed by 30 secs at 60 °C).

Nucleic acid extracts for BAV-3 and BHV-1 were quantified using the QIAcuity in either an 8.5k partition 24-well (Qiagen 250011), 8.5k partition 96-well (Qiagen 250021), or a 26k 24-well partition plate (Qiagen 250001) based upon availability. In the case of 8.5k partition plates, 20 μL reactions consisted of 5 μL of 4x Probe PCR Master Mix, 2 μL of 10x primer-probed mix, 8 μL of nuclease-free water, and 5 μL of a 1:1,000 dilution of the desired nucleic acid extract. 15 μL of the resulting reaction mixture was added to the 8.5k partition plate. In the case of 26k partition plates, 44 μL reactions consisted of 11 μL of 4x Probe PCR Master Mix, 4.4 μL of 10x primer-probed mix, 17.6 μL of nuclease-free water, and 11 μL of a 1:1,000 dilution of the desired nucleic acid extract. 42 μL of the resulting reaction mixture was added to the 26k partition plate. Plates were then heated for 2 mins at 95 °C followed by 40 cycles consisting of 15 secs at 95 °C and 30 secs at 60 °C.

Following thermal cycling, endpoint fluorescent intensity measurements were read on the Green fluorescent channel with an exposure time of 500 ms and a gain of 6.

### 2.6. Quantification of unextracted viruses via digital polymerase chain reaction

Unextracted or whole viruses from cell culture supernatants were quantified using digital PCR using the same reaction formulation presented in section 4.5 with the exception that the desired nucleic acid was replaced with a variable dilution of virus culture supernatant. Virus culture supernatants were diluted 1:100; 1:1,000; and 1:10,000 before being added to the dPCR reaction formulation. No template control (NTC) reactions were conducted by adding 11 µL in place of diluted virus culture supernatant. All reactions were conducted in triplicate.

Virus concentration was quantified using the QIAcuity software suite. Fluorescent thresholds for determining positive and negative partitions were set uniformly for all dilution levels of a given virus by manually determining the value such that all partitions in the NTC reactions were counted as negative. The threshold was then set at a whole integer such that the set threshold less the maximum RFU of all partitions in the NTC reactions was < 2 RFU. If the threshold was greater than 10 RFU from the mean RFU across all partitions in the NTC reactions, it was determined that at least one false positive had occurred, and the quantification was re-conducted for that virus. Virus concentrations for each replicate and mean concentrations for the three replicates were calculated from negative and positive partition counts and reported according to the software suite.

The final virus concentration in the solution was determined by selecting the dilution level at which all replicates had a negative partition percentage (defined as the number of negative partitions divided by the number of total valid partitions multiplied by 100) between 5 and 95%. The virus concentration in the supernatant was then calculated from the mean concentration as reported by the software to account for the 1:4 dilution when adding to the dPCR reaction mix and then the selected dilution level when diluting from the neat virus culture supernatant. If multiple dilution levels had negative partition percentages between 5 and 95%, the dilution level that had the lowest 95% confidence interval around the mean concentration as reported by the QIAcuity software suite was selected.

For dPCR quantification in parallel with qLAMP reactions, an in-house python program was used to analyze raw data produced by the QIAcuity software suite. This software performed the same computations at the QIAcuity software suite but allowed for more nuanced control over the calculation of confidence intervals and hyperwells. Details on the programs’ availability can be found in “Data Availability”.

### 2.7. Determination of optimal reaction final sample concentration

Field samples were used neat, diluted 1:1 in nuclease-free water, or diluted 1:10 in nuclease-free water to result in sample concentrations of 100%, 50%, or 10%, respectively. BVDV-1 virus culture supernatant was diluted in nuclease-free water 10-fold four times resulting in a 1:10 to a 1:10,000 virus culture supernatant dilution. The last three dilutions (1:100, 1:1000, and 1:10,000) were then added to the field sample dilutions at a ratio of 1:10 resulting in a final sample concentration of 90%, 45%, and 9% at a BVDV-1 virus concentration of 1:100; 1:1,000; or 1:10,000. 5 μL of the resulting field sample and virus dilutions was then added to 15 μL QIAcuity OneStep Advanced RT-dPCR master mix. 15 μL of the resulting mixture was then added to a 96-well, 8.5k QIAcuity dPCR plate and sealed according to manufacturer protocol. The plate was then heated according to the same thermal cycling profile as for quantifying nucleic acids above.

### 2.8. Loop-mediated isothermal amplification primer design and nomenclature

LAMP primer sets were designed using Primer Explorer v5 (https://primerexplorer.jp/lampv5e/index.html). Primer sets were designed to target genes unique to a specific virus when compared to the other four viruses we investigated in this study. Approximately 3 – 5 primer sets were designed per gene targeted. Specifically, we designed primers to target the bICP0, bICP22, gC, gD, US8, and UL23 genes of BHV-1; the E1a, E1b, E3, Hexon, Penton, and pV genes of BAV-3; the M2, NS1, and NS2 genes of BRSV; the F, M, and L genes of BPIV-3; and the 5’ untranslated region (5UTR) and the E2 and NS3 genes of BVDV-1. GenBank accession numbers and nucleotide indices for sequences used for LAMP primer design are provided in Table S2. Additionally, sequences in FASTA format can be found in the dataset accompanying this publication (see “Data Availability”).

We named primer sets used in this study by concatenating the virus abbreviation, the gene target, and a unique primer ID, each separated by a period. The unique primer ID began with “1” and we incremented it by one for each newly designed primer set targeting a given region. We refer to primer sets as “candidate primer sets” before and throughout screening until we select them as the final primer set for characterizing primer set performance in field samples, at which point we simply refer to them as the “primer set”.

### 2.9. Primer screening and selection

Designed primer sets were subjected to a regime of LAMP reactions to screen for optimal performance of a variety of LAMP reaction performance metrics. LAMP primer sets were first screened to select for a lack of non-specific amplification due to primer dimerization in NTC reactions and fast reaction time. This was accomplished by executing LAMP reactions with four technical replicates using viral culture supernatant genomic extracts in positive reactions and nuclease-free water in NTC reactions. Genomic extracts were diluted to a concentration of 2.0 × 10^4^ viral genomic copies per μL and five μL were added per LAMP reaction (i.e., 10^5^ copies/reaction). LAMP primer sets exhibiting amplification in the NTC reactions before 30 mins were not considered for further screening and selection. Primer sets were scored according to a modified algorithm previously developed by the Verma Lab (Pascual-Garrigos et al. 2021). Modifications to the algorithm are detailed in Supplemental Information, Section 1.

Primer sets passing initial screening were then subjected to a series of assays to determine the LOD. The LOD was determined by first running a 10-fold dilution series from 1.0 × 10^5^ copies per reaction to 1.0 × 10^0^ copies per reaction which is referred to as a “coarse LOD”. Each concentration was conducted in triplicate and NTC reactions were included in triplicate for each primer set. The coarse LOD was determined as the lowest concentration in which all technical replicates—and all technical replicates at higher concentrations—amplified before 30 mins. Once a coarse LOD was determined, a series of six, two-fold dilutions were performed in triplicate with associated NTC reactions per primer set. The resulting LOD is referred to as the preliminary LOD. For all LOD experiments, at least one replicate below the LOD and not in the NTC reactions must not have amplified before 30 mins for the LOD to be valid.

### 2.10. Loop-mediated isothermal amplification reaction and formulation

A 2x primer mix was made consisting of 3.2 μM of both FIP and BIP primers, 0.8 μM of both LF and LB primers, and 0.4 μM of both F3 and B3 primers. The 2x primer mix was heated at 90 °C for 10 mins in a dry bath and was allowed to cool at room temperature for 5 mins. This heating cycle was repeated weekly to minimize the formation of primer dimers in the primer mix.

LAMP reactions consisted of 12.5 uL of 2x WarmStart Mastermix (DNA/RNA) with UDG and UTP spiked-in, 1x LAMP fluorescent dye (New England Biolabs (NEB) B1700; Ipswich, MA, USA), 2.5 µL of 2x primer mix, and 5 uL of template nucleic acid at the indicated concentration. 2x WarmStart Mastermix with UDG/dUTP was prepared by adding to 3.5 µL of 100 µM dUTP (Fisher Scientific R0133) and 5.0 uL of Antarctic Thermolabile UDG (NEB M0372) diluted 1:10 in nuclease free water to a single 1.25 mL tube of 2x WarmStart Mastermix (NEB M1800) which contains WarmStart Reverse Transcriptase in its formulation. NTC reactions used nuclease-free water in place of template nucleic acid. LAMP reactions were run using three or four technical replicates as indicated. Reactions were run on a qTower 3G (Analytik Jena 844-00554-4) or qTower 3G Touch (Analytik Jena 844-00506-2) for 60 mins at 65 °C (1 °C/sec ramp rate). Every 60 secs, fluorometric readings were taken on the Green Channel with a gain of 3 (qTower 3G Touch) or 2 (qTower 3G). Following 60 mins of heating, enzymes were inactivated at 85 °C for 5 mins (8.0 °C/sec ramp rate).

### 2.11. Cross-reactivity analysis

Cross-reactivity of selected primer sets with the other viruses examined in this study was determined by conducting 24 different LAMP reactions with the indicated primer set and varying the template. 5 µL of nucleic acid extract for each virus was added to 20 µL of LAMP reaction to result in a final concentration of target nucleic acid extract of 1.0 × 10^4^ copies per reaction. Each primer set was challenged in triplicate with each virus individually.

### 2.12. Field sample collection and processing

Field samples were collected from 24 cattle that appeared apparently healthy according to the DART method (Juge et al. 2022; Kamel et al. 2024). Nasal swab collection was carried out according to the swab collection methodology previously reported in our fieldwork (Centeno-Martinez et al. 2022) Briefly, a lint-free Kimwipe® was used to clear debris (dirt, excess mucous, saliva, etc.) from the external surface of the nasal cavity. Samples were then collected by inserting a double-tipped dry swab into one nasal cavity and briefly inserted into the nasal cavity of the animal and the perimeter was swabbed. The same was done to the opposite nasal cavity using the same swab. The swab was then placed on ice and transported to a laboratory setting and was not placed in any buffer. Identification numbers of cattle were recorded for each sample. Sample collection was performed in accordance with Purdue Institutional Animal Care and Use Committee (IACUC) protocol number 1906001911. Once collected, field samples were refrigerated dry at 4 °C until processing as described below. Samples were processed within 72 hours of collection.

Samples were processed using a modified procedure to collect the viral content from the supernatant. Briefly, 700 μL of nuclease-free water was added to a clean 1.5 mL Eppendorf microcentrifuge tube. Excess mucous on the double-tipped swabs was removed by gently using the inside wall of the collection tube to scrape the excess mucous from the tip. Each tip of the double-tipped swab was then placed into separate 1.5 mL Eppendorf tubes such that two tubes were used for a single double-tipped swab. The swab shafts were then trimmed using a pair of scissors and placed into a 1.5 mL microcentrifuge tube. Care was taken to ensure that the scissors were clean before cutting each sample to prevent cross-sample contamination. Samples were then vortexed for 5 mins to resuspend the contents of the swab. Swabs were then removed from the tubes and decontaminated in bleach before disposal. The resuspension was then centrifuged at 5,000 x g at 20 °C for 10 mins. The supernatant from both tubes corresponding to one sample was then pooled in a sterile microcentrifuge tube and kept at -20 °C until use. Prior to usage, samples were thawed and centrifuged at 5,000 RCF at 20 °C for 5 mins to ensure excess mucosal content was not introduced in the LAMP reaction.

### 2.13. Analytical sensitivity and analytical specificity analysis

Analytical sensitivity and analytical specificity analyses were conducted using specifications and guidance set forth by the United States Food and Drug Administration (FDA) for human diagnostic development (Hess et al. 2012; U.S. Food & Drug Administration 2007; U.S. Food & Drug Administration 2024). 30 replicates comprised of 15 samples conducted in duplicate were used for the sensitivity and specificity analyses in order to capture variance between samples. The top 15 samples when sorted first by concentration of the target virus and then alphabetically were selected for the analysis of each virus. All samples contained background viral concentrations less than the preliminary LOD for the tested primer set except for BAV-3.pV.3 which required some samples between 1x and 2x the preliminary LOD.

Whole virus from cell culture supernatant was spiked into selected field samples at a concentration equivalent to the preliminary LOD (1x LOD) and at a concentration of twice the preliminary LOD (2x LOD). Additionally, nuclease-free water was used in place of whole virus for NTC reactions. dPCR was used to verify the concentration of the spiked virus in both the 1x LOD and 2x LOD concentrations. Only one sample was tested and used as the representative concentration of all samples since the dPCR was not able to quantify more than 2 samples at this relatively low concentration due to throughput limitations.

ROC curves were constructed by varying the reaction time at which a given reaction was determined to be “positive” or “negative.” The optimum cutoff point was determined to be the reaction time which minimized the distance from (0,1) on the ROC curve, which formally is the point which minimized the error, 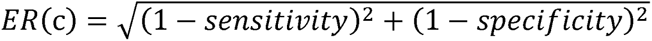 (Unal 2017). The sensitivity at this optimum cutoff point was then used to represent the sensitivity of the primer set at the respective concentration. This analysis was repeated for 1x and 2x LOD and raw data is available as outlined in Data Availability. If a primer set did not achieve a sensitivity of 95% (19 out of 20 samples amplify) at 1x LOD, then the results from 2x LOD were used to calculate all further metrics. Establishment of the final LOD was determined as the concentration of either 1x LOD or 2x LOD which had a sensitivity exceeding or closest to 95%.

### 2.14. *In silico* inclusivity analysis

Inclusivity analyses were conducted using a BLAST search of the target sequence (Table S2) with which primers were designed (Supplementary File 2) with all sequences available for the target virus in NCBI as categorized by the taxon ID for the respective virus. Taxon IDs used in this analysis are as follows: BAV-3=129950, BHV-1 = 10320, bPIV-3 = 3052729, BRSV = 3050248, and BVDV-1 = 11099. Default *blastn* values were used except the argument for maximum returned alignments was set at the largest value. All returned BLAST alignments with a query cover greater than or equal to 50% were downloaded and compiled into a single FASTA file. Primer sets (with inner primer separated into constituent primer parts) were additionally compiled into a separate FASTA file.

Primers were then aligned to returned BLAST matches using the “msa.sh” file of BBMap v39.33 (Bushnell 2014), which returns the highest homology match of each primer against each returned BLAST sequence. The results were filtered to ensure inclusivity only considered alignments of primer sets with their intended target (i.e. only alignments of BHV-1 primer sets with BHV-1 sequences were considered). The CIGAR string which is string representation of the alignment was used to calculate the number of mismatches within an alignment. The number of mismatches across all primers in a primer set for a given target sequence were summed together. The portion of primer set alignments containing *n* mismatches from a given number of alignment sequences for a target was then calculated, where *n* was allowed to vary from 0 (for no mismatches in any primer) to 20+. We then calculated the cumulative inclusivity for the number of primer sets containing 10 or fewer mismatches and reported this percentage as the primer set inclusivity.

### 2.15. Data Availability

Raw data used to support the analyses and results of this study is located at https://doi.org/10.17632/xfv4nyrj7t.1. Python files, documentation, and scripts used during the analysis of the data collected can be found at https://github.itap.purdue.edu/VermaLab/ReactionAnalysisLibrary (for code and methodology related to digital assay quantification and inclusivity) and https://github.itap.purdue.edu/VermaLab/PrimerScoring (for cases of primer scoring, LOD, and sensitivity/specificity analysis).

Supplementary Data can be accessed according to the online version of this article. Supplementary Data and Information contains the following: Supplementary File 1 contains all supplementary figures, supplementary tables, and additional discussions pertaining to analysis decisions. Supplementary File 2 contains nucleotide sequences of all LAMP primers used in this study, while Table S1 displays dPCR primers. Supplementary File 3 contains a copy of the GitHub repositories, including code and documentation, at the time of article submission. Please note that updated code may be available at the Github repositories above.

## 3. Results

This study highlights two novel improvements: the demonstration of LAMP and RT-LAMP assays targeting BAV-3, BHV-1, bPIV-3, BRSV, and BVDV-1 in unextracted, minimally processed field samples; and the development of a pipeline for designing, screening, and selecting LAMP/RT-LAMP assays. Figure 1 shows a **H** process for our field-based LAMP assays compared to traditional diagnostics. Since our workflow does not require transportation to a laboratory setting or an extraction step requiring costly equipment and specialized training, sample-to-answer time for our assay (1.8 hrs) is almost half of that required for traditional diagnostics (3.5 hrs). Furthermore, the centrifugation step could be replaced with a filtration step upon proper validation to further decrease the sample-to-answer time.

To showcase the utility of our assays in complex sample, we will present the data in reverse chronological order in which it was collected. Accordingly, we will first present results for performance in field samples which will be followed by a detailed discussion of primer design and screening.

### 3.1. Comparison of performance in complex sample versus extracted nucleic acids

After selecting the primer sets for the five pathogens, we evaluated their performance using viral nucleic extracts and using cultured whole virus spiked into field samples (containing a final sample concentration of 1.8% v/v after dilution in nuclease-free water and spiking in the whole virus). (Figure S1 – S2; Table S3 – S4) Table 2 shows the reported preliminary LOD with dPCR confirmation of each selected primer set targeting each virus in both nuclease-free water and complex sample. Ratios of the preliminary LOD when we used whole virus in diluted field sample to extracted nucleic acids in water ranged from 0.5 (in the case of BRSV) to 16 (in the case of BHV-1). Except for BHV-1, all ratios were less than or equal to 10.

**Table 2:** Tested, Observed, and preliminary LODs of the qLAMP/RT-qLAMP reactions using either nucleic acid extract in nuclease-free water or whole virus from cell culture supernatant spiked into complex sample (final reaction concentration of 1.8%). The ratio of preliminary LODs in the two environments is reported to show the change in LOD when inhibitors from complex sample are present in the reaction.

| Primer Set | Nucleic Acid Extract LOD (copies/reaction) |  |  | Whole Virus in Complex Sample LOD (copies/reaction) |  |  | Ratio of Preliminary LOD (field sample / extracted nucleic acids) |
| --- | --- | --- | --- | --- | --- | --- | --- |
|  | Expected LOD | Observed LOD [(95% confidence Interval)] | Preliminary LOD | Expected LOD | Observed LOD [(95% confidence Interval)] | Preliminary LOD |  |
| **BAV-3.pV.3** | 12.5 | 10 (10, 12) | 12.5 | 50 | 27 (24, 31) | 50 | 4 |
| **BHV-1.gD.4** | 6.25 | 4 (3, 6) | 6.25 | 50 | 102 (77, 129) | 100 | 16 |
| **bPIV-3.L.1** | 25 | 42 (34, 52) | 50 | 500 | 230 (-54, 515) | 500 | 10 |
| **BRSV.M2.1** | 500 | 815 (789, 843) | 1,000 | 500 | 456 (349, 565) | 500 | 0.5 |
| **BVDV-1.5UTR.3** | 250 | 235 (194, 276) | 250 | 500 | 472 (421, 523) | 500 | 2 |

### 3.2. Determination of optimal sample dilution for reverse-transcription loop-mediated isothermal amplification assay

Figure 2A shows the viral quantification of BVDV-1 at various dilution levels of viral culture supernatant diluted in either neat samples (final reaction concentration: 18%); 50% sample diluted in nuclease-free water (final reaction concentration: 9%); 10% sample diluted in nuclease-free water (final reaction concentration: 1.8%); or nuclease-free water with no sample. (Table S5, Figure S3) We can see across all viral dilution levels that the quantification of the 0% field sample concentration (“NTC”) and the quantification of the 1.8% field sample enclose the confidence bounds of one another, indicating that no significant variation exists in the quantification of those two levels. The same does not hold when comparing the 0% and 9% field sample concentrations at viral dilution levels of 1:100 and 1:1,000, nor for any viral dilution level in the 18% field sample (final reaction concentration). Therefore, we determine that the optimum “tolerable” field sample concentration that will not significantly affect dPCR quantification or qLAMP/RT-qLAMP performance is 1.8% (final reaction concentration). We thus use a final reaction concentration of 1.8% for all further studies containing field samples unless otherwise noted.

**Figure 2:**
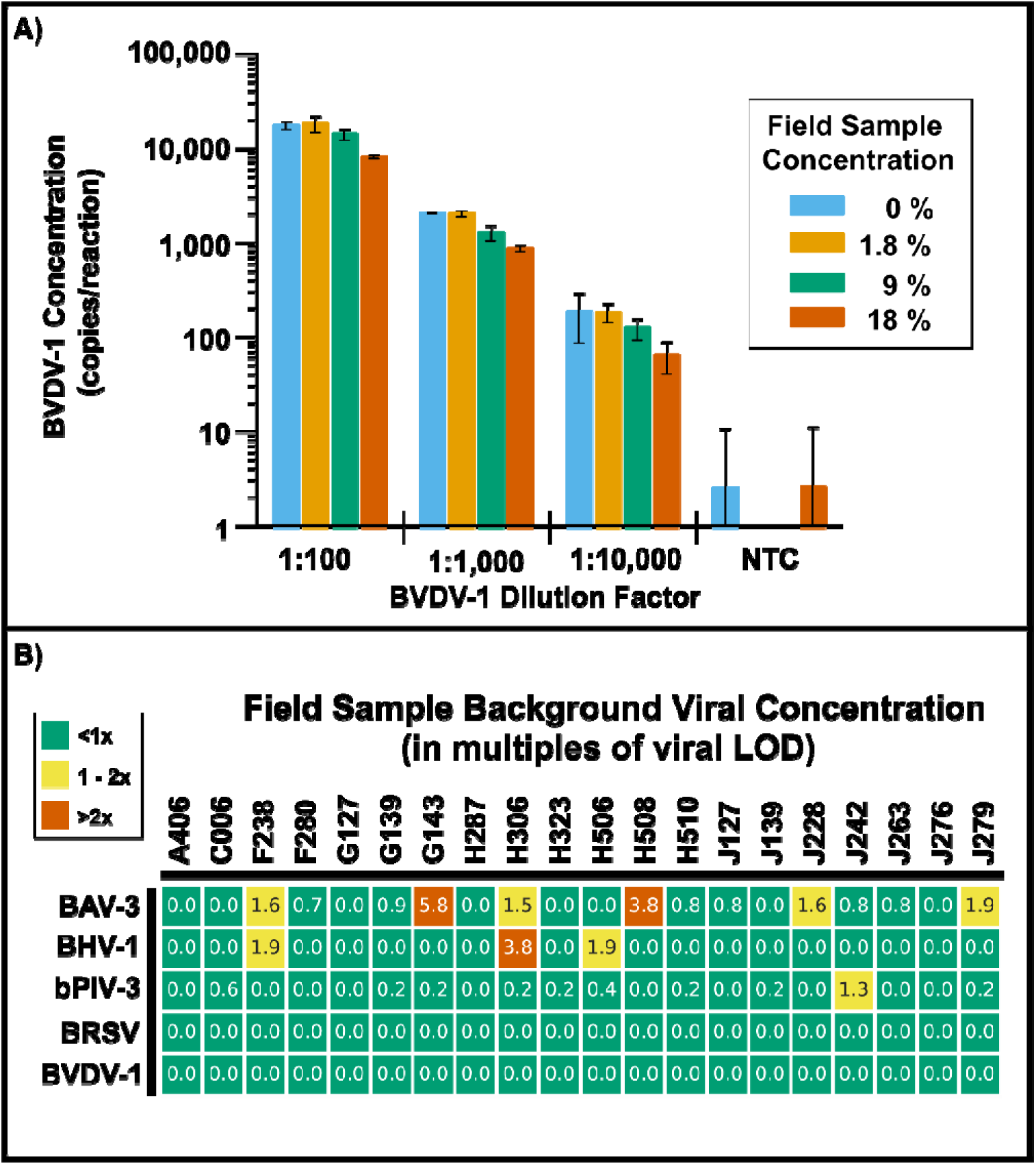
**A)** Effect of field sample concentration on dPCR quantification consistency of BVDV-1 virus culture supernatant using dPCR primer set BVDV-n. The color indicates the final field sample concentration in the dPCR reaction and the virus culture supernatant concentration before being added to the dPCR reaction is provided along the horizontal axis. **B)** Concentration of viruses in this investigation natively in the background of collected field samples. Concentrations as measured using dPCR of diluted field samples (9% final sample concentration) are reported in a multiple of LOD, which varies by virus. dPCR primer sets used for quantification of viruses are: BAV-3: BAV-3.Kishimoto.Hexon; BHV-1: BHV-1.Chandranaik.gB; bPIV-3: BPI3ii.Thonur; BRSV: BRSV.Boxus.N; BVDV-1: BVDV-n. Cells highlighted in vermillion indicate that the background concentration meets or exceeds twice the concentration at the for the selected primer set targeting that virus and is therefore unfit for usage in further testing. Cells highlighted in yellow indicate the background viral concentration is within 1x of the LOD for the selected primer set and therefore should be avoided for further usage in testing and sensitivity and specificity studies. Cells highlighted in green are the field samples selected for the characterization of selected LAMP primer sets in complex samples.

Since we spiked target viruses in complex samples to characterize the performance of our designed assays, we must ensure that our samples do not contain appreciable amounts of our target virus. To this end, we quantified the viral concentration of all target viruses in the sample background. (Table S6; Figure S4 – S5) We set the final sample concentration in the dPCR reactions to 2.25%, matching the template dilution factor (effectively 1:4 after adding the template to the dPCR reaction) used in other quantifications in this study. We quantified the target virus in the background of 20 field samples for each virus and prioritized which samples to use in downstream characterization based on the measured concentrations. We expressed background viral concentrations as a multiple of the primer set’s LOD targeting the respective virus. (Figure 2B) We excluded samples with a concentration greater than 2x the LOD from further characterization. Moreover, we only used samples with background viral concentrations greater than 1x but less than 2x of the LOD if additional sample volume was necessary (which was only the case for BAV-3). We specifically note that four samples tested negative for any of our target viruses (less than 10% of the respective LOD). We used these samples for downstream characterization of primer sets in all cases except for sensitivity and specificity analysis, which required 15 samples tested in duplicate.

Please see Supplementary File 1 for additional discussion on the selection of the optimal field sample dilution ratio.

### 3.3. Sensitivity and specificity tests for selected primer sets

Once we selected a final primer set for our assays and determined a limit of the detection, we proceeded to determine the analytical sensitivity, specificity, and accuracy of the assay in complex field sample. Figure 3 and Table S7 show the results of these analyses for each target virus. Response times for DNA and RNA viruses showed no significant difference, ranging between 29 mins (for BVDV-1.5UTR.1) and 45 mins (for BAV-3.pV.3).

**Figure 3:**
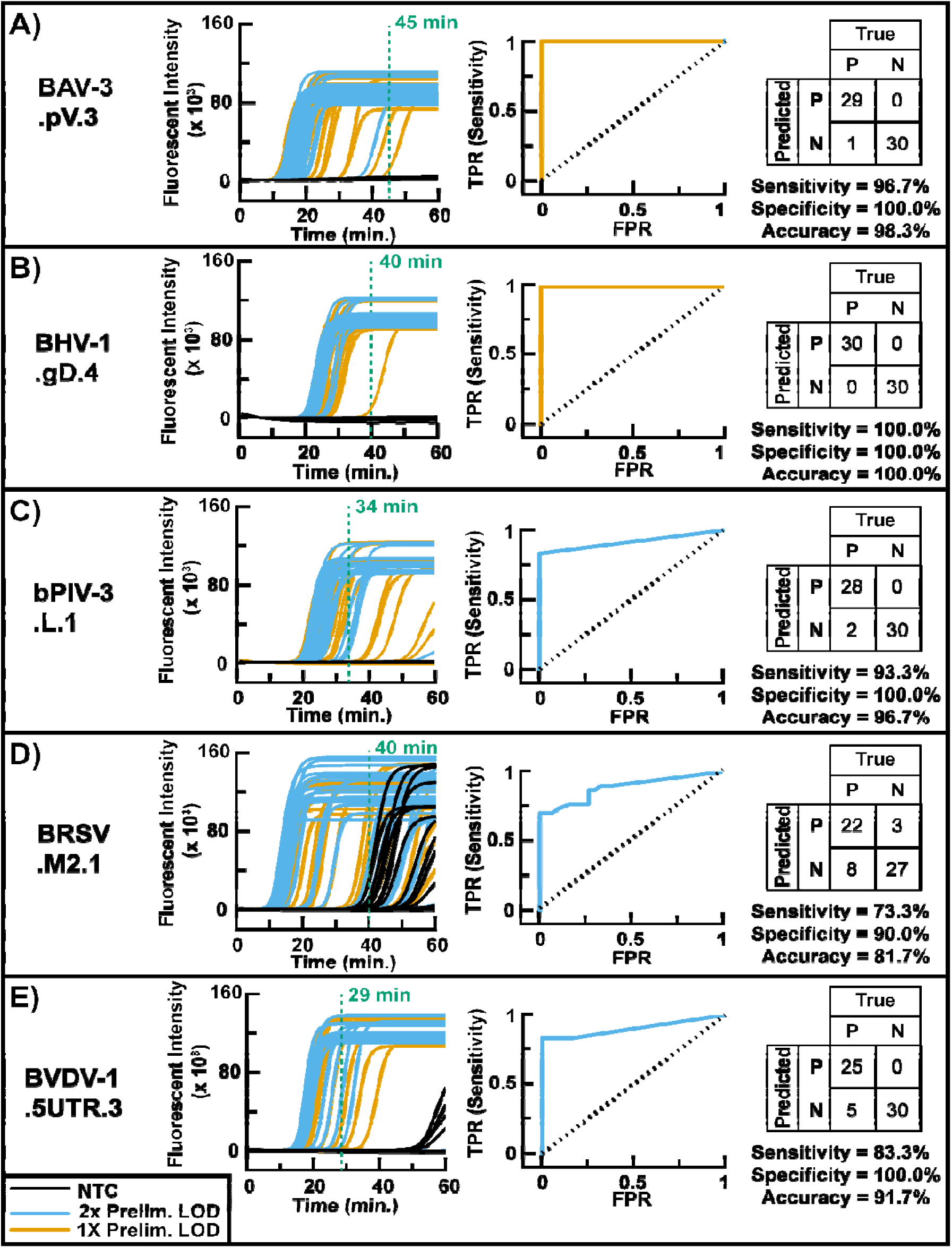
Sensitivity and specificity analysis with accompanying receiver operator characteristic (ROC) curve for **A)** BAV-3.pV.3 **B)** BHV-1.gD.4, **C)** bPIV-3.L.1, **D)** BRSV.M2.1, and **E)** BVDV-1.5UTR.3. The leftmost chart for each primer set is the fluorescent trace of the qLAMP/RT-qLAMP reactions of 30 replicates (15 field samples ran in duplicates) using 5 μL of whole virus from cell culture supernatant spiked into 9% field sample (final reaction concentration of 1.8%) to result in a final concentration equivalent to twice the limit of detection (LOD; blue line), the limit of detection (orange line), or nuclease-free water in place of whole virus (black line). Optimum reaction threshold times which minimize error are indicated in green and annotated with the respective threshold time. The chart to the right is an ROC curve constructed by varying the reaction threshold time to determine positive reactions using the lower of either 2x or 1x LOD such that the sensitive at the optimum reaction time either exceeds 95% or is maximized. Finally the chart to the left indicates the counts (from top left to bottom right quadrant) for true positives, false positives, false negatives, and true negatives when using the optimum reaction threshold times. Using these counts, the sensitivity, specificity, and accuracy are reported

Table 3 displays the results of the analytical sensitivity analysis at both 1x and 2x LOD for each primer set. As we expect given the relative stability of DNA versus RNA, both DNA viruses had a higher accuracy (approaching 100%) while the RNA viruses had accuracies ranging from 81.7% to 96.7%. Analytical specificities for all viruses except for BRSV were 100%. Both BAV-3.pV.3 and BHV-1.gD.4 attained sensitivities of 95% at 1x LOD. Accounting for the processing of the sample to the swab, dilution of complex sample to 1.8% (final reaction concentration), and addition of 5 μL of template to the reaction, the LOD expressed as the viral concentration per swab is on the order of 2 × 10^5^ copies per swab for the DNA viruses.

**Table 3:**
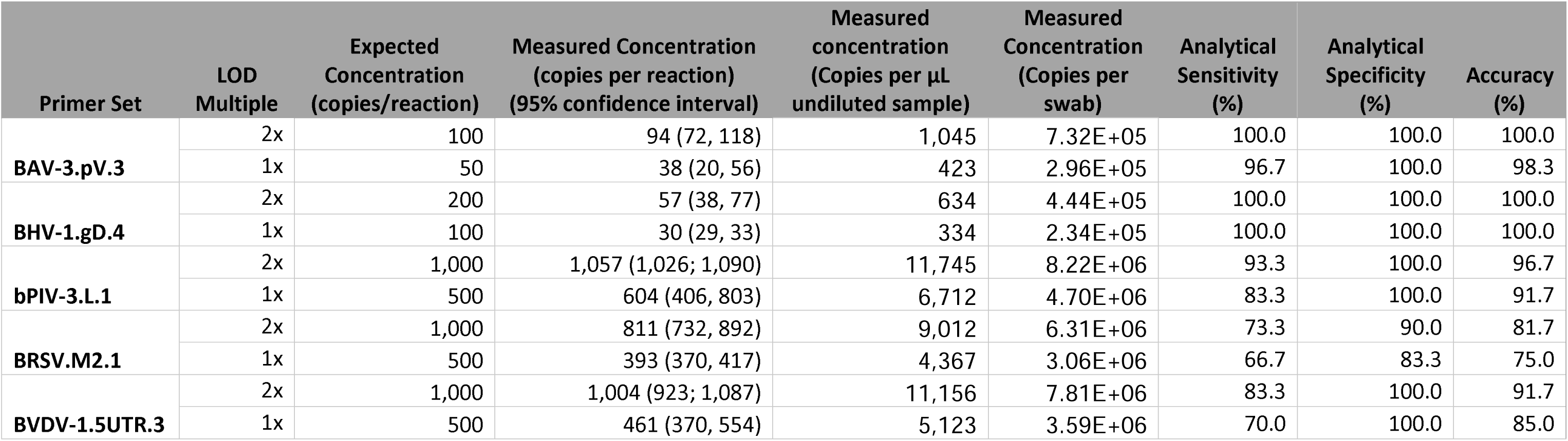
Expected concentration (in genomic copies per reaction) and digital PCR (dPCR) parallel quantification measurements for whole virus from cell culture supernatant spiked into complex field samples (1.8% final reaction concentration) for sensitivity and specificity analyses conducted at the preliminary LOD and a concentration of twice the preliminary LOD

| Primer Set | LOD Multiple | Expected Concentration (copies/reaction) | Measured Concentration (copies per reaction) (95% confidence interval) | Measured concentration (Copies per μL undiluted sample) | Measured Concentration (Copies per swab) | Analytical Sensitivity (%) | Analytical Specificity (%) | Accuracy (%) |
| --- | --- | --- | --- | --- | --- | --- | --- | --- |
| **BAV-3.pV.3** | 2x | 100 | 94 (72, 118) | 1,045 | 7.32E+05 | 100.0 | 100.0 | 100.0 |
|  | 1x | 50 | 38 (20, 56) | 423 | 2.96E+05 | 96.7 | 100.0 | 98.3 |
| **BHV-1.gD.4** | 2x | 200 | 57 (38, 77) | 634 | 4.44E+05 | 100.0 | 100.0 | 100.0 |
|  | 1x | 100 | 30 (29, 33) | 334 | 2.34E+05 | 100.0 | 100.0 | 100.0 |
| **bPIV-3.L.1** | 2x | 1,000 | 1,057 (1,026; 1,090) | 11,745 | 8.22E+06 | 93.3 | 100.0 | 96.7 |
|  | 1x | 500 | 604 (406, 803) | 6,712 | 4.70E+06 | 83.3 | 100.0 | 91.7 |
| **BRSV.M2.1** | 2x | 1,000 | 811 (732, 892) | 9,012 | 6.31E+06 | 73.3 | 90.0 | 81.7 |
|  | 1x | 500 | 393 (370, 417) | 4,367 | 3.06E+06 | 66.7 | 83.3 | 75.0 |
| **BVDV-1.5UTR.3** | 2x | 1,000 | 1,004 (923; 1,087) | 11,156 | 7.81E+06 | 83.3 | 100.0 | 91.7 |
|  | 1x | 500 | 461 (370, 554) | 5,123 | 3.59E+06 | 70.0 | 100.0 | 85.0 |

RNA viruses, on the other hand, could not achieve 95% sensitivity even at 2x LOD. Instead, the sensitivity attained was 93.3%, 73.3%, and 83.3% for primer sets targeting bPIV-3, BRSV, and BVDV-1, respectively, at 2x LOD. Established LODs for the RNA viruses ranged from 9.0 × 10^4^ to 1.1 × 10^5^ copies/μL of undiluted sample when rounded up to the nearest 100 copies which corresponds to a concentration of 6.31 × 10^6^ to 8.22 × 10^6^ copies per swab.

### 3.4. Overview of the loop-mediated isothermal amplification primer screening process in idealized conditions

Once we designed primer sets for a target gene, which we refer to as candidate primer sets, we screened them using a variety of experiments to quantify and characterize both the performance of the candidate primer sets and their specificity in detecting the intended target pathogen. To optimize this process in terms of time and resources used, we designed a primer screening process consisting of four stages – Stage I to Stage IV – to select a final primer set in idealized conditions. In each stage, primer sets were evaluated and discarded from further consideration if certain performance criteria were not met. We illustrate this process in the flowchart provided in Figure 4.

**Figure 4:**
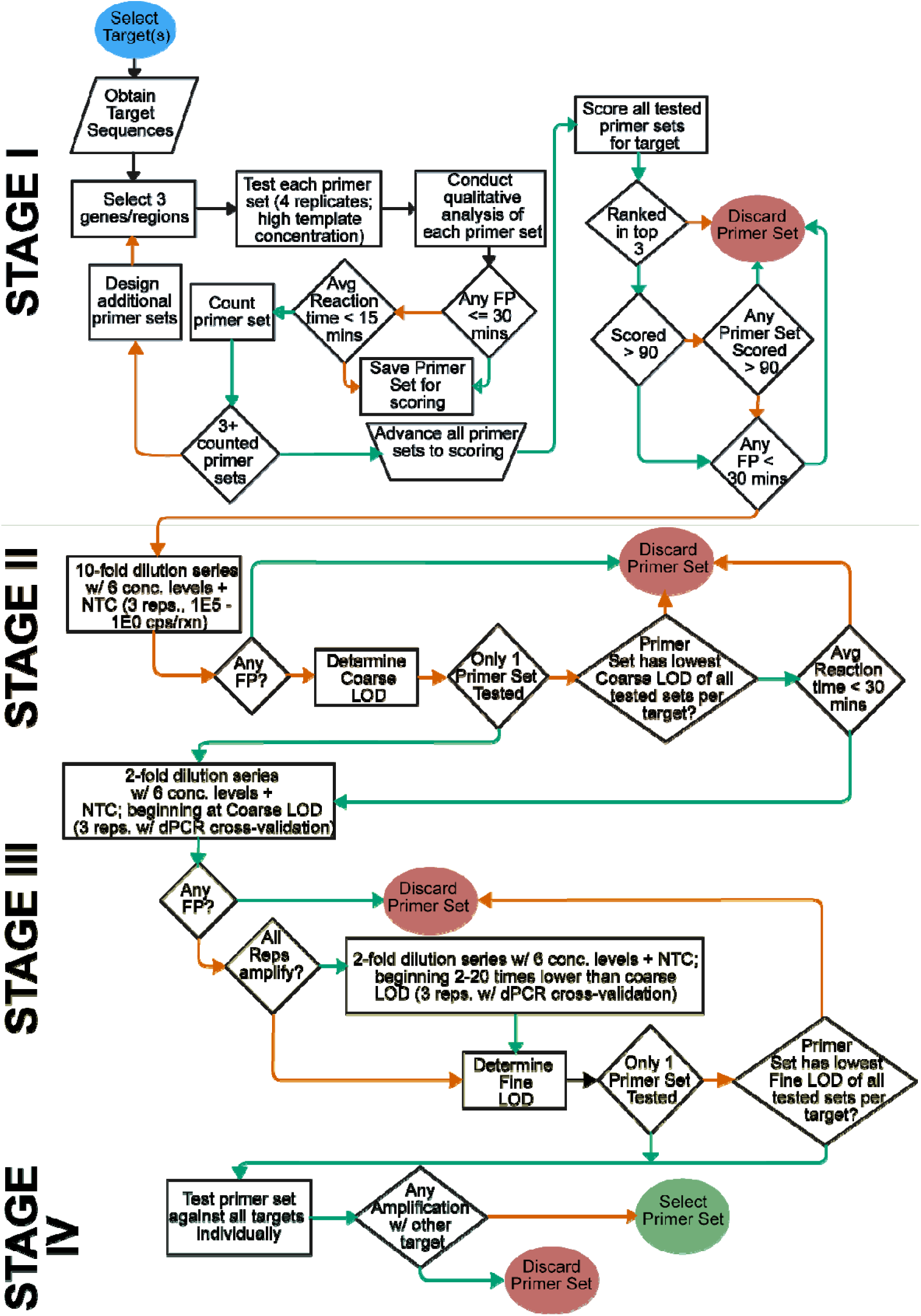
Schematic decision diagram detailing the primer design, screening, and selection processes. The diagram flows from the top left to the bottom right. Diamonds indicate a decision point that can have a “yes” or “no”. The route taken depending on that decision is indicated by a green or red arrow, respectively, flowing from the diamond. Abbreviations used: Average (“Avg.”), false positive (FP), limit of detection (LOD), with (“w/”), concentration (“conc.”), replicates (“reps”), digital PCR (dPCR).

We designed Stage I of the primer screening process to investigate the optimal performance of the primer sets in ideal conditions. We used Stage II to determine a concentration range – a coarse LOD – for primer sets which we used to focus our efforts in Stage III. We used Stage III to determine the final (“fine”) LOD. Finally, we designed Stage IV to investigate the cross-reactivity of final primer sets with other viruses used in the study.

### 3.5. Primer screening and selection

We designed a total of 86 LAMP/RT-LAMP primer sets targeting the five viruses examined in this study (Supplemental File 2). Tables S8 – S12 display the overall score and performance characteristics of the designed primer sets based on the primer screening data. (Figures S6 – Figure S13) Of the 86 primer sets we screened in Stage I, we advanced only 11 primer sets to Stage II. (Table 4)

**Table 4:** Select Stage I screening characteristics and overall scores (out of 100) of candidate primer sets advanced to Stage II screening.

| Primer Set | Maximum Intensity<br>[RFU (Standard Deviation)] | Reaction time<br>[mins (Standard Deviation)] | False Positives | Overall Score |
| --- | --- | --- | --- | --- |
| **BAV-3.pV.2** | 89,680 (± 11,932) | 9.00 (± 0.0) | 0 | 95.22 |
| **BAV-3.pV.3** | 118,937 (± 9,845) | 11.00 (± 0.7) | 1 | 92.85 |
| **BAV-3.Penton.3** | 90,266 (± 23,516) | 7.00 (± 0.0) | 0 | 92.76 |
| **BHV-1.gD.4** | 69,113 (± 3,871) | 6.50 (± 0.5) | 0 | 92.75 |
| **bPIV-3.L.3** | 108,573 (± 22,114) | 12.75 (± 0.4) | 0 | 86.52 |
| **bPIV-3.L.1** | 104,277 (± 23,480) | 9.50 (± 0.5) | 0 | 86.49 |
| **bPIV-3.M.3** | 101,690 (± 8,426) | 21.75 (± 0.8) | 0 | 84.96 |
| **BRSV.NS2.3** | 120,363 (± 9,134) | 8.00 (± 0.0) | 0 | 98.41 |
| **BRSV.M2.1** | 88,979 (± 7,799) | 7.00 (± 0.0) | 0 | 96.04 |
| **BVDV-1.E2.2** | 126,762 (± 9,135) | 11.50 (± 0.5) | 0 | 94.06 |
| **BVDV-1.5UTR.3** | 119,312 (± 9,599) | 9.50 (± 0.5) | 1 | 91.55 |

We determined the Coarse LOD in Stage II to be 100 copies/reaction for DNA viruses (BAV-3 and BHV-1; Figure S14) and 1,000 copies/reaction for RNA viruses (bPIV-3, BRSV, and BVDV-1). (Table 2 and Table S13; Figures S15 – S16) We advanced a total of seven candidate primer sets to Stage III, two targeting each of BAV-3 and BRSV, and then one primer set for each of BHV-1, bPIV-3, and BVDV-1. (Figures S17 – S18) Of the seven primer sets advanced to Stage III, the best LOD was selected from the viruses containing multiple primer sets were selected as the final primer set to advance to Stage IV. (Table 2 and Table S14 – S15)

We chose five primer sets – one for each virus – to advance to Stage IV, the final stage of primer screening. Stage IV screening showed no significant cross-reactivity with other viruses, thus confirming that each primer set is specific to its intended virus. (Figure S19) Accordingly, we selected each of these primer sets as the final primer for field sample studies and reported their sequences in Table 5.

**Table 5:**
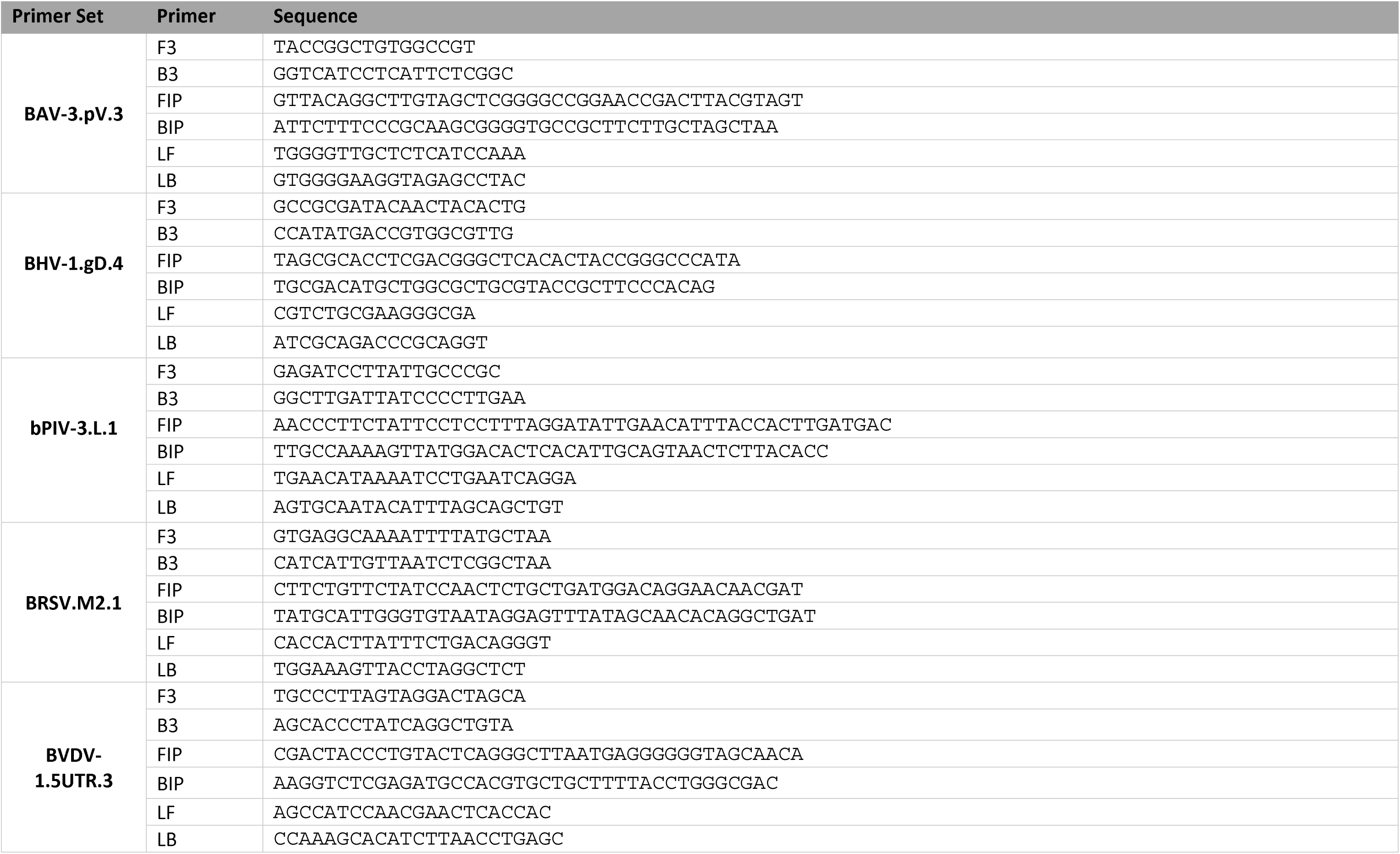
Sequences for final selected primer sets used during this study.

Finally, *in silico* inclusivity analyses were conducted for all primer sets. Table S16 shows the number of target viral sequences for each inclusivity calculation and cumulative inclusivity for primer set alignments with 10 or less mismatches across the entire set. All selected primer sets had the highest *in silico* inclusivity of all primer sets designed for the respective viral targets, except for BRSV.M2.1, which had a lower cumulative inclusivity than all other BRSV primer sets except BRSV.M2 and BRSV.M3.

## 4. Discussion

### 4.1. Comparison of performance in complex sample versus extracted nucleic acids

We previously reported that LAMP assays typically have an LOD five to ten times worse in complex matrices than in nuclease-free water, likely due to inhibitors (Davidson et al. 2021). Consistent with these observations, LODs of all primer sets except BHV-1 worsened by a factor of 10 or less. The LOD for BAV-3.pV.3 and BHV-1.gD.4 (both enveloped DNA viruses) worsened by factors of four and 16, respectively. Primer sets targeting RNA viruses, however, showed less worsening overall compared to DNA viruses but their absolute LODs in complex samples were about ten times higher (500 versus 100 copies per reaction).

The generalized “leveling” effect of the LOD for all primer sets targeting RNA viruses to 500 copies per reaction is likely caused by some general reaction mechanism rather than inherent properties of the primer sets. We typically assume inhibitors in complex sample reduce detection of RNA to a greater extent than DNA for two reasons: 1) RNA is generally more susceptible to environmental degradation than DNA, and 2) RT-qLAMP for RNA targets requires two enzymatic reactions which can be inhibited – one for reverse-transcription to cDNA and another for LAMP amplification. Since we determined the LODs in complex samples using whole virus as opposed to bare nucleic acids, viral capsids may have provided protection for nucleic acids thus reducing environmental degradation. Confirming inhibition of both enzymatic reactions and capsid protection during RT-qLAMP would require controlled studies on RNA virus stability and reverse-transcriptase efficiency in the presence of inhibitors but is beyond the scope of this study.

### 4.2. Sensitivity and Specificity tests for selected primer sets

We spiked whole virus into healthy nasal swabs at 1× and 2× the preliminary LOD for each viral target to maximize the chance of achieving 95% or greater sensitivity. Using swabs from healthy cattle provides a sampling of background microflora enabling us to confirm our primer sets are not cross reactive with commensal microbes provided NTCs reactions do not amplify. We did not adjust spike-in concentrations since tested viruses were below 1× LOD. Following determination of 1× and 2× LOD, we varied cutoff times for positive and negative amplification to construct a receiver operator characteristic (ROC) curve and determine analytical sensitivity, specificity, and accuracy.

Generally, DNA viruses displayed higher accuracy than RNA viruses. We expect this since we conducted assays in complex samples, and DNA is typically more stable than RNA. Furthermore, the reverse transcriptase we used (WarmStart RTx included in the LAMP master mix) is active from 45–65 °C (manufacturer provided). Reduced enzymatic activity near the limits of this range likely contributes to the lower accuracy of RNA virus primer sets. Furthermore, preheating of swabs before adding to LAMP reaction could degrade viral capsid and worsen LoDs.

All primer sets except BRSV.M2.1 had analytical specificities of 100%. For BRSV.M2.1, NTC reactions showed notable amplification after 40 min, possibly due to interactions with background microflora or from host DNA. Because we performed early primer screening in nuclease-free water and later stages used only one complex sample, these late NTC amplifications likely reflect variance in sample backgrounds which became evident when we used multiple samples for the sensitivity and specificity analysis.

The sensitivity of primer sets targeting DNA viruses reached 95% at 1× LOD and was higher than RNA viruses. In some cases, dPCR quantification differed from expected values based on initial stock calculations, such as BHV-1.gD.4, where dPCR quantification measured nearly three times higher than initial stock dilutions. We attribute these discrepancies to errors in stock quantification, errors in dilution, or potential errors during reaction setup. Because we ran dPCR in parallel and it showed little replicate variance, we considered these errors acceptable as they did not impact the reliability of dPCR LOD quantification.

After we account for sample processing (resuspending both swabs in 1.4 mL of nuclease-free water and diluting to 1.8% before adding to the reaction), the LODs are on the order of 10 and 10 copies per swab for DNA and RNA viruses, respectively. At these LODs, these primer sets may help veterinarians and caretakers guide treatment for symptomatic cattle but are unlikely to aid early detection or surveillance. Nevertheless, since we validated their performance in complex samples, these characterized primer sets add to currently available diagnostics. Unlike currently available diagnostics, these RT-LAMP primer sets are deployable chute-side. Further development, however, is needed to improve their LOD in complex sample and diagnostic utility in field settings.

### 4.3. Primer screening and selection

Typically, LAMP assay developers must design multiple primer sets and perform extensive wet-lab testing since robust methods for sequence-based performance prediction are not yet available. However, to our knowledge, efficient methods to conduct this screening and selection process to determine optimal primer sets are not discussed in the literature. To address this gap, we formulated a screening and selection process with defined performance criteria to quickly determine and characterize primer sets viable for testing in field samples (Figure 4). We will briefly review this process, and a more detailed description of each step is provided in Supplemental File 1.

In Stage I, we establish an outer bound for each primer set’s performance under ideal conditions without inhibitors by testing in nuclease-free reactions spiked with 1.0×10 copies per reaction. In this manner, we can assess both primer efficiency and non-specific interactions within primer sets by examining the positive reaction and NTC reaction profiles, respectively. We then rank primer sets using our lab’s previously developed ranking methodology and retain rankings for backup selections in case downstream testing fails for the selected primer set.

Originally, we designed at least three primer sets to targeting at least three distinct regions of the pathogen genome and screened them in Stage I. BAV-3 and BHV-1, however, had a relatively low number of well-performing primer sets and so we conducted another round of design for these primer sets. We roughly define a well performing primer set as one showing amplification before 20 mins and lacking amplification in the NTCs.

In Stage II and Stage III, we balance the materials and time needed to efficiently determine each primer set’s LOD. Then, in Stage III, we perform a 2-fold dilution series starting at the coarse LOD’s upper bound to refine the LOD estimation and determine a preliminary LOD. Using dPCR, we quantified the concentration of the starting template concentration in Stage II and all dilution levels in Stage III to ensure accuracy and confidence in LOD determinations.

We designed Stage II to identify a coarse LOD by conducting a 10-fold dilution series to identify the decade range of the LOD, termed the “coarse LOD.” This allows us to hone in on the decade containing the true LOD while minimizing the number of experiments needed to determine the true LOD. We defined the coarse LOD as the lowest concentration at which all replicates above and at that concentration level amplified. We determined the fine LOD in Stage III in same manner but with the additional requirement that the reported final LOD is comparable to dPCR quantification ran in parallel. This added confidence to our final LOD. In some cases, we conducted additional runs to determine the fine LOD because all replicates amplified in first-round testing, including concentrations below the course LOD we determined in Stage II.

Finally, in Stage IV, we test each candidate primer set for cross-reactivity with other viruses in this study by using high-concentration nucleic acid extracts of target viruses. Primer sets showing no amplification at these high concentrations are unlikely to cross-react in complex samples which generally contain lower viral concentrations resulting from sample processing and dilution. Our designed primer sets did not cross-react within the timeframe determined by sensitivity and specificity analyses. BVDV-1.L.1 showed late partial cross-reactivity with BRSV at 50 minutes but did not reach plateau. (Figure S14) We did not consider it a significant concern since inhibitors in complex samples would likely further delay and suppress amplification. BRSV.M2.1 showed NTC amplification in two of four replicates around 45 minutes. Because BRSV.M2.1 was not cross-reactive with other viruses, we selected it as the final primer set but noted its potential for non-specific amplification.

Following selection of primer sets, we investigated the *in silico* inclusivity of designed primer sets. Interestingly, we found that generally, the primer sets we advanced had comparatively high cumulative inclusivity. We do note that the availability of viral sequences may limit this analysis in practice (for instance, we only had two sequences for BHV-1.pV targets). Nevertheless, when sufficient sequences are available, *in silico* inclusivity analysis provides another avenue for selecting primer sets to further save development time and resources.

### 4.4. Assay advantages, limitations, and future outlook

The LAMP assays we developed in this study, along with the methodology used for their development, offer three distinct advantages to the scientific community. Specifically, we note that (i) the LAMP assays presented here enable the detection of five viruses commonly implicated in the onset and progression of BRD from samples that are minimally processed and require only dilution in nuclease-free water. To our knowledge, the literature does not provide well-performing LAMP primer sets for these viruses. (ii) We validated the performance of these primer sets in diluted field samples and reported established LODs that are competitive with other LAMP primer sets present in the literature. We verified these LODs using a novel dPCR parallel quantification method, which, to our knowledge, has not been previously employed for this purpose in the literature. (iii) Finally, this study presents a robust and well-defined methodology for designing, screening, selecting, and characterizing LAMP primer sets. This process reduces the time and resources required to develop LAMP primer sets for novel nucleic acid targets.

This study’s progress has three limitations offering opportunities for further development: **(i)** We have validated these assays using real-time fluorescent data, but validation using colorimetric methods more amenable to field settings is still needed. We caution that end-point colorimetric methods may yield false positives, particularly in primer sets showing late-stage amplification in no-template reactions. **(ii)** Each LAMP assay requires about 30 minutes to reach requisite sensitivity, which may limit use in rapid, high-throughput field applications. Methods to accommodate or accelerate this reaction time are needed to increase field adoption. **(iii)** RNA virus primer sets have LODs above 100 copies per reaction and analytical sensitivities less than 95% potentially resulting from reverse transcriptase inhibition by sample inhibitors. Furthermore, we did not perform any sample-size calculation to reach a desired statistical power resulting in an under-powered evaluation of sensitivity and specificity. This shortcoming was driven by the low number of BRD-suspect cattle available at our collection site. Therefore, prior to adoption and translation to field settings, users should conduct large-scale, multisite field trials to validate performance and utility in real-world settings.

Future work to improve upon these advantages and limitations are: **(i)** Developers of colorimetric LAMP systems can use these assays to enable naked-eye detection of pathogen in field settings. Rather, we suggest real-time colorimetric image analysis with a simple imaging setup for field use instead of end-point image analysis. Recent studies develop compact imaging devices integrated with portable heaters and demonstrate LAMP detection in various settings and on paper-based substrates (Wang et al. 2021; Diaz et al. 2021; Papadakis et al. 2022; Wang et al. 2022; Wang et al. 2023; Strachan et al. 2023; Wang et al. 2024; Kamel et al. 2025). **(ii)** Assay developers could streamline the developed primer selection pipeline by devising LOD prediction models. Examples of such models could include artificial intelligence algorithms using primer and template sequences as input or using Stage I screening data to forecast downstream performance. We recommend integrating a primer validation step into this pipeline using samples testing positive for target pathogens with traditional diagnostics to determine clinical sensitivity and specificity before widespread field deployment. The required sample size should account for desired confidence intervals and pathogen prevalence (WOAH 2024). **(iii)** Finally, investigating the mechanisms underpinning the poor LODs of RNA viruses may provide insights applicable to a wider array of RNA-based enzymatic reactions beyond RT-LAMP. Researchers may use these insights to develop diagnostics for RNA pathogens traditionally requiring lab-based testing to achieve requisite analytical performance. Such innovations could transform rapid detection, deployment, and surveillance of emerging animal and zoonotic diseases.

## 5. Acknowledgments

The authors wish to thank Dr. Rebecca P. Wilkes and Ms. Katrina Miller for their assistance in the development of protocols for culturing and propagating the viruses investigated herein. Furthermore, we wish to thank Dr. Jiangshan Wang for his assistance in translating articles cited in this publication from Mandarin to English.

The following reagent was obtained through BEI Resources, NIAID, NIH: Bovine Parainfluenza Virus 3, SF-4, NR-3234.

## 7. Declarations and Disclosures

### 7.1. Animal Care and Ethics Declaration

All samples were collected with the approval of the Institutional Animal Care and Use Committee (IACUC), Protocol Number 1906001911.

### 7.2. Declaration of AI or AI-assisted technologies in the writing process

During the preparation of this work, the authors used artificial intelligence (AI; in particular, ChatGPT model 4o and Grammarly) to review grammar and provide limited stylistic improvements. The authors assert that all content, including modifications suggested by AI, has been reviewed and edited to convey the authors’ intent and original substance.

## 8. Author Contributions

**Josiah Levi Davidson:** Conceptualization, Methodology, Software, Validation, Formal Analysis, Investigation, Resources, Data Curation, Writing – Original Draft Preparation, and Writing – Review & Editing. **Murali Kannan Maruthumuthu** Conceptualization, Methodology, Investigation, and Resources. **Mohamed Kamel:** Methodology, Investigation, and Resources. **Suraj Mohan:** Conceptualization, Investigation, Methodology, and Resources. **Ana Pascual-Garrigos:** Methodology, Investigation, and Resources. **Andres Dextre:** Investigation and Resources. **Ruth Eunice Centeno-Delphia:** Methodology and Resources. **Jon Schoonmaker:** Conceptualization and Resources. **Timothy A. Johnson:** Conceptualization, Methodology, and Resources. **Jacquelyn P. Boerman:** Conceptualization and Methodology. **Deepti Pillai:** Conceptualization and Methodology. **Jennifer Koziol:** Conceptualization and Methodology **Aaron Ault** Conceptualization and Resources. **Mohit S. Verma:** Conceptualization, Methodology, Resources, Writing – Review & Editing, Supervision, Project Administration, Funding Acquisition.

## References

Ahmed B, Raut B, Pauley A, Davidson JL, Yang S, Verma MS. 2025. Development of a portable paper-based biosensor for the identification of genetically modified corn (*Zea mays*) and soybean (*Glycine max*). Biosens Bioelectron. 287:117690. 10.1016/j.bios.2025.117690

Ayaz Kök S, Üstün S, Taşkent Sezgin H. 2023. Diagnosis of Ruminant Viral Diseases with Loop-Mediated Isothermal Amplification. Mol Biotechnol. 65(8):1228–1241. 10.1007/s12033-023-00674-6

Blakebrough-Hall C, McMeniman JP, González LA. 2020. An evaluation of the economic effects of bovine respiratory disease on animal performance, carcass traits, and economic outcomes in feedlot cattle defined using four BRD diagnosis methods. J Anim Sci. 98(2):1–11. 10.1093/jas/skaa005

Boxus M, Letellier C, Kerkhofs P. 2005. Real Time RT-PCR for the detection and quantitation of bovine respiratory syncytial virus. J Virol Methods. 125(2):125–130. 10.1016/j.jviromet.2005.01.008

Buczinski S, Broes A, Savard C. 2024. Frequency of Bovine Respiratory Disease Complex Bacterial and Viral Agents Using Multiplex Real-Time qPCR in Quebec, Canada, from 2019 to 2023. Vet Sci. 11(12):631. 10.3390/vetsci11120631

Buczinski S, Forté G, Francoz D, Bélanger A M. 2014. Comparison of Thoracic Auscultation, Clinical Score, and Ultrasonography as Indicators of Bovine Respiratory Disease in Preweaned Dairy Calves. J Vet Intern Med. 28(1):234–242. 10.1111/jvim.12251

Bushnell B. 2014. BBMap: A Fast, Accurate, Splice-Aware Aligner. LBL Publ.(LBNL-7065E):2.

Calderón Bernal JM, Fernández A, Arnal JL, Baselga C, Benito Zuñiga A, Fernández-Garyzábal JF, Vela Alonso AI, Cid D. 2023. Cluster analysis of bovine respiratory disease (BRD)-associated pathogens shows the existence of two epidemiological patterns in BRD outbreaks. Vet Microbiol. 280:109701. 10.1016/j.vetmic.2023.109701

Centeno-Martinez RE, Glidden N, Mohan S, Davidson JL, Fernández-Juricic E, Boerman JP, Schoonmaker J, Pillai D, Koziol J, Ault A, et al. 2022. Identification of bovine respiratory disease through the nasal microbiome. Anim Microbiome. 4(1):15. 10.1186/s42523-022-00167-y

Chandranaik BM, Rathnamma D, Patil SS, Kovi RC, Dhawan J, Ranganatha S, Isloor S, Renukaprasad C, Prabhudas K. 2013. Development of a Probe Based Real Time PCR Assay for Detection of Bovine Herpes Virus-1 in Semen and Other Clinical Samples. Indian J Virol. 24(1):16–26. 10.1007/s13337-012-0112-1

Churchill KJ, Winder CB, Goetz HM, Wilson D, Uyama T, Pardon B, Renaud DL. 2025. Evaluating case definitions of respiratory disease in dairy calves: A scoping review. J Dairy Sci. 108(4):4030–4048. 10.3168/jds.2024-25827

Davidson JL, Wang J, Maruthamuthu MK, Dextre A, Pascual-Garrigos A, Mohan S, Putikam SVS, Osman FOI, McChesney D, Seville J, Verma MS. 2021. A paper-based colorimetric molecular test for SARS-CoV-2 in saliva. Biosens Bioelectron. 9:100076. 10.1016/j.biosx.2021.100076

Diaz LM, Johnson BE, Jenkins DM. 2021. Real-time optical analysis of a colorimetric LAMP assay for SARS-CoV-2 in saliva with a handheld instrument improves accuracy compared with endpoint assessment. J Biomol Tech JBT. 32(3):158–171. 10.7171/jbt.21-3203-011

Fan Q, Xie Zhixun, Xie L, Liu J, Pang Y, Deng X, Xie Zhiqin, Peng Y, Wang X. 2012. A reverse transcription loop-mediated isothermal amplification method for rapid detection of bovine viral diarrhea virus. J Virol Methods. 186(1):43–48. 10.1016/j.jviromet.2012.08.007

Fulton RW. 2020. Viruses in Bovine Respiratory Disease in North America. Vet Clin North Am Food Anim Pract. 36(2):321–332. 10.1016/j.cvfa.2020.02.004

Gaudino M, Nagamine B, Ducatez MF, Meyer G. 2022. Understanding the mechanisms of viral and bacterial coinfections in bovine respiratory disease: a comprehensive literature review of experimental evidence. Vet Res. 53(1):70. 10.1186/s13567-022-01086-1

Griffin D. 2014. The monster we don’t see: subclinical BRD in beef cattle. Anim Health Res Rev. 15(2):138–141. 10.1017/S1466252314000255

Hess AS, Shardell M, Johnson JK, Thom KA, Strassle P, Netzer G, Harris AD. 2012. Methods and recommendations for evaluating and reporting a new diagnostic test. Eur J Clin Microbiol Infect Dis. 31(9):2111–2116. 10.1007/s10096-012-1602-1

Huang X, Tang G, Ismail N, Wang X. 2022. Developing RT-LAMP assays for rapid diagnosis of SARS-CoV-2 in saliva. eBioMedicine. 75:103736. 10.1016/j.ebiom.2021.103736

Juge AE, Hall NJ, Richeson JT, Daigle CL. 2022. Using Canine Olfaction to Detect Bovine Respiratory Disease: A Pilot Study. Front Vet Sci. 9:902151. 10.3389/fvets.2022.902151

Kamel M, Davidson JL, Schober JM, Fraley GS, Verma MS. 2025. A paper-based loop-mediated isothermal amplification assay for highly pathogenic avian influenza. Sci Rep. 15(1):12110. 10.1038/s41598-025-95452-6

Kamel MS, Davidson JL, Verma MS. 2024. Strategies for Bovine Respiratory Disease (BRD) Diagnosis and Prognosis: A Comprehensive Overview. Animals. 14(4):627. 10.3390/ani14040627

Kishimoto M, Tsuchiaka S, Rahpaya SS, Hasebe A, Otsu K, Sugimura S, Kobayashi S, Komatsu N, Nagai M, Omatsu T, et al. 2017. Development of a one-run real-time PCR detection system for pathogens associated with bovine respiratory disease complex. J Vet Med Sci. 79(3):517–523. 10.1292/jvms.16-0489

Li J, Wang J, Zhang S, Zhang J, Guo L. 2019. Establishment of a new RT-LAMP detection assay for BRSV. Chin J Vet Sci. 39(5):867–871. 10.16303/j.cnki.1005-4545.2019.05.10

Li Jian-you LJ, Wang C, Guo L, He H. 2017. Establishment and Preliminary Application of Nano-PCR and LAMP Methods for Bovine Parainfluenza Type-3 Virus. China Anim Husb Vet Med. 44(10):2837–2844.

Mari V, Losurdo M, Lucente MS, Lorusso E, Elia G, Martella V, Patruno G, Buonavoglia D, Decaro N. 2016. Multiplex real-time RT-PCR assay for bovine viral diarrhea virus type 1, type 2 and HoBi-like pestivirus. J Virol Methods. 229:1–7. 10.1016/j.jviromet.2015.12.003

Mohan S, Pascual-Garrigos A, Brouwer H, Pillai D, Koziol J, Ault A, Schoonmaker J, Johnson T, Verma MS. 2021. Loop-Mediated Isothermal Amplification for the Detection of Pasteurella multocida, Mannheimia haemolytica, and Histophilus somni in Bovine Nasal Samples. ACS Agric Sci Technol. 1(2):100–108. 10.1021/acsagscitech.0c00072

Murray GM, O’Neill RG, More SJ, McElroy MC, Earley B, Cassidy JP. 2016. Evolving views on bovine respiratory disease: An appraisal of selected control measures – Part 2. Vet J. 217:78–82. 10.1016/j.tvjl.2016.09.013

Nickell JS, Hutcheson JP, Renter DG, Amrine DA. 2021. Comparison of a traditional bovine respiratory disease control regimen with a targeted program based upon individualized risk predictions generated by the Whisper On Arrival technology. Transl Anim Sci. 5(2):1–13. 10.1093/tas/txab081

O’Donoghue S, Waters SM, Morris DW, Earley B. 2025. A Comprehensive Review: Bovine Respiratory Disease, Current Insights into Epidemiology, Diagnostic Challenges, and Vaccination. Vet Sci. 12(8):778. 10.3390/vetsci12080778

Papadakis G, Pantazis AK, Fikas N, Chatziioannidou S, Tsiakalou V, Michaelidou K, Pogka V, Megariti M, Vardaki M, Giarentis K, et al. 2022. Portable real-time colorimetric LAMP-device for rapid quantitative detection of nucleic acids in crude samples. Sci Rep. 12(1):3775. 10.1038/s41598-022-06632-7

Pascual-Garrigos A, Maruthamuthu MK, Ault A, Davidson JL, Rudakov G, Pillai D, Koziol J, Schoonmaker JP, Johnson T, Verma MS. 2021. On-farm colorimetric detection of Pasteurella multocida, Mannheimia haemolytica, and Histophilus somni in crude bovine nasal samples. Vet Res. 52(1):126. 10.1186/s13567-021-00997-9

Peltzer D, Tobler K, Fraefel C, Maley M, Bachofen C. 2021. Rapid and simple colorimetric loop-mediated isothermal amplification (LAMP) assay for the detection of Bovine alphaherpesvirus 1. J Virol Methods. 289:114041. 10.1016/j.jviromet.2020.114041

Pratelli A, Cirone F, Capozza P, Trotta A, Corrente M, Balestrieri A, Buonavoglia C. 2021. Bovine respiratory disease in beef calves supported long transport stress: An epidemiological study and strategies for control and prevention. Res Vet Sci. 135:450–455. 10.1016/j.rvsc.2020.11.002

Rafiq N, Verma MS. Design and development of a field-deployable water bath for loop-mediated isothermal amplification assay. bioRxiv. Preprint:1–17. 10.1101/2024.05.14.594127

Sarmiento-Silva RE, Nakamura-Lopez Y, Vaughan G. 2012. Epidemiology, Molecular Epidemiology and Evolution of Bovine Respiratory Syncytial Virus. Viruses. 4(12):3452–3467. 10.3390/v4123452

Shen Y, Liu J, Zhang Y, Ma X, Yue H, Tang C. 2020. Prevalence and characteristics of a novel bovine adenovirus type 3 with a natural deletion fiber gene. Infect Genet Evol J Mol Epidemiol Evol Genet Infect Dis. 83:104348. 10.1016/j.meegid.2020.104348

Strachan S, Chakraborty M, Sallam M, Bhuiyan SA, Ford R, Nguyen N-T. 2023. Maximising Affordability of Real-Time Colorimetric LAMP Assays. Micromachines. 14(11):2101. 10.3390/mi14112101

Studer E, Schönecker L, Meylan M, Stucki D, Dijkman R, Holwerda M, Glaus A, Becker J. 2021. Prevalence of BRD-Related Viral Pathogens in the Upper Respiratory Tract of Swiss Veal Calves. Anim Open Access J MDPI. 11(7):1940. 10.3390/ani11071940

Thonur L, Maley M, Gilray J, Crook T, Laming E, Turnbull D, Nath M, Willoughby K. 2012. One-step multiplex real time RT-PCR for the detection of bovine respiratory syncytial virus, bovine herpesvirus 1 and bovine parainfluenza virus 3. BMC Vet Res. 8(1):37. 10.1186/1746-6148-8-37

Timsit E, Dendukuri N, Schiller I, Buczinski S. 2016. Diagnostic accuracy of clinical illness for bovine respiratory disease (BRD) diagnosis in beef cattle placed in feedlots: A systematic literature review and hierarchical Bayesian latent-class meta-analysis. Prev Vet Med. 135:67–73. 10.1016/j.prevetmed.2016.11.006.

Unal I. 2017. Defining an Optimal Cut-Point Value in ROC Analysis: An Alternative Approach. Comput Math Methods Med. 2017(1):3762651. 10.1155/2017/3762651

U.S. Food & Drug Administration. 2007. Statistical Guidance on Reporting Results from Studies Evaluating Diagnostic Tests - Guidance for Industry and FDA Staff. Rockville, MD, USA: U.S. Food and Drug Administration.

U.S. Food & Drug Administration. 2024. Enforcement Policy for Certain In Vitro Diagnostic Devices for Immediate Public Health Response in the Absence of a Declaration under Section 564 - Draft Guidance for Laboratory Manufacturers and Food and Drug Administration Staff. Rockville, MD, USA: U.S. Food and Drug Administration.

Wang J, Davidson JL, Kaur S, Dextre AA, Ranjbaran M, Kamel MS, Athalye SM, Verma MS. 2022. Paper-Based Biosensors for the Detection of Nucleic Acids from Pathogens. Biosensors. 12(12):1094. 10.3390/bios12121094

Wang J, Dextre A, Pascual-Garrigos A, Davidson JL, Maruthamuthu MK, McChesney D, Seville J, Verma MS. 2021. Fabrication of a paper-based colorimetric molecular test for SARS-CoV-2. MethodsX. 8:101586. 10.1016/j.mex.2021.101586

Wang J, Kaur S, Kayabasi A, Ranjbaran M, Rath I, Benschikovski I, Raut B, Ra K, Rafiq N, Verma MS. 2024. A portable, easy-to-use paper-based biosensor for rapid in-field detection of fecal contamination on fresh produce farms. Biosens Bioelectron. 259:116374. 10.1016/j.bios.2024.116374

Wang J, Ranjbaran M, Ault A, Verma MS. 2023. A loop-mediated isothermal amplification assay to detect *Bacteroidales* and assess risk of fecal contamination. Food Microbiol. 110:104173. 10.1016/j.fm.2022.104173

WOAH. 2024. Principles and methods of validation of diagnostic assays for infectious diseases. In: Man Diagn Tests Vaccines Terr Anim Thirteen Ed. 13th ed. Paris, France: World Organization of Animal Health; p. 99–123.

Wong K, Xagoraraki I. 2010. Quantitative PCR assays to survey the bovine adenovirus levels in environmental samples. J Appl Microbiol. 109(2):605–612. 10.1111/j.1365-2672.2010.04684.x

