## Supplementary File 1 for "Detection of five viruses commonly implicated with Bovine Respiratory Disease using loop-mediated isothermal amplification"

**for**

### 1. Detailed review of Primer Screening and Selection

#### 1.1. Primer Screening, Stage I: Characterization and scoring of candidate primer set performance

In Stage I, candidate primer sets are screened using fluorometric qLAMP/RT-qLAMP in nuclease-free water using nucleic acid extracts from cell culture supernatant at high concentrations. In this manner, we remove the effect of any potential inhibitors in the sample background and provide the best opportunity to measure a candidate primer set’s performance by only providing its intended target as the template in the reaction. We note that it is still possible for off-target interactions to occur at other locations in the target genome since we have provided the entire viral genome in this step; however, this is an advantage of our approach as we want to ensure we capture these interactions since they may also be present in the end-stage reaction where the entire genome may also be present.

We use an elevated concentration of full-length target nucleic acid extract (1.0 x 10^5^ copies per reaction) in nuclease-free water for Stage I screening. Figure S 6 – Figure S 13 shows the amplification curves for each designed primer set that we then used to calculate average amplification times. We qualitatively analyzed the performance of each candidate primer set gauging three characteristics: I) quick reaction time which we defined as the time at which exponential amplification began being quicker than 15 minutes, II) the lack of false positive amplification before the 30-minute time points, and III) consistent reaction profiles especially when it comes to reaction time.

Initially, we designed candidate primer sets to target three distinct regions of the target pathogens genome with each target region having three primer sets developed (except for BHV-1 where two regions had four primer sets). We noted poor performance in the BAV-3 and BHV-1 candidate primer sets. Specifically, the BAV-3 candidate primer sets performed poorly since only one candidate primer set produced no false positives (BHV-1.pV.2; Figure S 6). In this stage, we permitted candidate primer sets to continue to Stage II if they displayed late-stage false positives, which we defined as occurring near or later than 30 minutes, such as the case for BAV-3.pV.3. In an ideal scenario, however, at least three primer sets per pathogen would not produce false positives leaving us with alternative primer sets downstream should performance be inadequate in Stage II and Stage III screening. Even though the BHV-1 candidate primer sets did not produce false positives, all reactions were accompanied by a slow reaction time (slower than 15 minutes; BHV-1.UL23.1, BHV-1.UL23.2, and BHV-US8.1; Figure S 9). To address the shortage of well-performing primer sets for BAV-3 and BHV-1, we designed a second set of primers targeting three additional regions of the target genome (Figure S 7, Figure S 8, and Figure S 10).

Given the need to conduct another round of primer design and screening, we advise that a total of at least 15 candidate primer sets be designed for each target pathogen to minimize the possibility that less than three candidate primer sets pass Stage I screening to continue to Stage II. Whereas it is not strictly required, we further advise that this corresponds to at least five primer sets targeting three district regions on the target pathogen genome.

Once we screened all primer sets, we scored their Stage I performance using an algorithm we previously published (Kamel et al. 2025). This algorithm is based on a ranked and weighted average of the amplification intensity, reaction time, and number of false positives. We selected the weights to prefer high amplification consistency over magnitude in the amplification intensity, by a factor of two (5% for the average intensity, and 10% for the standard deviations). For the reaction time, however, we prioritized quicker reaction times at high template concentrations over reaction time consistency since reaction times will tend to become more stochastic with decreasing template concentrations in a manner that is not necessarily straightforward to select for at high template concentrations (unpublished data). Thus, we allotted 15% of the overall score to the average reaction time between replicates, while we gave the standard deviation of the reaction time only 10% of the overall score. We allotted 60% of the score for ranking the occurrence of false positives with each successive false positive incurring increasing penalties (i.e. the first false positive accounted for 6% of the overall score, the second accounted for 12%, the third for 18%, etc.)

We present the results of Stage I scoring in Table S 8 - Table S 12 with candidate primer sets listed in descending order of the overall score. At this step, we did not discriminate between regions on the viral genome and instead opted to group them all together to select the best candidate primer sets for each virus. We selected at most three primer sets with scores above 90 to proceed to Stage II of screening. If no primer set exceeded 90, as was the case for bPIV-3, we selected the top three ranked primer sets, regardless of the score. We note that even though low primer scores can advance to Stage II, in many instances, low primer scores may require designing additional candidate primer sets due to failing to meet the selection criteria for Stage II/Stage III. Primer sets were excluded from continuing to Stage II if any false positives occurred before the 30-minute time point.

Table 4 of the main text shows candidate primer sets that passed initial screening and advanced to Stage II along with their average amplification times. Of specific note, however, is that BHV-1 only had one primer set that advanced to stage II (BHV-1.gD.4) as the other primer set scoring above 90 displayed a false positive at 27 minutes (BHV-1.US8.2). In cases such as these, an additional stage of more careful primer design and screening will likely be needed. For our purposes, we continued to Stage II with only a single primer set for BHV-1 since we had already gone through one stage of primer redesign, and BHV-1.gD.4 displayed exceptional performance (it displayed the fastest reaction time of all candidate primer sets advanced to Stage II).

#### 1.2. Primer Screening, Stage II: Quantification of LOD for candidate primer sets

We structured Stage II to quantify the limit of detection (LOD) of candidate primer sets efficiently to minimize the number of reactions required. We refer to the preliminary LOD as the final LOD tested before conducting sensitivity and specificity analysis. For the primer screening and selection, we consider only the preliminary LOD in the absence of inhibitors in nuclease-free water. Once determined, the preliminary LOD in the absence of nuclease-free water can be used to determine the starting concentration (generally one to two orders of magnitude higher) to conduct a similar LOD analysis with complex sample.

To this end, as with Stage I, we use extracted nucleic acid from viral culture supernatant diluted in nuclease-free water to minimize the impact of any inhibitors present in complex sample. We set about determining the preliminary LOD by separating our testing into two: Stage II and III. Stage II uses a series of 10-fold dilutions to determine the upper decade to begin testing for the preliminary LOD. We term this range as the “coarse LOD”. Stage III uses a series of 2-fold dilutions beginning at the coarse LOD to determine the preliminary LOD. As discussed in detail during the Stage III screening below, we recommend changing this dilution to an alternating series of 5-fold and 2-fold dilutions to more quickly determine the preliminary LOD.

We utilized digital PCR (dPCR) conducted on a Qiagen QIAcuity 5-plex with primers and probes already validated and published in literature for the detection of each virus to obtain accurate quantification of nucleic acid extract used as a template in the qLAMP/RT-qLAMP reactions. For Stage II, we quantified the extracted viral template from the cell culture supernatant using dPCR before dilution. Using this quantification, we then diluted each aliquot to the final reaction concentration of 1.0 x 10^5^ copies per reaction needed for the starting concentration in the Stage II 10-fold dilution series. For the determination of preliminary LODs in Stage III, we used the coarse LOD determined in Stage II as the starting concentration for the 2-fold dilution series. Furthermore, we additionally conducted dPCR of the dilutions used at each concentration in parallel with qLAMP/RT-qLAMP in Stage III to ensure accurate quantification of the template in the final reaction.

Concentrations from dPCR assays were determined using a modified methodology based on that used by the manufacturer in their analysis software (QIAGEN 2023). Analysis of data was conducted using Python and is available in the published code which can be accessed according to the information in the “Data Availability” section of the main manuscript. The average copy number per μL of sample was rounded to one decimal point. When concentrations as determined by dPCR are reported as copies per reaction, the average copy number after accounting for the volume of sample in the reaction was rounded down to the nearest whole number, so as not to inflate the amount of template in the reaction. In a similar fashion, the lower bound of reported 95% confidence intervals on the concentration per reaction was rounded up while the upper bound was rounded down to the nearest whole number so as not to “expand” the confidence interval.

We selected one to two primer sets from the Stage II results for each virus to continue to Stage III. We selected primer sets using three criteria: I) the primer set must not show amplification in the NTC replicates, II) the primer set has the best LOD when compared to alternative primer sets, and III) the primer set must have an average reaction time faster than 20 minutes for all replicates at the concentration level of the LOD. For LOD determination, we did not allow for any false positives regardless of whether it was late-stage amplification or not. We determined the coarse LOD for Stage II as the lowest concentration level at which all replicates amplify and all replicates for all concentration levels greater than this limit also amplify. Furthermore, we require that at least one replicate fail to amplify in the concentration level immediately below the LOD to determine an accurate “cut-off” concentration for the LOD. In the event that more than one primer set satisfied all conditions, we advanced multiple primer sets for Stage III.

Figure S 14 through Figure S 16 show the fluorescent traces of Stage II screening of candidate primer sets advanced from Stage I. The average reaction time of candidate primer sets that we tested in Stage II at all tested concentration levels are shown in Table S 13 with candidate primer sets that we advanced to Stage III highlighted in black. The coarse LOD for all examined candidate primer sets was greater than or equal to 100 copies per reaction. Two viruses displayed multiple candidate primer sets each with equivalent coarse LODs: BAV-3 (all BAV-3 candidate primer sets; 100 copies per reaction) and BRSV (BRSV.M2.1 and BRSV.NS2.3; 1,000 copies per reaction). For BAV-3, two candidate primer sets (BAV-3.pV.2 and BAV-3.pV.3) performed almost identically at the coarse LOD (100 copies per reaction) with the exception that one replicate in the BAV-3.pV.3 reaction was delayed by about 5 minutes. We attributed this slight delay to a potential pipetting error. BAV-3.Penton.3, however, had a replicate that amplified around 40 minutes, and the remaining two replicates amplified about 5 minutes slower than the average reaction time of BAV-3.pV.2 and BAV-3.pV.3. As such, we did not advance BAV-3.Penton.3 to Stage III. Given the obvious performance advantage of BAV-3.pV.2/BAV-3.pV.3 and given they performed almost identically at the coarse LOD, we elected to advance both candidate primer sets to Stage III. Likewise, both primer sets for BRSV.M2.1 and BRSV.NS2.3 performed almost identically (with BRSV.NS2.3 having a slight delay of approximately one minute on average) and as such both were advanced to Stage III. All the other viruses had a single candidate primer set per virus that performed superior to the other candidate primer sets in terms of LOD (BHV-1.gD.4, bPIV-3.L.1, and BVDV-1.5UTR.3).

We also want to note that only one primer set (BVDV-1.E2.2) produced a false positive. BVDV-1.E2.2 did not produce a false positive during primer screening, so we re-tested this coarse LOD several times and each time the false positive remained or the number of false positives increased (unpublished data). The result shown in this publication was the best LOD that we were able to obtain from BVDV-1.E2.2 and it emphasizes that at this stage of the screening process, if any false positive occurs, the primer set should be rejected from further screening. We hypothesize that false positives occurring at this stage are likely due to the primer set beginning to lose stability as a result of freeze-thaw cycles or degradation over time which would ultimately translate to an untenably short shelf life for a final diagnostic. Consequently, BVDV-1.E2.2 was not advanced to stage III.

For Stage III screening, we began with the coarse quantification limit determined in Stage II and conducted a series of qLAMP/RT-qLAMP reactions at two-fold dilutions to determine the fine LOD. The same criteria above were used to determine the concentration we designated as the LOD. To increase confidence in our reported LOD, we quantified the template using dPCR in parallel with qLAMP/RT-qPCR rather than assuming its concentration following a dilution series as we did in Stage II. During Stage II, such an assumption was warranted as we only needed to determine an approximate LOD (the coarse LOD) to guide our testing and serve as the starting concentration in Stage III and we would be conducting another test to verify that LOD before determining the preliminary LOD. For Stage III, however, our determination of the preliminary LOD (in this case dPCR) will be final pending verification by sensitivity and specificity analysis. The sensitivity and specificity analysis is quite costly (in terms of reagents and samples) and time-consuming. Therefore, an additional validation method is desired to ensure we are reporting preliminary LODs in order to minimize the potential for having to repeat the sensitivity and specificity analysis at an alternative concentration due to improper determination of the preliminary LOD. In this manner, we can account for any retention and annealing of nucleic acids to plastics during the dilution series along with pipetting error to ensure the actual concentration of the template tested was of a comparable magnitude to that reported. Given we are quantifying at lower concentration levels for Stage III, we elected to use dPCR with a higher partition count (as compared to those used for Stage II quantification) to obtain a quantification with more confidence at the concentrations we are testing.

Figure S 17 and Figure S 18 display the fluorometric traces of the qLAMP/RT-qLAMP amplification curves for candidate primer sets which we advanced to Stage III. BHV-1.gD.4 performed surprisingly well at all concentrations tested when we began the two-fold dilutions at 100 copies/reaction. (Figure S 17) Given that all replicates at all concentrations amplified and the NTC replicates were clear, we could not definitively define an LOD since no replicates failed to amplify and thus another LOD test was needed starting at a lower concentration; we termed these as Round 1 and Round 2 tests. Upon initial inspection, we tested two concentration levels in Round 1 which were below 10 copies/reaction and showed consistent amplification; (Figure S 17) however, this stands in contrast with the 10 copies/reaction concentration level which had a single replicate fail to amplify in the Stage II testing. (Figure S 14) bPIV-3.L.1 presented a similar conundrum when beginning Round 1 at 1,000 copies/reaction as indicated by Stage II results (Figure S 15) and amplified below 100 copies/reaction in Stage III. (Figure S 18) We attributed these apparent conflicts to the adhesion of templates and amplification of user error when conducting the dilutions at such a low concentration level. To verify the concentrations are within an acceptable range of what we expect given the initial quantification of the template, we conducted dPCR for any experiment where we expected concentrations to approach the LOD of a candidate primer set. This phenomenon was first observed for the bPIV-3.L.1 primer set starting at 1,000 copies/reaction, and when this set was taken we did not yet conduct the dPCR cross-validation in parallel with qLAMP/RT-qLAMP. Beginning with the Stage III quantification at the lower starting concentration level (100 copies/reaction) for bPIV-3.L.1, we ran dPCR cross-validation in parallel for any qLAMP/RT-qLAMP reaction we expected to approach a primer set’s preliminary LOD (including both of the BHV-1 starting concentration ). Additionally, for LOD studies in samples (and a future modification to this testing process which we now advise), we used a series of alternating 5-fold and 2-fold dilutions to more quickly arrive at the preliminary LOD and minimize the probability that we would have to conduct serial studies to determine the preliminary LOD.

The preliminary LOD was reported as the lowest *predicted* concentration level that is greater than the *actual* concentration measured via dPCR at the concentration level indicated as the preliminary LOD according to qLAMP/RT-qLAMP results. (Main Text, Table 2) For instance, the lowest concentration level at which all replicates amplified using bPIV-3.L.1 was at a predicted concentration of 25 copies/reaction. (Figure S 18, Table S 14) The concentration we measured via dPCR was 42 copies/reaction (average of 3 replicates) with a 95% confidence interval of 34 and 52 copies/reaction. (Table S 15) The closest predicted concentration level we tested that lies within or above this confidence interval was 50 copies/reaction, and thus was the reported preliminary LOD. We want to emphasize that the reported preliminary LOD does not necessarily correlate to the predicted concentration at which all three replicates are amplified, as is the case for bPIV-3. In this manner, we can be sure that – accounting for dilution variation and other sources of error – our preliminary LOD is at least what we reported, even though the true preliminary LOD may be less than the reported value.

BAV-3.pV.2 and BAV-3.pV.3 displayed the same coarse LOD of 100 copies/reaction in Stage II. (Table S 13) For the fine LOD, however, it is apparent that BAV-3.pV.3 has an LOD (12.5 copies/reaction) that is approximately four-fold lower than that of BAV-3.pV.2 (50 copies/reaction; Figure S 17 and Table S 14). This is confirmed by the dPCR results for BAV-3, and since we ran both primer sets at the same time using the same dilution series, we can be relatively confident that this difference in detection levels is valid. (Table S 15) Additionally, BAV-3.pV.2 began to exhibit amplification in one of the NTC reactions potentially indicating a similar stability problem as that of BVDV-1.E2.2. (Figure S 17) Due to this false positive amplification in the NTC reaction, we could not definitively determine a preliminary LOD for BAV-3.pV.2. Consequently, we chose BAV-3.pV.3 for further validation in field samples and ended screening for BAV-3.

BHV-1.gD.4 also displayed a coarse LOD of 100 copies/reaction in Stage II, (Table S 13) but the preliminary LOD was located somewhere below 12.5 copies/reaction (10 copies/reaction via dPCR quantification), requiring an additional round of testing to determine the LOD. For round 2, we used a dilution series beginning at 25 copies/reaction to determine the LOD. Due to ambiguities in the performance of the RT-qLAMP reaction at a concentration level of 3.125 copies/reaction and discrepancies in the dPCR parallel quantification for the two rounds, we determined that this primer set has a preliminary LOD of 6.25 copies/reaction. (See Section 1.4 for a comprehensive discussion on these discrepancies)

For the RNA viruses, bPIV-3.L.1 displayed amplification for all replicates down to the 25 copies/reaction concentration level which is the lowest of all of the RNA viruses. (Figure S 18) As with BHV-1.gD.4, all replicates amplified at all concentration levels tested using bPIV-3.L.1 when beginning the round 1 dilution series at 1,000 copies/reaction. Thus, similarly, we conducted a second dilution series, but in the case of bPIV-3, we began with a concentration of 100 copies/reaction which is a ten-fold dilution from the starting concentration used in round 1. In the round 2 dilution series, we see that 25 copies/reaction is the last concentration level at which all replicates amplify. (Figure S 18) As described in the example we previously discussed, the dPCR parallel quantification of this concentration level indicated that the concentration of template present in this reaction was 42 copies per reaction. (Table S 15) Thus, we report an LOD for bPIV-3.L.1 as 50 copies/reaction since it is included in the confidence intervals for the dPCR parallel quantification of this concentration level.

BRSV.M2.1 had all replicates amplify for the 500 copies/reaction predicted concentration level. dPCR parallel quantification results indicate that the true concentration of this level, however, is 816 copies/reaction with a 95% confidence interval between 788 and 844 copies per reaction. As a result, we report the LOD for BRSV.M2.1 as the next highest predicted level tested which is 1,000 copies/reaction. Of all the primer sets that we designed and screened, BRSV.M2.1 was the worst performing in terms of limit of detection. This is interesting to note, however, because even at this LOD, BRSV.M2.1 is the best-performing primer set in terms of amplification speed with an average reaction time of 8 minutes.

Finally, BVDV-1.5UTR.3 has a predicted and reported LOD of 250 copies/reaction which is confirmed by dPCR parallel quantification which has a measured LOD of 235 copies/reaction with a 95% confidence interval between 194 copies and 276 copies. We do note, though, that even though this LOD may be lower than BRSV.M2.1 and thus might appear to perform “better” than BRSV.M2.1, this primer set is the slowest and most inconsistent at the LOD with an average reaction time of 36 minutes. (Table S 14) Whereas we did advance BVDV-1.5UTR.3 to the final stage of screening, we did so with caution due to its performance.

#### 1.3. Primer Screening, Stage III: Determination of cross-reactivity of candidate primer set

To ensure viruses were specific for their intended targets, we conducted a series of cross-reactivity reactions against the four other viruses being investigated for each selected primer set using genomic extracts diluted in water. We designed these Stage III cross-reactivity tests to provide the highest probability of interactions with the tested pathogens. As such, we tested interactions between primer sets and a high concentration (1.0 x 10^4^ copies/reaction) of the extracted nucleic acid of the investigated viruses. We tested each virus separately to ensure that reactions between viral templates would not confound results for off-target interactions with our designed primer sets. We chose a concentration of 1.0 x 10^4^ copies/reaction as all primer sets exhibited similar performance at 1.0 x 10^4^ copies/reaction and 1.0 x 10^5^ copies/reaction in terms of reaction time and consistency (Figure S 14 – Figure S 16; Table S 13). Thus, we decided that 1.0 x 10^4^ copies/reaction was a sufficient concentration to interrogate any off-target interactions with other viruses.

Figure S 19 shows the results of the fluorescent qLAMP/RT-qLAMP amplification curves for each primer set when tested against each virus in this study. As expected, all primer sets show clear amplification in all replicates when tested against the viruses we designed them to target. The only primer set that shows any off-target interactions with another virus we tested was BVDV-1.5UTR.3 which showed some late-stage interactions with BRSV after 50 minutes. Given the NTC replicates are clear for BVDV-1.5UTR.3, this amplification is likely arising from some off-target interaction with BRSV. We did not find reason to be concerned, however, since it occurred so late in the reaction and was easily distinguishable from. Additionally, even though BVDV-1.5UTR.3 tends to perform inconsistently near the LOD, we expect that these off-target interactions would be further mitigated in complex solution when the nucleic acid is not as easily accessible, further alleviating any concerns we might have with this primer set as a diagnostic candidate. We do want to note that BRSV.M2.1, however, shows substantial amplification (albeit late-stage) in the NTCs indicating potential stability issues or cross-contamination issues since BRSV.M2.1 amplifies so intensely and early. Given the intensity of the non-specific amplification in the NTC, we caution against its usage in end-point detection methods, but given the time delay in the appearance of the amplification in NTCs, BRSV.M2.1 is a viable diagnostic tool for real-time diagnostic methods, such as the fluorescent RT-qLAMP mentioned here.

All primer sets passed stage III screening since none of them displayed substantial off-target amplification occurring at similar times as true amplification with the intended target. We do note, however, that this cross-reactivity study only tested specificity with other viruses used in this study; further investigation is needed to see if there may be interactions with other viruses that may infect the bovine respiratory tract.

#### 1.4. Supplemental Discussion on dPCR Quantification for LOD of BHV-1.gD.4 using nucleic acid extract

BHV-1.gD.4 had all replicates amplify at all concentration levels tested when starting at 100 copies/reaction, (Table S 14) the concentration level indicated by Stage II screening. (Table S 13) As a result, we needed to run a second round of concentration levels to determine where the true LOD was located as round 1 was inconclusive. For round 2, we selected a starting concentration of 25 copies/reaction. We selected this starting level to ensure we tested at least 2 concentration levels above and below the ending concentration level of round 1 (3.125 copies/reaction).

After re-running the LOD at lower concentration levels, we see the first replicate which failed to amplify at 3.125 copies/reaction, which had previously amplified in Round 1. dPCR parallel quantification (which at these levels are approaching the detection limit of dPCR for the number of partitions we used) for Round 1 shows that at 3.125 copies/reaction, the 95% confidence bounds produced a negative concentration indicating that we may begin to have some uncertainty in the parallel quantification at this concentration. (Table S 15) Furthermore, the dPCR parallel quantification for round 2 indicated that both the 6.25 and 3.125 copies/reaction concentration levels have practically the same average quantification: 4 copies per reaction (95% confidence interval of 3 – 6 and 1 – 7 copies per reaction for the 6.25 copies/reaction and 3.125 copies/reaction levels, respectively). The larger confidence intervals around the 3.125 copies/reaction dPCR quantification which enclose the entirety of the quantification at the concentration above (6.25 copies/reaction) render us unconfident in the dPCR quantification at or below 3.125 copies/reaction in round 2 due to limitations of the dPCR method at this low concentration without expanding the number of partitions.

Given this discrepancy in observed performance, lack of confidence in the dPCR parallel quantification at 3.125 copies/reaction in either round, and acknowledging we had additional failed amplifications at both concentration levels below 3.125 copies/reaction concentration level in Round 2, we set the preliminary LOD for BHV-1.gD.4 at 6.25 copies/reaction. We can still be confident in the dPCR parallel quantification results for round 2 at 6.25 copies per reaction, which confirms the measured concentration includes 6.25 copies/reaction.

The dPCR machine we used in this study, the Qiagen Qiacuity, enables “hyperwelling” which simply put, allows us to increase the number of partitions (and the confidence in quantification at lower concentration levels) by counting three separate wells as an “extension” of one another. Whereas the technical implementation of this is simply a software change, functionally, we must ensure that each well contains the same sample so what we are doing by “hyperwelling” in practice is simply increasing the volume that is interrogated by dPCR. We ran the dPCR quantification in triplicate on a 24-well plate to ensure we had the most partitions in a single well. Given space limitation on a 24-well plate, we would not have been able to run a hyperwell version of LOD concentrations in triplicate (this would require three wells per hyperwell X three hyperwells per concentration level X seven concentration levels, including NTC = 43 wells of 26k partitions, and we have at most 24). If required to quantify at such a low concentration level, especially during sensitivity and specificity testing, we would advise using such a hyperwelling approach. In the case of sensitivity and specificity tests, only 2 concentration levels are required (namely equivalent to and twice the concentration of the preliminary LOD) plus an NTC condition. Even that, however, would require that we only run the NTCs in duplicate not triplicate as we were again constrained by space on the QIAcuity® 26k reaction plate. To run both levels and the NTC condition in triplicate using 3 actual wells per hyperwell would require 27 wells, while the 26k plate only has 24 wells.

### 2. Determination of optimal sample dilution for reverse transcription loop-mediated isothermal amplification assay

Complex samples contain a variety of inhibitors that can impact the performance of dPCR and LAMP amplification (Nixon et al. 2014). Consequently, we need to determine an optimal sample concentration that is capable of detecting our target virus with qLAMP and quantifying our target virus with dPCR. To determine the optimal sample concentration that significantly impacts quantification, we simultaneously varied both the final reaction sample concentration and the target virus concentration that we sought to quantify. This approach enabled us to simultaneously investigate the impact of inhibitors in complex samples across a range of target concentrations. Given that dPCR is capable of quantifying viruses extraction-free and is less sensitive to inhibitors than LAMP amplification, any complex sample concentration at which quantification does not differ significantly from quantification in the absence of inhibitors (nuclease-free water) using dPCR is suitable for conducting LAMP amplification, so long as the final reaction concentration remains constant in both formats (Nixon et al. 2014; Pavšič et al. 2016). To account for potential impacts on the reverse transcription process required or RNA targets, we determine the optimal sample concentration for BVDV-1, an RNA virus.

### 3. Supplementary Figures

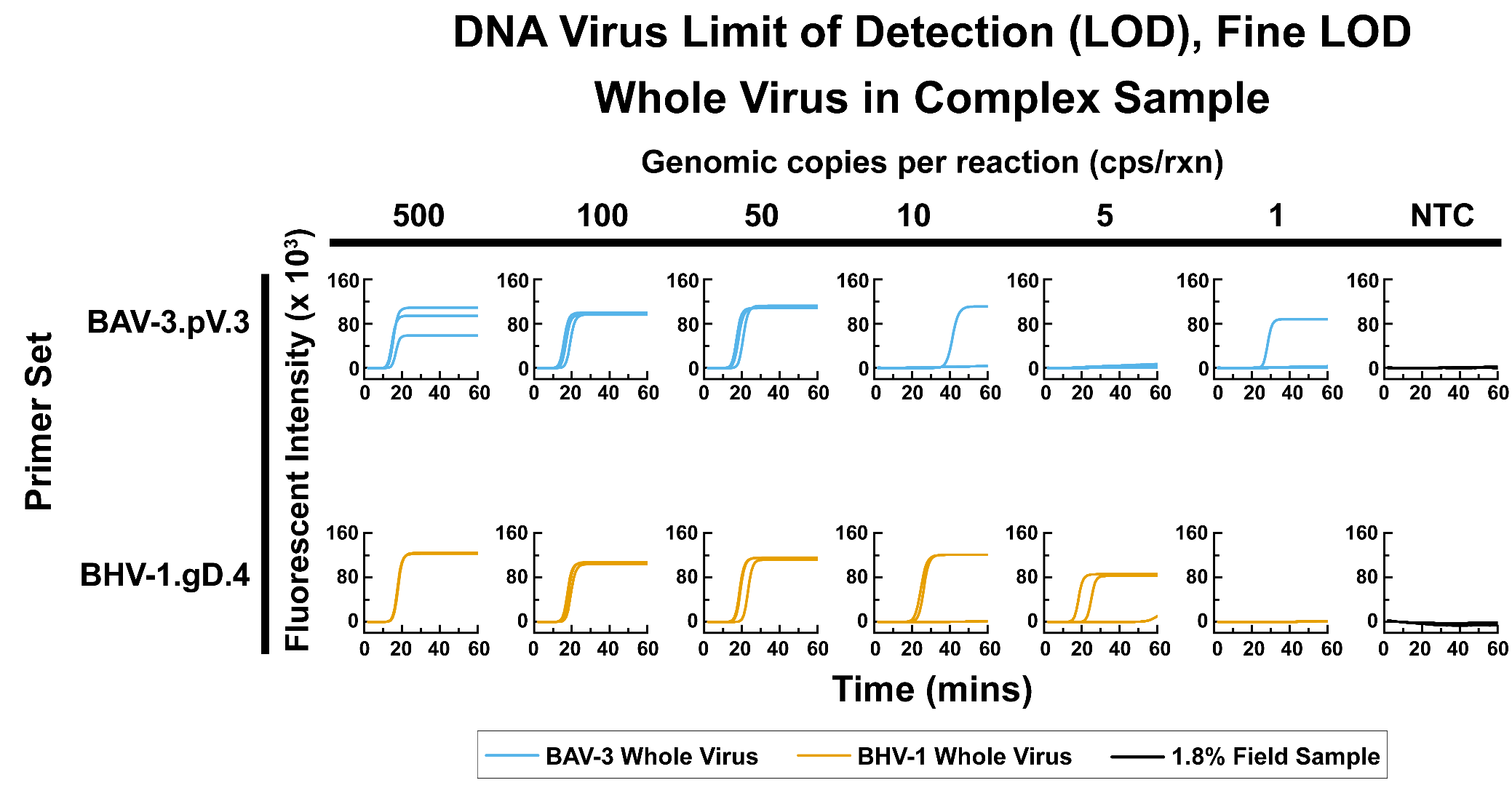

Figure S 1: Fine LOD for selected primer sets targeting DNA viruses in this study in complex field sample. Where indicated, a 2x solution of viral culture supernatant (untreated and unextracted) was diluted in complex field sample which had been prediluted to 9% using DEPC-treated nuclease-free water. 5 µL of the resulting dilution was then added to 20 µL of qLAMP mastermix resulting in a final reaction volume of 25 µL. For NTC reactions, 5 µL of the complex field sample pre-diluted to 9% using DEPC-treated nuclease-free water was added to 20 µL of qLAMP master mix resulting in a final reaction volume of 25 µL.

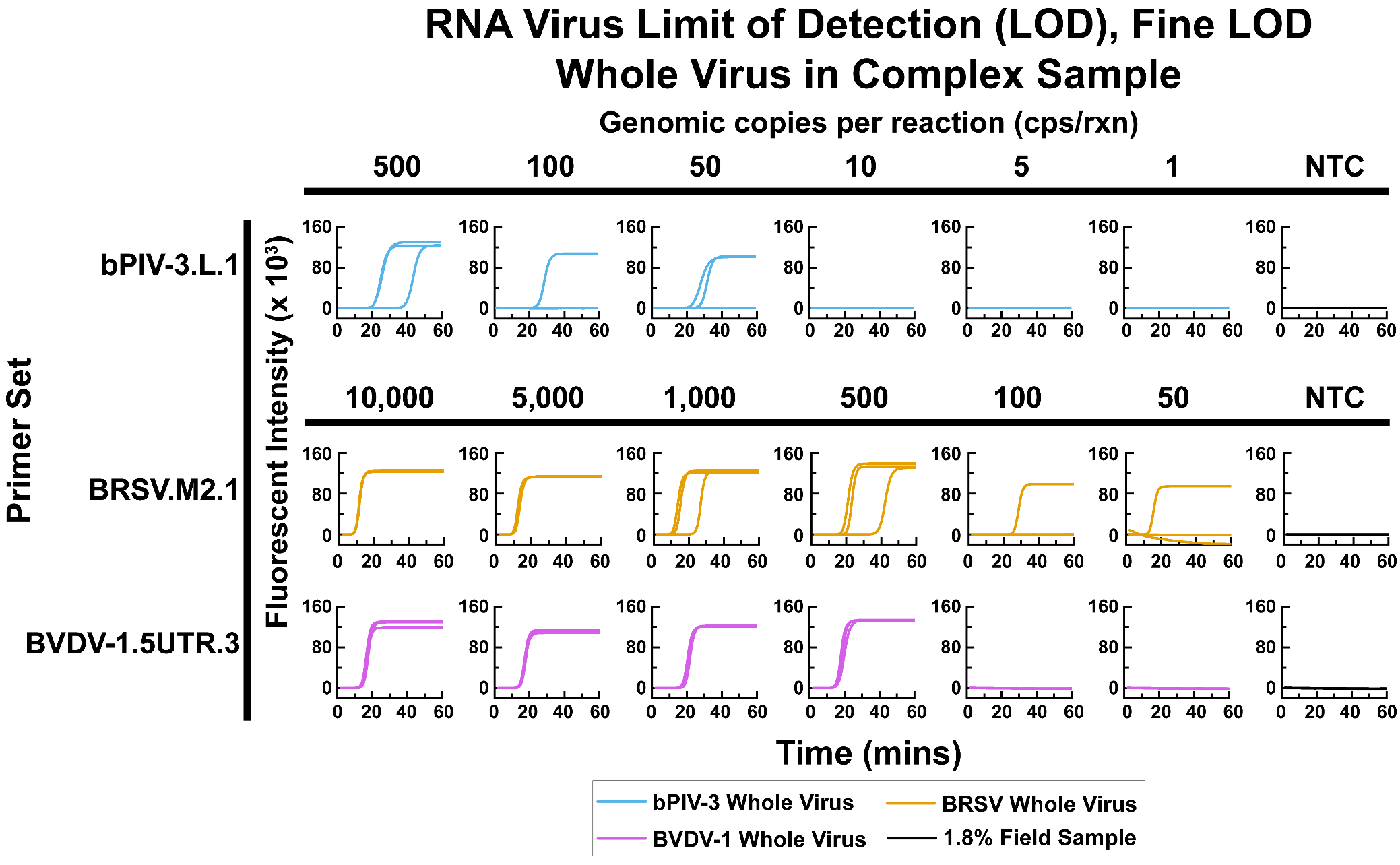

Figure S 2: Fine LOD for selected primer sets targeting RNA viruses in this study in complex field sample. Where indicated, a 2x solution of viral culture supernatant (untreated and unextracted) was diluted in complex field sample which had been prediluted to 9% using DEPC-treated nuclease-free water. 5 µL of the resulting dilution was then added to 20 µL of qLAMP mastermix resulting in a final reaction volume of 25 µL. For NTC reactions, 5 µL of the complex field sample pre-diluted to 9% using DEPC-treated nuclease-free water was added to 20 µL of qLAMP master mix resulting in a final reaction volume of 25 µL.

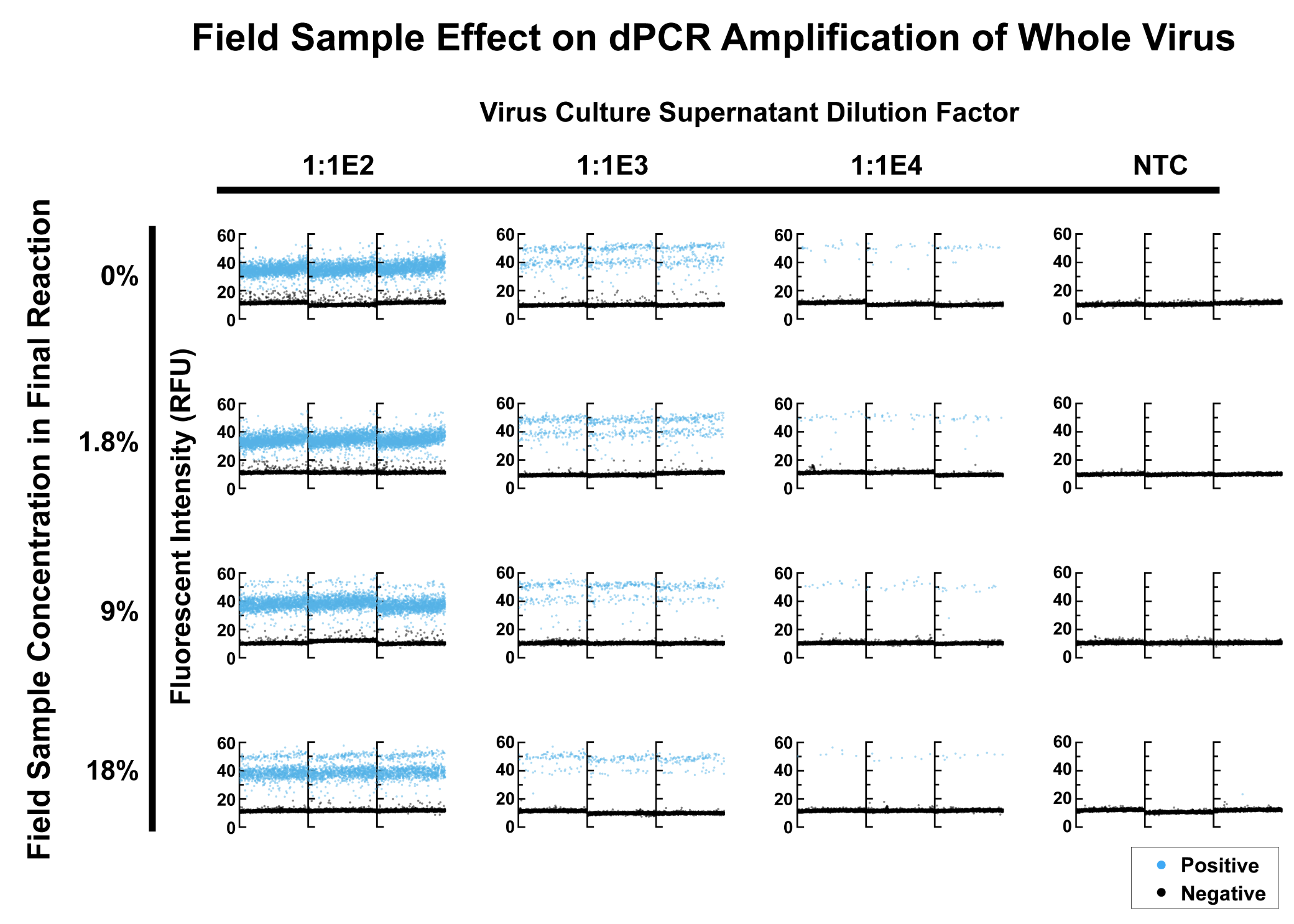

Figure S 3: Digital PCR (dPCR) amplification of BVDV-1 Virus culture supernatant (whole virus, untreated and unextrated) in various concentrations of complex sample background. Reactions are presented in triplicate. Virus culture supernatant at various dilutions was first diluted 1:10 in field samples appropriately diluted in nuclease-free water to arrive at the final dilution factor listed. For No Template Control (NTC) reactions, nuclease-free water was used in place of virus culture supernatant. 5 µL of either the diluted virus culture supernatant in complex sample or nuclease-free water was then mixed well with 15 µL of dPCR master mix. A common threshold of 20 RFU was used to determine positive partition amplifications from negative partition amplifications on the green fluorescent channel.

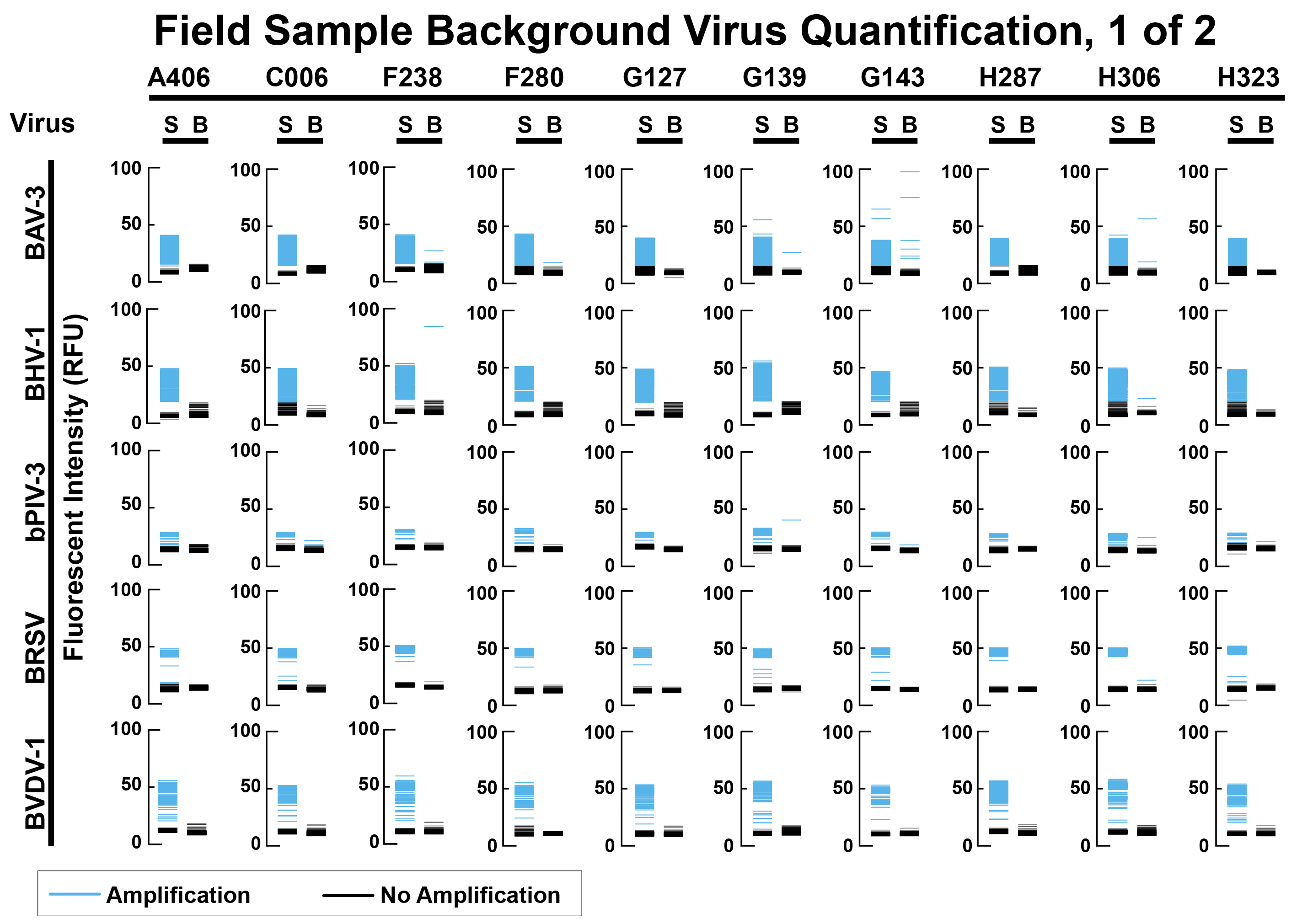

Figure S 4: Quantification of viruses in processed field samples via dPCR. Samples were diluted to a final reaction concentration of 2.25% using nuclease-free water. Whole viruses from cell culture supernatant were spiked into reactions containing the indicated field sample (labeled “S” for “spiked” and located on the left of each graph) at a concentration of 1,000 copies per reaction to verify assay conditions were sufficient for detection. Background sample reactions are labeled “B” which are located to the right of each graph, containing 5 µL of background sample with no virus spiked in. Blue indicates the digital partition exceeded the threshold value and is considered a positive amplification/partition indicative of the presence of the target virus. Threshold values were 15, 19, 18, 18, and 18 RFU for BAV-3, BHV-1, bPIV-3, BRSV, and BVDV-1, respectively.

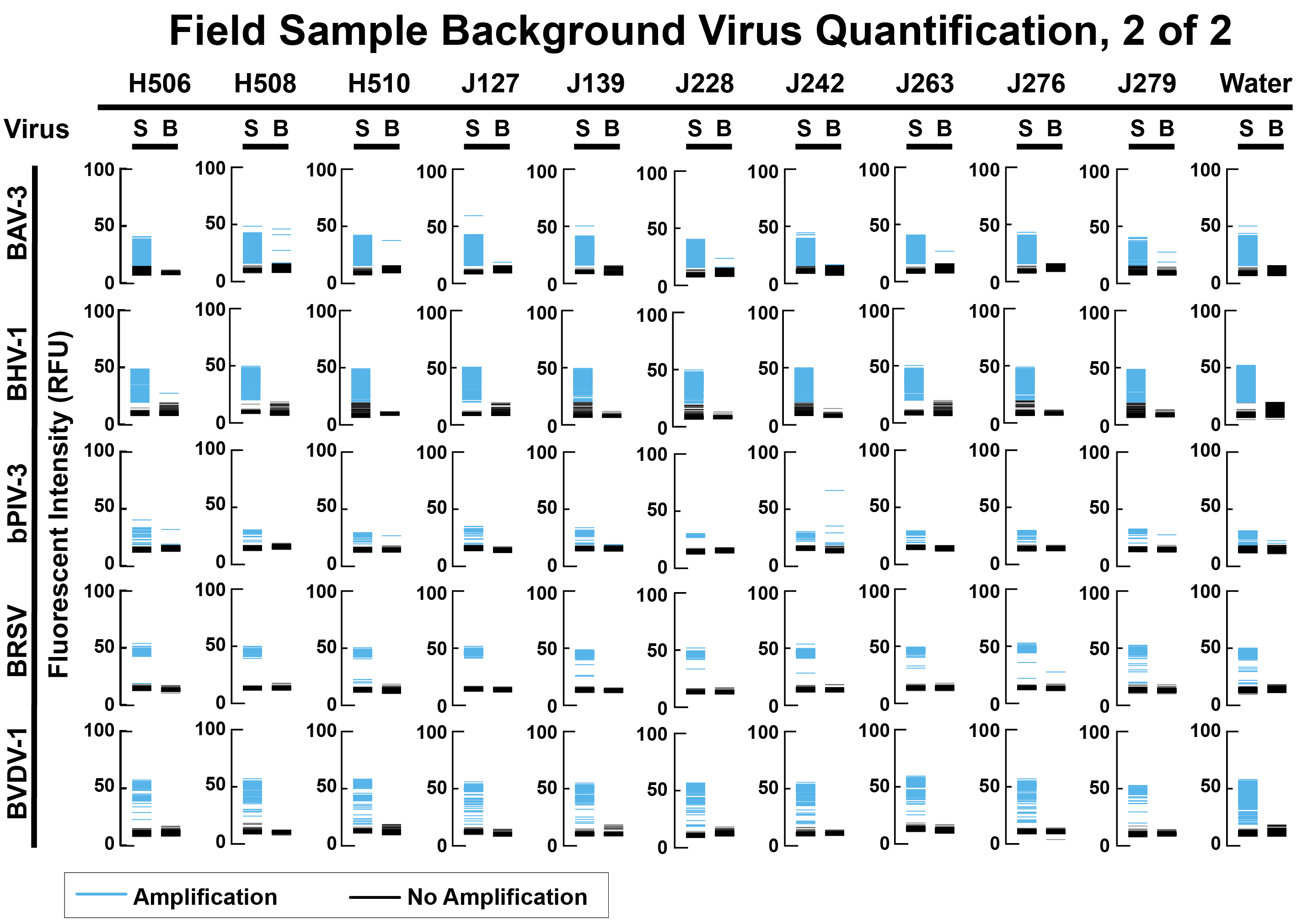

Figure S 5: Quantification of viruses in processed field samples via dPCR. Samples were diluted to a final reaction concentration of 2.25% using nuclease-free water. Whole viruses from cell culture supernatant were spiked into reactions containing the indicated field sample (labeled “S” for “spiked” and located on the left of each graph) at a concentration of 1,000 copies per reaction to verify assay conditions were sufficient for detection. Background sample reactions are labeled “B” which are located to the right of each graph, containing 5 µL of background sample with no virus spiked in. Blue indicates the digital partition exceeded the threshold value and is considered a positive amplification/partition indicative of the presence of the target virus. Threshold values were 15, 19, 18, 18, and 18 RFU for BAV-3, BHV-1, bPIV-3, BRSV, and BVDV-1, respectively. Sample controls are located in the column indicated “Water” wherein all other reaction conditions except for the presence of the sample were maintained.

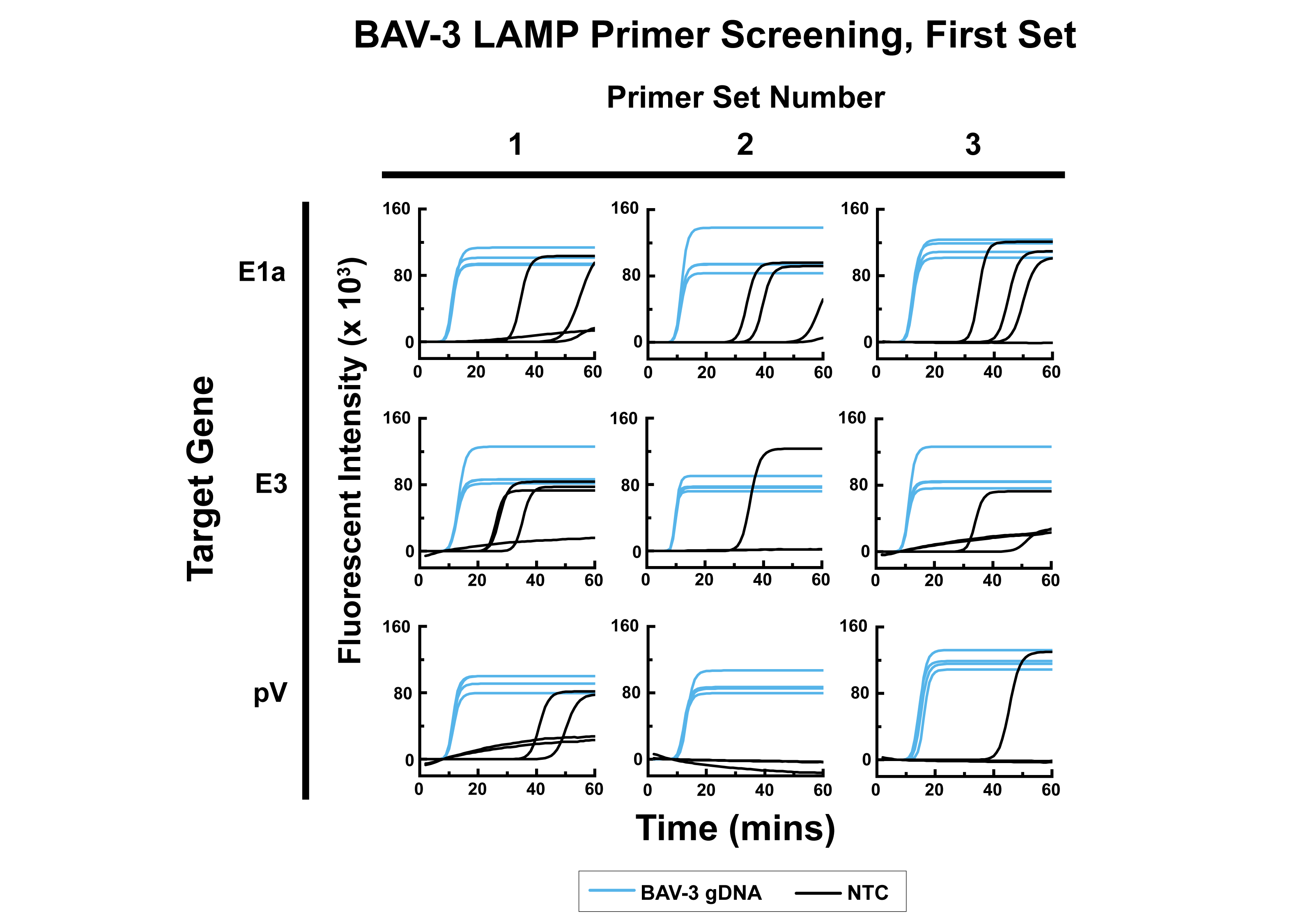

Figure S 6: Fluorometric traces for screening of the first round of LAMP primer sets targeting BAV-3. The target gene is along the vertical axis while the primer set ID for that gene is indicated along the horizontal axis. Reactions consist of 20 µL of LAMP master mix combined with either 5 µL of BAV-3 gDNA extract from virus culture supernatant at 2.0 x 10^4^ copies/µL resulting in a final reaction containing 1.0 x 10 ^5^ copies/reaction (blue lines) or 5 µL of nuclease-free water (black lines). The total reaction volume is 25 µL. Reactions were heated at 65 °C for 60 minutes and the fluorescent intensity was read every 60 seconds.

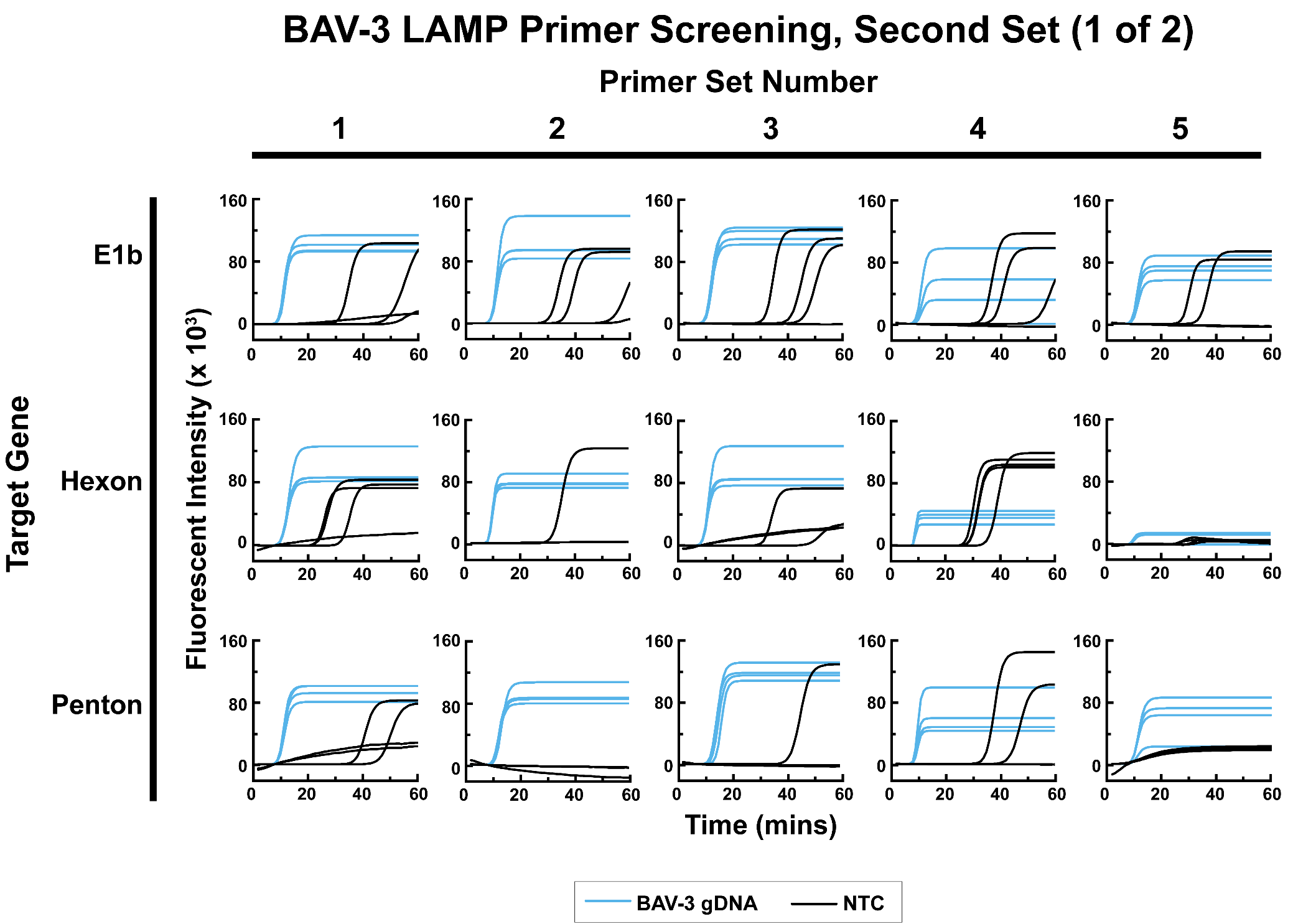

Figure S 7: Fluorometric traces for screening of the second round of LAMP primer sets targeting BAV-3 for primer set IDs between 1 and 5. The target gene is along the vertical axis while the primer set ID for that gene is indicated along the horizontal axis. Reactions consist of 20 µL of LAMP master mix combined with either 5 µL of BAV-3 gDNA extract from virus culture supernatant at 2.0 x 10^4^ copies/µL resulting in a final reaction containing 1.0 x 10 ^5^ copies/reaction (blue lines) or 5 µL of nuclease-free water (black lines). The total reaction volume is 25 µL. Reactions were heated at 65 °C for 60 minutes and the fluorescent intensity was read every 60 seconds.

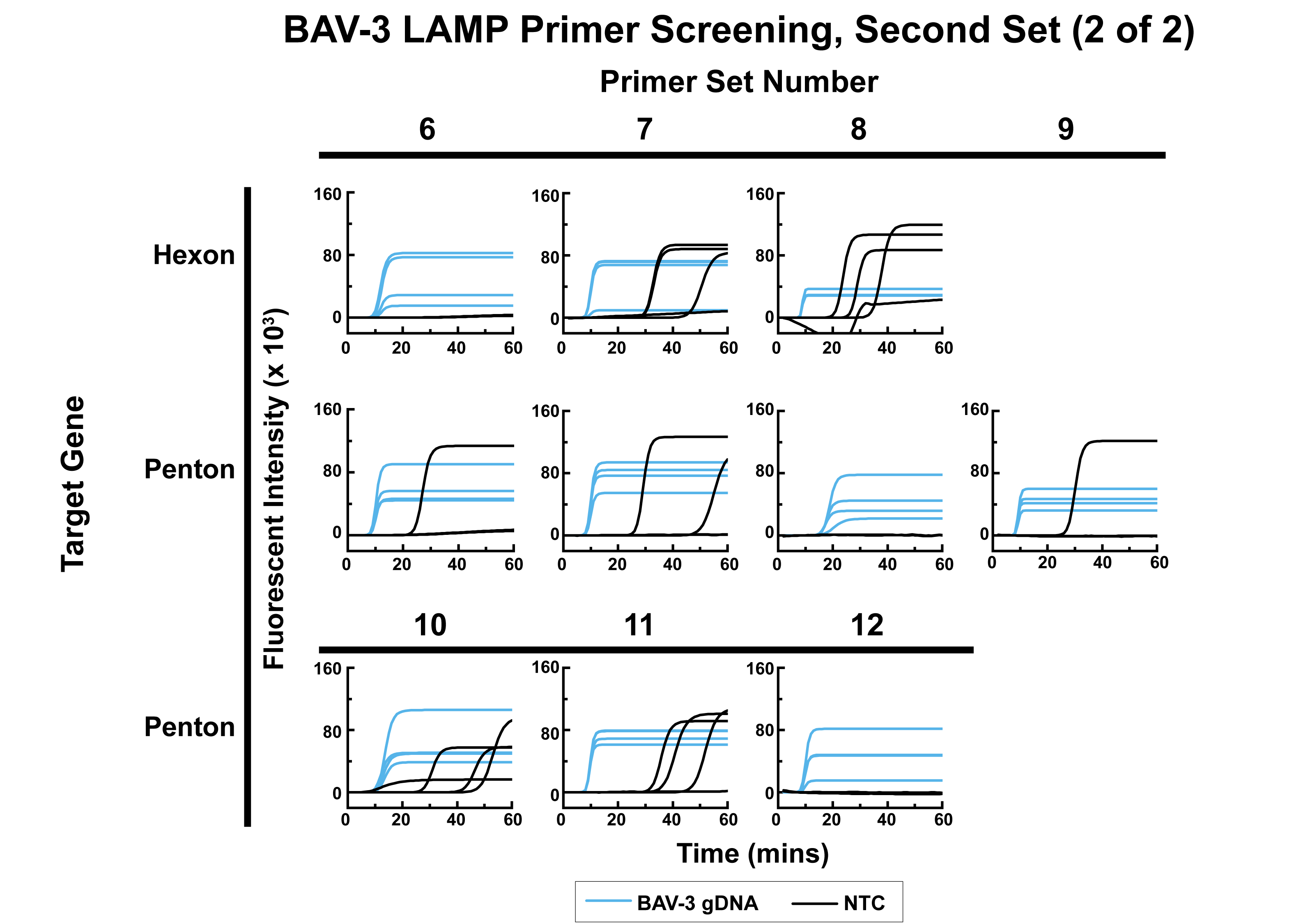

Figure S 8: Fluorometric traces for screening of the second round of LAMP primer sets targeting BAV-3 for primer set IDs between 6 and 12. The target gene is along the vertical axis while the primer set ID for that gene is indicated along the horizontal axis. Reactions consist of 20 µL of LAMP master mix combined with either 5 µL of BAV-3 gDNA extract from virus culture supernatant at 2.0 x 10^4^ copies/µL resulting in a final reaction containing 1.0 x 10 ^5^ copies/reaction (blue lines) or 5 µL of nuclease-free water (black lines). The total reaction volume is 25 µL. Reactions were heated at 65 °C for 60 minutes and the fluorescent intensity was read every 60 seconds.

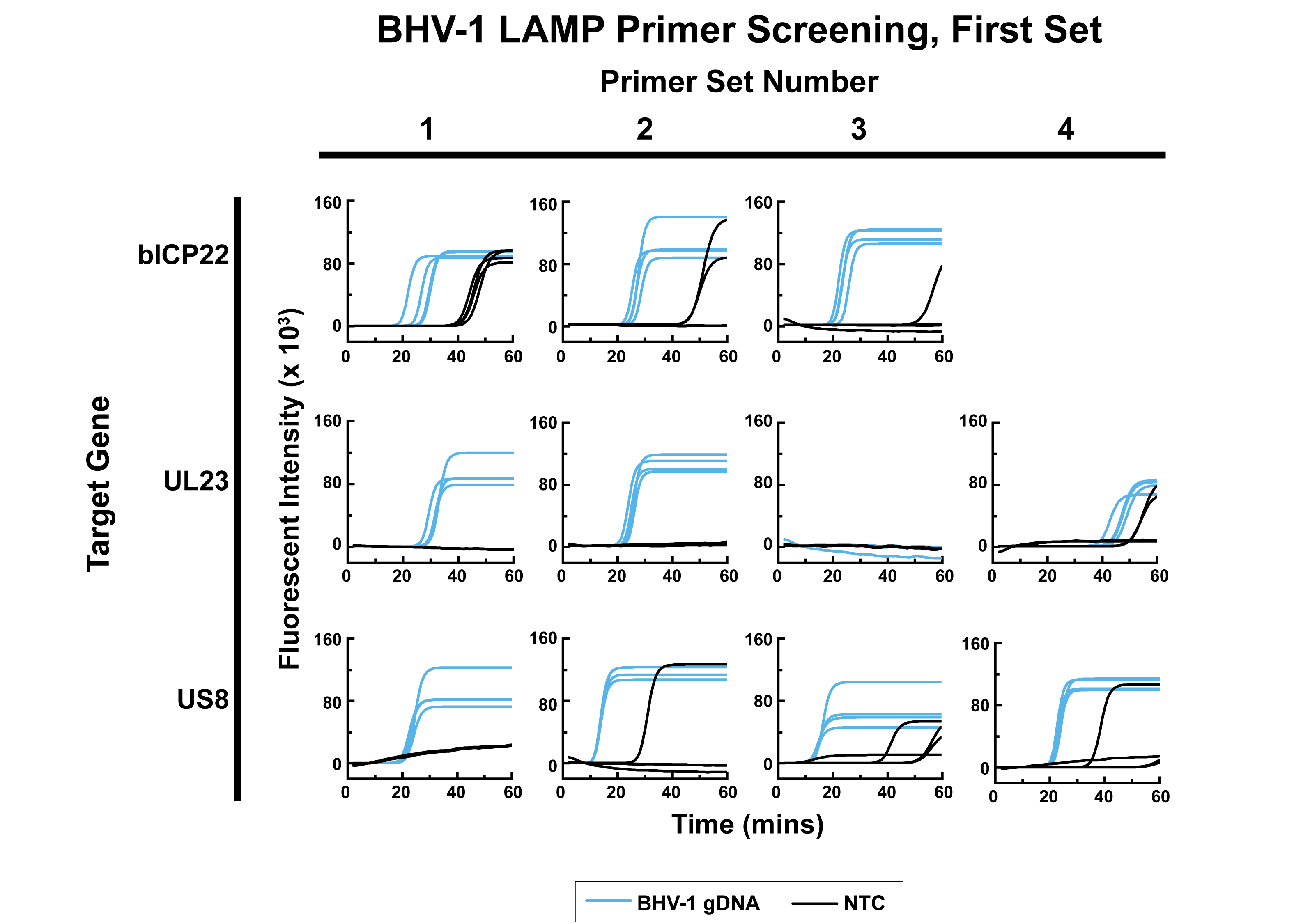

Figure S 9: Fluorometric traces for screening of the first round of LAMP primer sets targeting BHV-1 for primer set IDs between 1 and 4. The target gene is along the vertical axis while the primer set ID for that gene is indicated along the horizontal axis. Reactions consist of 20 µL of LAMP master mix combined with either 5 µL of BHV-1 gDNA extract from virus culture supernatant at 2.0 x 10^4^ copies/µL resulting in a final reaction containing 1.0 x 10 ^5^ copies/reaction (blue lines) or 5 µL of nuclease-free water (black lines). The total reaction volume is 25 µL. Reactions were heated at 65 °C for 60 minutes and the fluorescent intensity was read every 60 seconds.

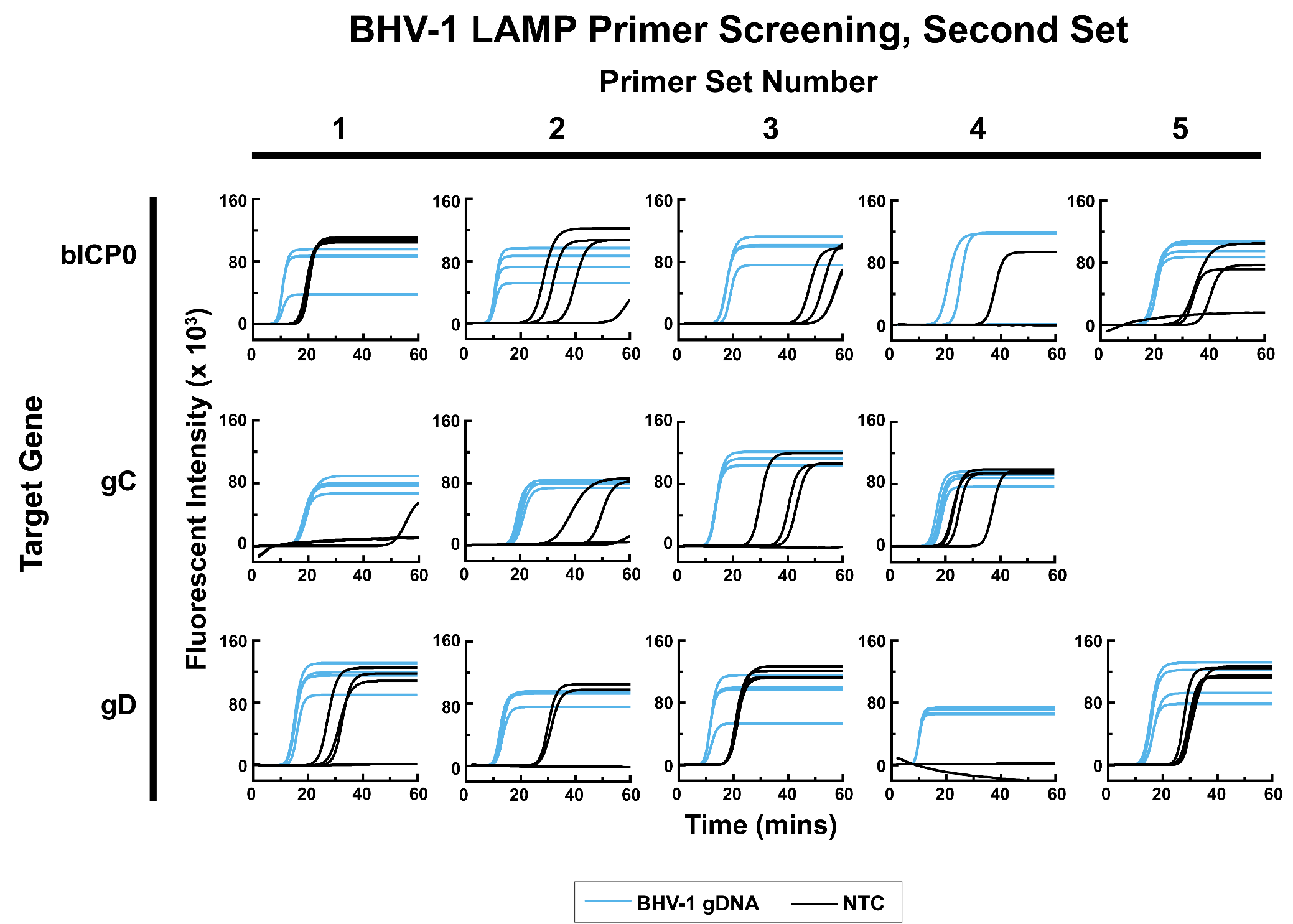

Figure S 10: Fluorometric traces for screening of the second round of LAMP primer sets targeting BHV-1 for primer set IDs between 1 and 5. The target gene is along the vertical axis while the primer set ID for that gene is indicated along the horizontal axis. Reactions consist of 20 µL of LAMP master mix combined with either 5 µL of BHV-1 gDNA extract from virus culture supernatant at 2.0 x 10^4^ copies/µL resulting in a final reaction containing 1.0 x 10 ^5^ copies/reaction (blue lines) or 5 µL of nuclease-free water (black lines). The total reaction volume is 25 µL. Reactions were heated at 65 °C for 60 minutes and the fluorescent intensity was read every 60 seconds.

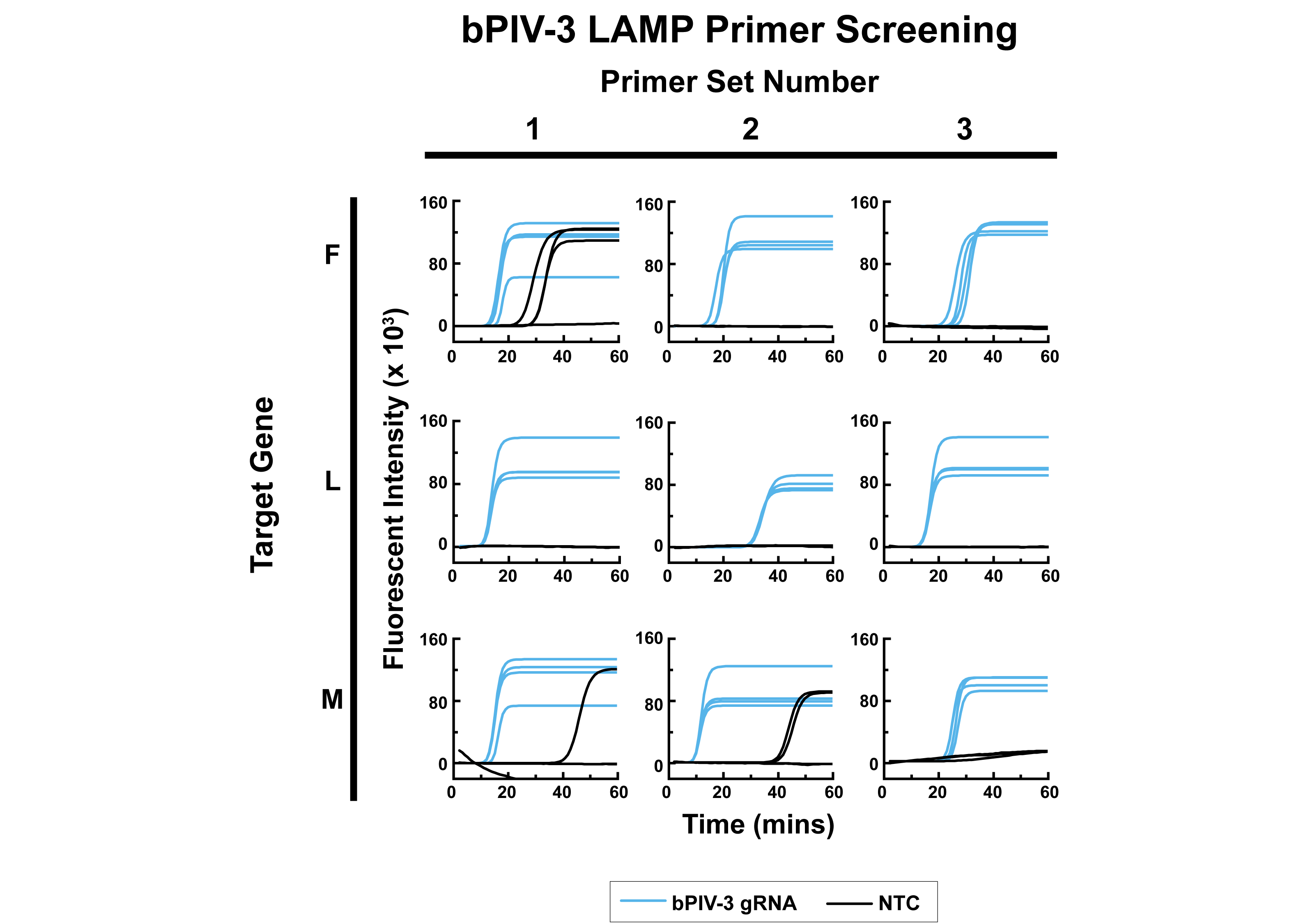

Figure S 11: Fluorometric traces for screening of LAMP primer sets targeting bPIV-3. The target gene is along the vertical axis while the primer set ID for that gene is indicated along the horizontal axis. Reactions consist of 20 µL of LAMP master mix combined with either 5 µL of bPIV-3 gRNA extract from virus culture supernatant at 2.0 x 10^4^ copies/µL resulting in a final reaction containing 1.0 x 10 ^5^ copies/reaction (blue lines) or 5 µL of nuclease-free water (black lines; No Template Control (NTC)). The total reaction volume is 25 µL. Reactions were heated at 65 °C for 60 minutes and the fluorescent intensity was read every 60 seconds.

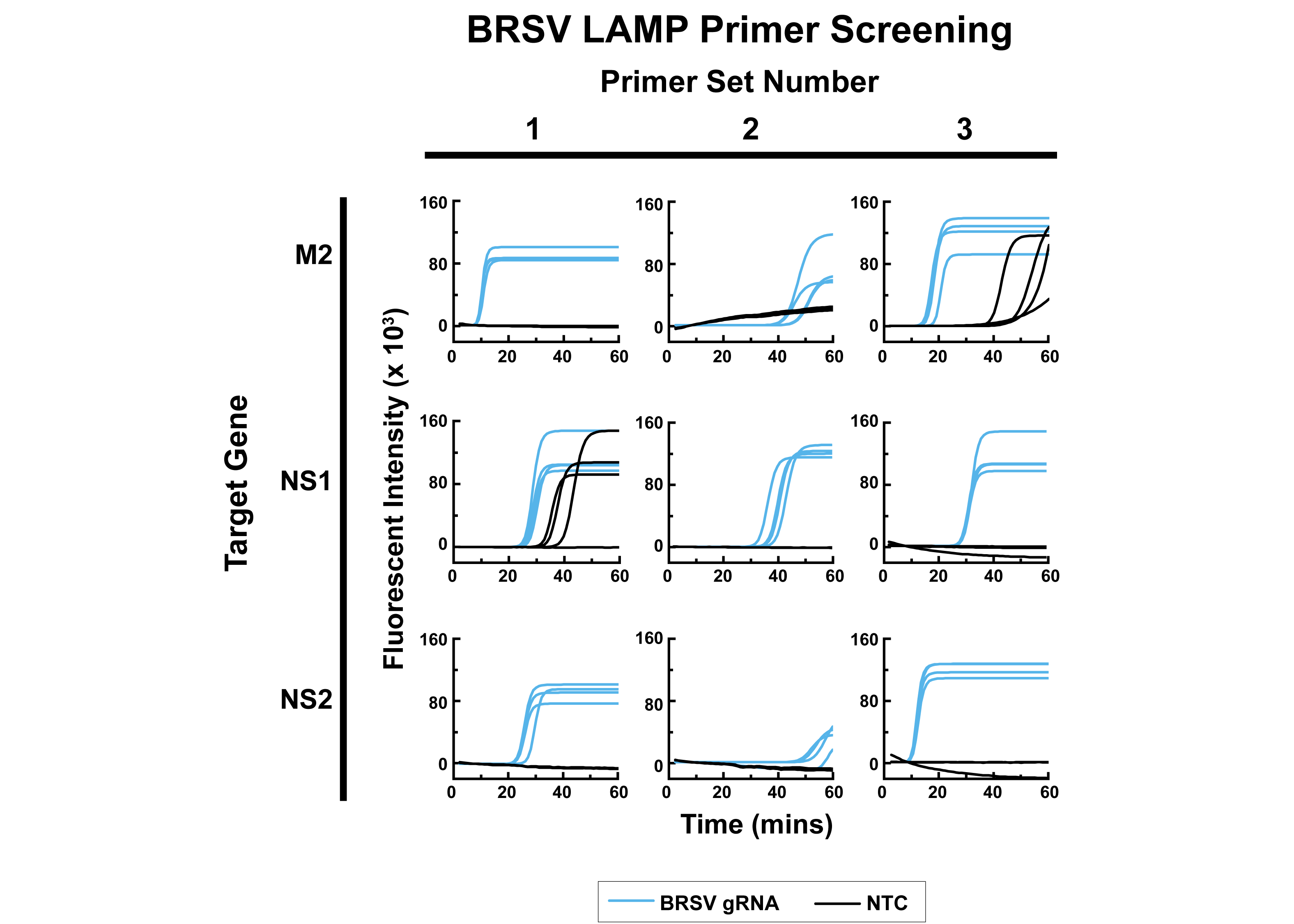

Figure S 12: Fluorometric traces for screening of LAMP primer sets targeting BRSV. The target gene is along the vertical axis while the primer set ID for that gene is indicated along the horizontal axis. Reactions consist of 20 µL of LAMP master mix combined with either 5 µL of BRSV gRNA extract from virus culture supernatant at 2.0 x 10^4^ copies/µL resulting in a final reaction containing 1.0 x 10 ^5^ copies/reaction (blue lines) or 5 µL of nuclease-free water (black lines; No Template Control (NTC)). The total reaction volume is 25 µL. Reactions were heated at 65 °C for 60 minutes and the fluorescent intensity was read every 60 seconds.

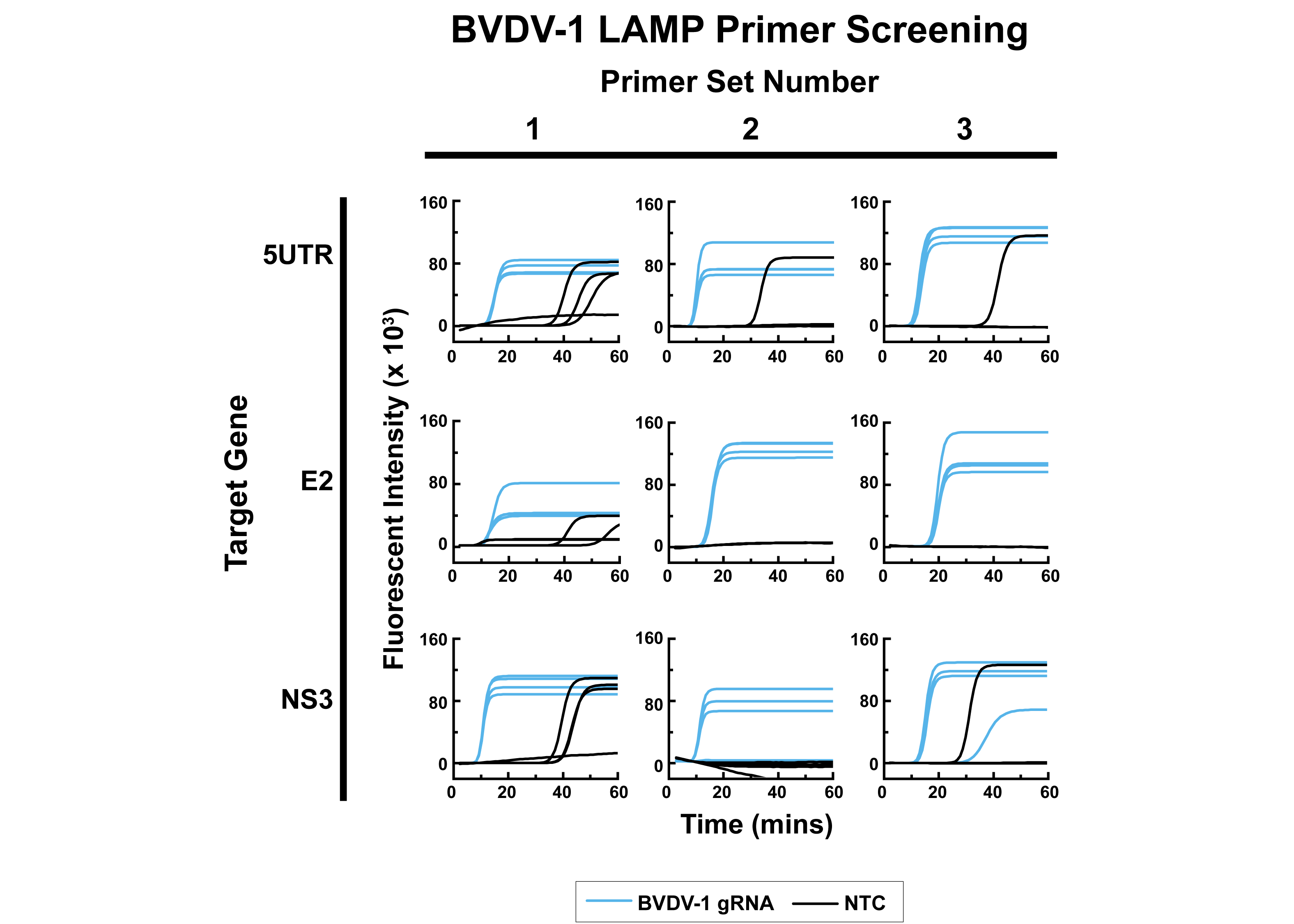

Figure S 13: Fluorometric traces for screening of LAMP primer sets targeting BVDV-1. The target gene is along the vertical axis while the primer set ID for that gene is indicated along the horizontal axis. Reactions consist of 20 µL of LAMP master mix combined with either 5 µL of BVDV-1 gRNA extract from virus culture supernatant at 2.0 x 10^4^ copies/µL resulting in a final reaction containing 1.0 x 10 ^5^ copies/reaction (blue lines) or 5 µL of nuclease-free water (black lines; No Template Control (NTC)). The total reaction volume is 25 µL. Reactions were heated at 65 °C for 60 minutes and the fluorescent intensity was read every 60 seconds.

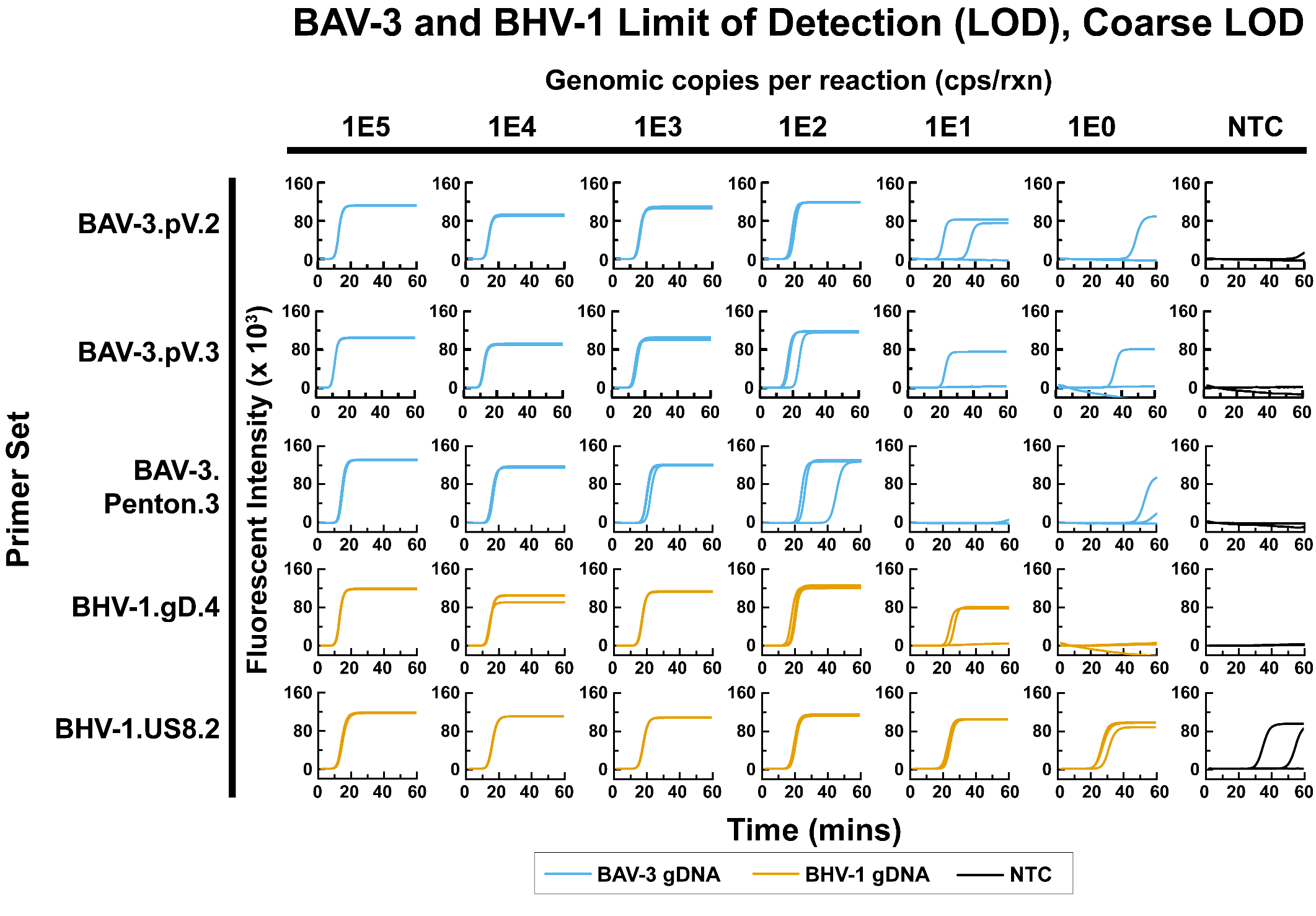

Figure S 14: LAMP fluorometric traces for primer sets advanced to Stage II of primer screening which target DNA viruses (BAV-3 and BHV-1). Reactions have a total volume of 25 µL consisting of 20 µL of LAMP master mix and 5 µL of template diluted in nuclease-free water to 20% of the reported concentration in copies per reaction, such that the final volume in copies per reaction is as reported. The template consisted of genomic DNA extract from BAV-3 and BHV-1 viral culture supernatant for blue and orange plots, respectively. Black plots consisted of no template and instead, 5 µL of nuclease-free water was used as a No Template Control (NTC). Reactions were heated at 65 °C for 60 minutes and the fluorescent intensity was read every 60 seconds.

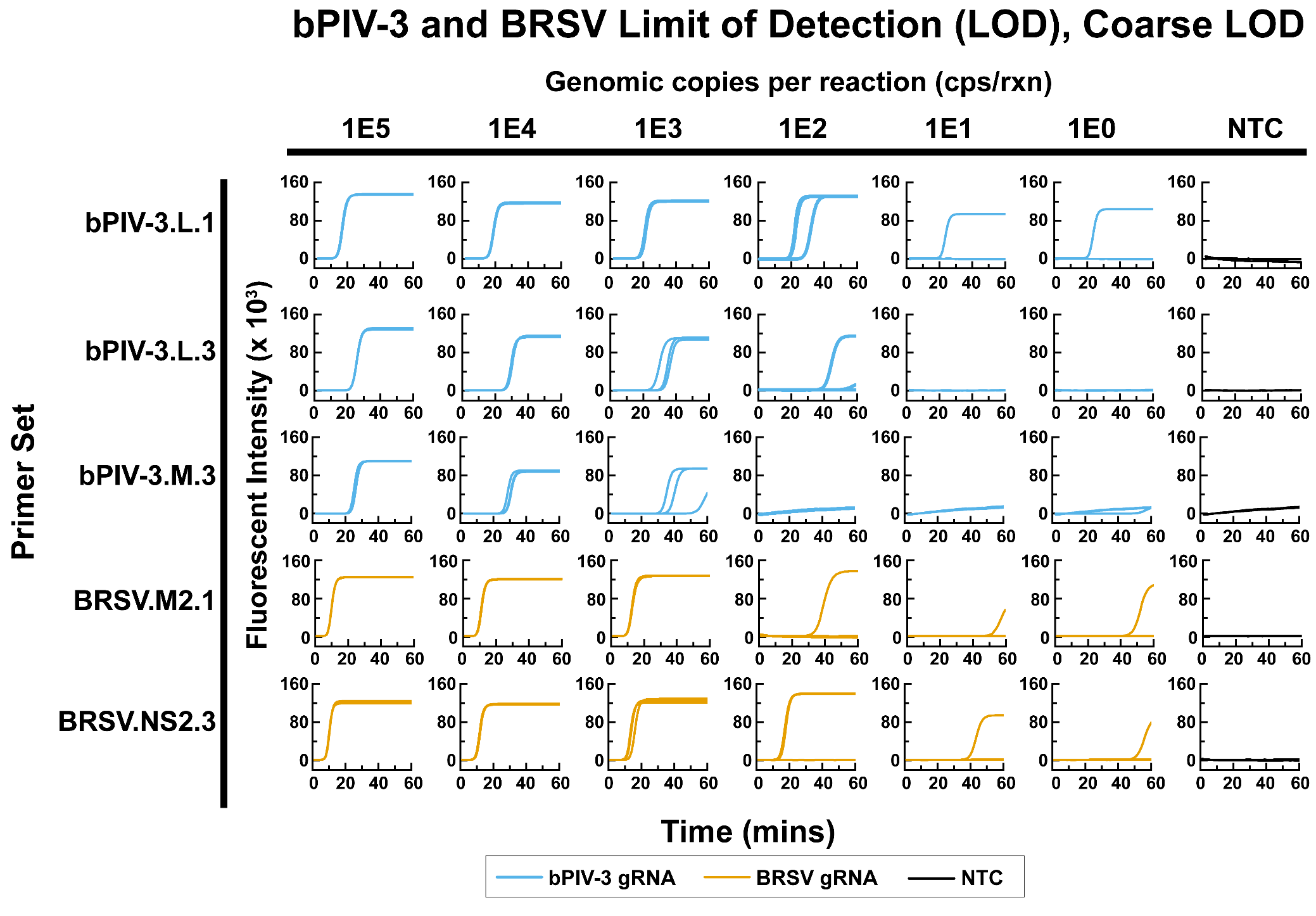

Figure S 15: LAMP fluorometric traces for primer sets advanced to Stage II of primer screening which target RNA viruses (bPIV-3 and BRSV). Reactions have a total volume of 25 µL consisting of 20 µL of LAMP master mix and 5 µL of template diluted in nuclease-free water to 20% of the reported concentration in copies per reaction, such that the final volume in copies per reaction is as reported. The template consisted of genomic RNA extract from bPIV-3 and BRSV viral culture supernatant for blue and orange plots, respectively. Black plots consisted of no template and instead, 5 µL of nuclease-free water was used as a No Template Control (NTC). Reactions were heated at 65 °C for 60 minutes and the fluorescent intensity was read every 60 seconds.

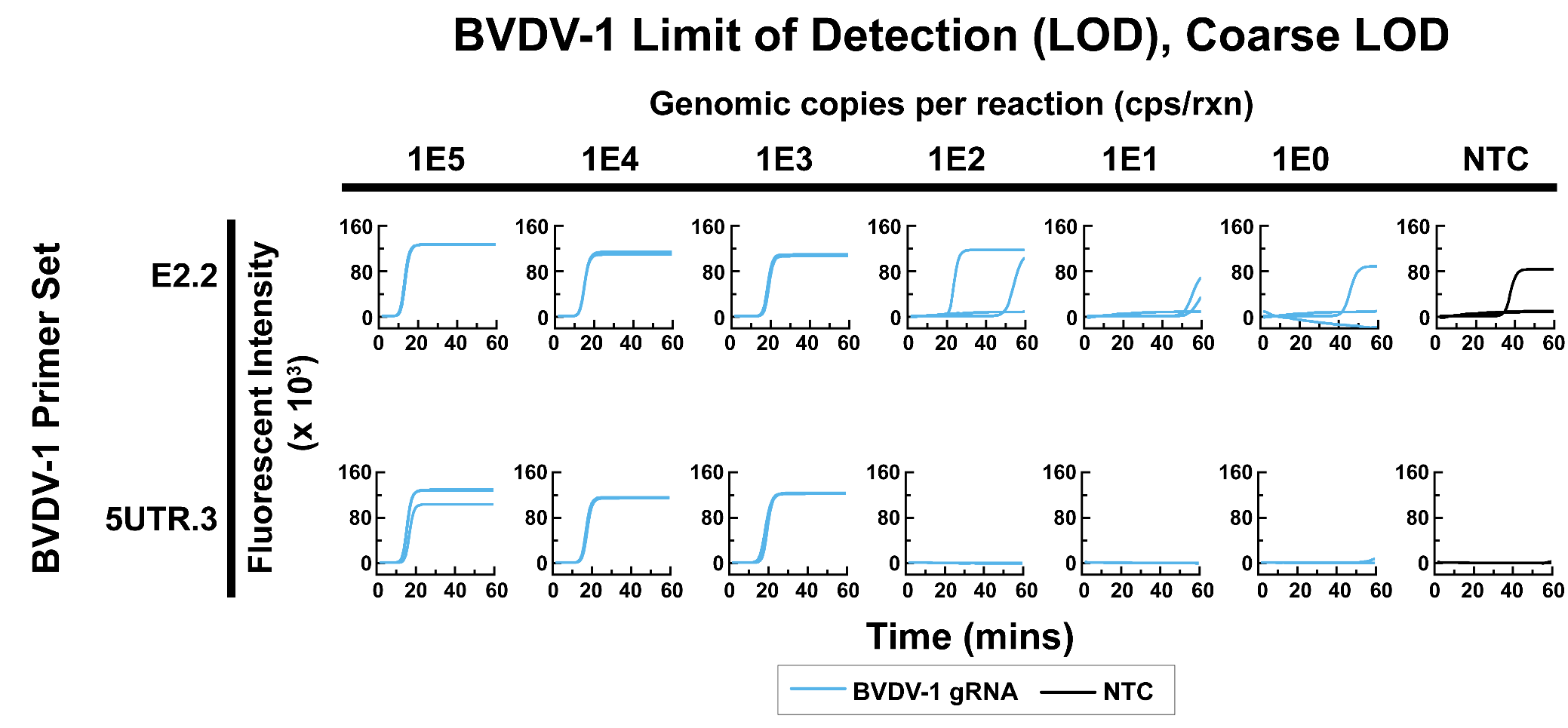

Figure S 16: LAMP fluorometric traces for primer sets advanced to Stage II of primer screening which target BVDV-1. Reactions have a total volume of 25 µL consisting of 20 µL of LAMP master mix and 5 µL of template diluted in nuclease-free water to 20% of the reported concentration in copies per reaction, such that the final volume in copies per reaction is as reported. The template consisted of genomic RNA extract from BVDV-1 viral culture supernatant for blue plots. Black plots consisted of no template and instead, 5 µL of nuclease-free water was used as a No Template Control (NTC). Reactions were heated at 65 °C for 60 minutes and the fluorescent intensity was read every 60 seconds.

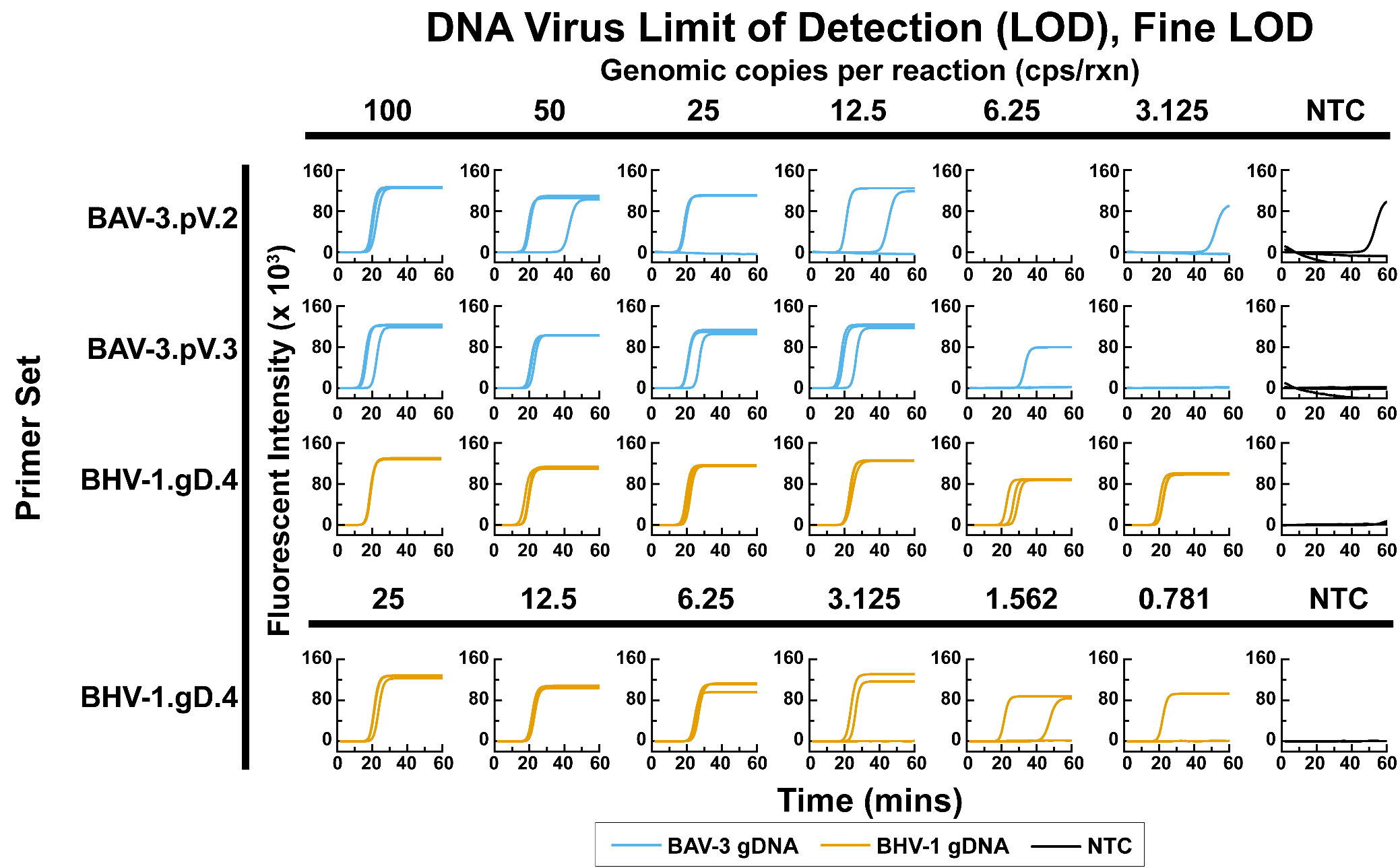

Figure S 17: LOD studies with a fine limit of detection for DNA viruses during this study (BAV-3 and BHV-1). Blue lines indicate reactions wherein 5 µL of whole viral extract from virus culture supernatants was placed into a qLAMP reaction as the template for a final reaction volume of 25 µL. The final template concentration is as indicated above the respective figure. NTC controls had 5 µL of DEPC-treated nuclease-free water placed into the reaction also resulting in a final volume of 25 µL. Reactions were heated at 65 °C for 60 minutes and the fluorescent intensity was read every 60 seconds.

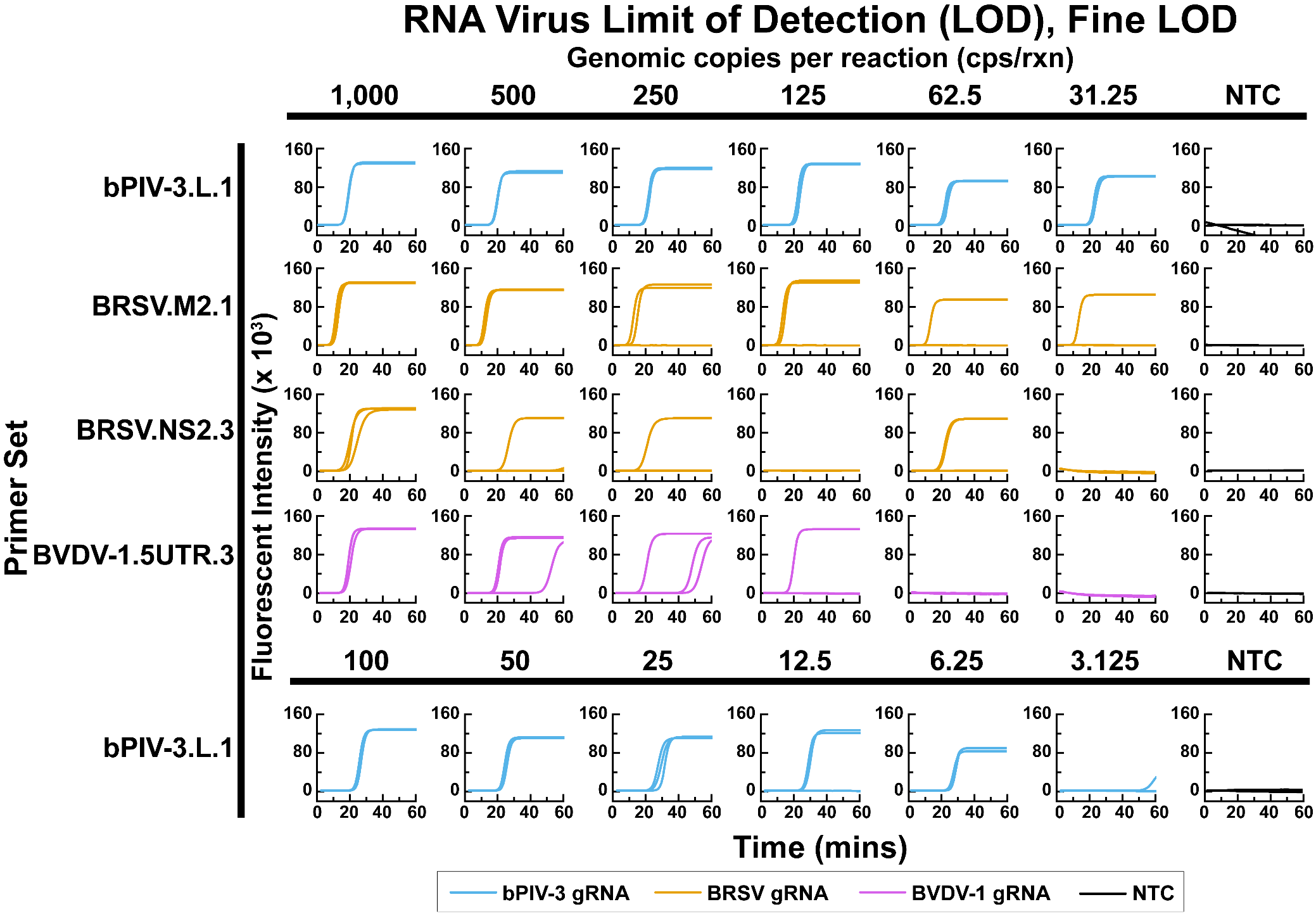

Figure S 18: LOD studies with a fine limit of detection for RNA viruses during this study (bPIV-3, BRSV, and BVDV-1). Blue lines indicate reactions wherein 5 µL of whole viral genomic extract (with carrier RNA) from virus culture supernatants was placed into a qLAMP reaction as the template for a final reaction volume of 25 µL. The final template concentration is as indicated above the respective figure. NTC controls had 5 µL of DEPC-treated nuclease-free water placed into the reaction also resulting in a final volume of 25 µL. Reactions were heated at 65 °C for 60 minutes and the fluorescent intensity was read every 60 seconds.

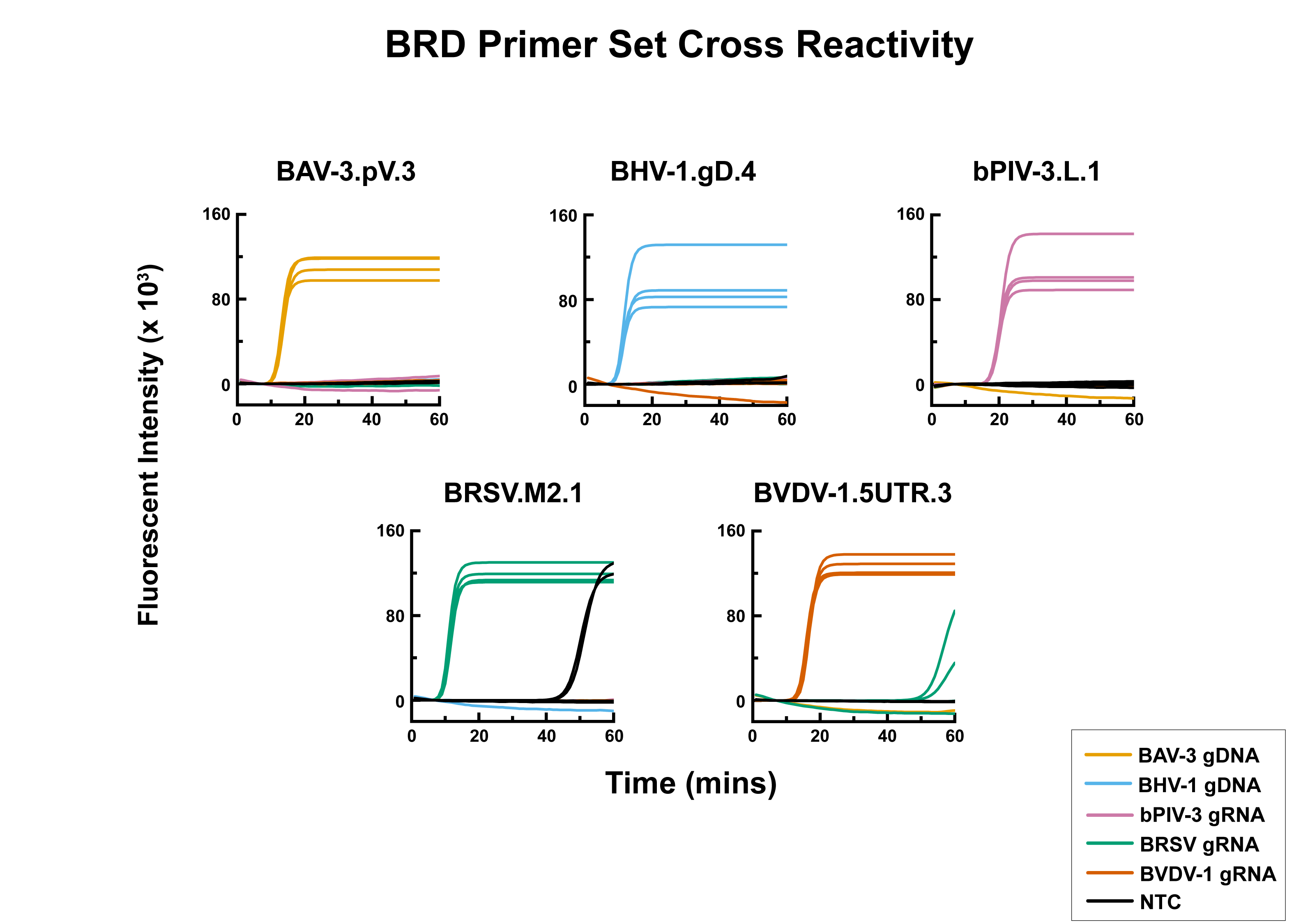

Figure S 19: Fluorometric traces of cross-reactivity tests during Stage IV of primer screening and selection. Reactions consisted of 20 µL of LAMP or RT-LAMP master mix containing the indicated primer set combined with 5 µL of genomic DNA or RNA extract (as indicated) from viral culture supernatant at a concentration of 2.0 x 10^3^ copies per µL for a final reaction concentration of 1.0 x 10^4^ copies per reaction for the case of plots that are colored. Traces plotted in black consisted of 5 µL of DEPC-treated nuclease-free water in place of the template and comprised the No Template Control reactions. All sets consisted of 4 replicates.

### 4. Supplementary Tables

Table S 1: Literature Primers and Probes used for dPCR quantification.

| Primer Set | Primer/Probe Type | Name | Sequence | Ref. |
| --- | --- | --- | --- | --- |
| BAV-3 | Forward | BAV-3.Kishimoto.Hexon.F | ATTACCAGCGTCAACCTCTAC | (Kishimoto et al. 2017) |
|  | Reverse | BAV-3.Kishimoto.Hexon.R | CCGCCGAGAGATAGTCATTAAA |  |
|  | Probe | BAV-3.Kishimoto.Hexon.P | [FAM]-TCCACTTTG-[ZEN]-GAAGCTATGCTCCGC-[IBFQ] |  |
| BHV-1 | Forward | BHV-1.Chandranaik.gB.F | TGCTCGACTACAGCGAGATACAG | (Chandranaik et al. 2013) |
|  | Reverse | BHV-1.Chandranaik.gB.R | CACGCGGTCAATGTCGTAGA |  |
|  | Probe | BHV-1.Chandranaik.gB.P | [FAM]-CCGCAACCA-[ZEN]-GCTGCACGAGC-[IBFQ] |  |
| bPIV-3 | Forward | BPI3ii.Thonur.F | TGATTGGATGTTCGGGAGTGA | (Thonur et al. 2012) |
|  | Reverse | BPI3ii.Thonur.R | AGAATCCTTTCCTCAATCCTGATATACT |  |
|  | Probe | BPI3ii.Thonur.P | [FAM]-TACAATCGA-[ZEN]-GGATCTTGTTCA-[IBFQ] |  |
| BRSV | Forward | BRSV.Boxus.N.F | GCAATGCTGCAGGACTAGGTATAAT | (Boxus et al. 2005) |
|  | Reverse | BRSV.Boxus.N.R | ACACTGTAATTGATGACCCCATTCT |  |
|  | Probe | BRSV.Boxus.N.P | [FAM]-ACCAAGACT-[ZEN]-TGTATGATGCTGCCAAAGCA-[IBFQ] |  |
| BVDV-1 | Forward | BVDV-n.F | GATGCCATGTGGACGAGGGC | (Mari et al. 2016) |
|  | Reverse | BVDV-n.R | CATGTGCCATGTACAGCAGAG |  |
|  | Probe | BVDV-1.P | [FAM]- CAATACAGT-[ZEN]-GGGCCTCTGCAGCA-[IBFQ] |  |

Table S 2: Accession numbers and genome locations for sequences used to design LAMP/RT-LAMP primer sets used in this study.

| Primer Set | Gene Target | GenBank Accession Number | Beginning Index (Inclusive) | Ending Index (Exclusive) |
| --- | --- | --- | --- | --- |
| BAV-3 | E1a | NC_001876.1 | 606 | 1,348 |
|  | E3 |  | 27,377 | 27,742 |
|  | pV |  | 15,068 | 16,300 |
|  | E1b |  | 1,850 | 3,112 |
|  | Hexon |  | 17,809 | 19,204 |
|  | Penton |  | 13,137 | 14,118 |
| BHV-1 | bICP22 | NC_001847.1 | 124,548 | 125,450 |
|  | UL23 |  | 63,260 | 64,339 |
|  | US8 |  | 121,714 | 123,441 |
|  | bICP0 |  | 100,475 | 102,700 |
|  | gC | NC_063268.1 | 16,636 | 18,232 |
|  | gD |  | 118,819 | 120,072 |
| bPIV-3 | F | NC_002161.1 | 4,855 | 6,723 |
|  | L |  | 9,298 | 11,107 |
|  | M |  | 3,703 | 4,851 |
| BRSV | M2 | NC_038272 | 7,514 | 8,473 |
|  | NS1 |  | 46 | 572 |
|  | NS2 |  | 579 | 1,072 |
| BVDV-1 | 5UTR | NC_001461.1 | 1 | 385 |
|  | E2 |  | 2,462 | 3,583 |
|  | NS3 |  | 5,423 | 7,422 |

Table S 3: Average reaction times of selected primer sets using unextracted whole virus from cell culture supernatant spiked into diluted complex field sample (1.8% final reaction concentration of sample).

| Primer Set | LOD | Template Concentration (copies/reaction) | | | | | | NTC |
| --- | --- | --- | --- | --- | --- | --- | --- | --- |
|  |  | **10000** | **5000** | **1000** | **500** | **100** | **50** |  |
| BRSV.M2.1 | 500 | 8.00 | 9.33 | 14.67 | 24.67 | 1 of 3 | 1 of 3 | 0 of 3 |
| BVDV-1.5UTR.3 | 500 | 13.67 | 13.33 | 17.00 | 14.67 | 0 of 3 | 0 of 3 | 0 of 3 |
|  |  | **500** | **100** | **50** | **10** | **5** | **1** |  |
| BAV-3.pV.3 | 50 | 12.00 | 13.33 | 14.67 | 1 of 3 | 0 of 3 | 1 of 3 | 0 of 3 |
| BHV-1.gD.4 | 50 | 14.00 | 14.67 | 16.33 | 2 of 3 | 2 of 3 | 0 of 3 | 0 of 3 |
| bPIV-3.L.1 | 500 | 28.00 | 1 of 3 | 2 of 3 | 0 of 3 | 0 of 3 | 0 of 3 | 0 of 3 |

Table S 4: Expected and measured concentration (using dPCR) of selected primer sets using whole virus in 9% complex field sample (final reaction concentration of 1.8%).

| Primer Set | Expected Concentration (copies/reaction) | Measured Concentration (copies/reaction)  (95% confidence interval) | Primer Set | Expected Concentration (copies/reaction) | Measured Concentration (copies/reaction)  (95% confidence interval) |
| --- | --- | --- | --- | --- | --- |
| BAV-3.pV.3 | 500 | 409 (363, 457) | **bPIV-3.L.1** | 500 | 230 (-54, 515) |
|  | 100 | 69 (59, 81) |  | 100 | 22 (19, 27) |
|  | 50 | 27 (24, 31) |  | 50 | 13 (9, 18) |
|  | 10 | 3 (3, 4) |  | 10 | 2 (-1, 7) |
|  | 5 | 2 (1, 4) |  | 5 | 1 (0, 4) |
|  | 1 | 1 (-1, 4) |  | 1 | 0 (0, 1) |
|  | 0 | 0 (0, 0) |  | 0 | 0 (0, 0) |
| BHV-1.gD.4 | 500 | 1008 (428, 1589) | **BRSV.M2.1** | 10,000 | 6,259 (6,048; 6,471) |
|  | 100 | 260 (129, 392) |  | 5,000 | 3,508 (3,325; 3,691) |
|  | 50 | 102 (77, 129) |  | 1,000 | 829 (791, 867) |
|  | 10 | 15 (2, 29) |  | 500 | 456 (349, 565) |
|  | 5 | 7 (4, 12) |  | 100 | 72 (52, 92) |
|  | 1 | 2 (0, 5) |  | 50 | 51 (34, 69) |
|  | 0 | 0 (0, 0) |  | 0 | 0 (0, 0) |
|  | | | **BVDV-1.5UTR.3** | 10,000 | 8,870 (8,606; 9,135) |
|  |  |  |  | 5,000 | 4,355 (3,987; 4,724) |
|  |  |  |  | 1,000 | 911 (721, 1102) |
|  |  |  |  | 500 | 472 (421, 523) |
|  |  |  |  | 100 | 76 (61, 92) |
|  |  |  |  | 50 | 31 (23, 39) |
|  |  |  |  | 0 | 0 (0, 1) |

Table S 5: dPCR Quantification of BVDV-1 viral culture supernatant dilutions in varying concentrations of complex field samples. Concentrations reported in this table were used in the generation of Figure 1B and were not rounded as were all other dPCR measurements.

|  | Viral Concentration  [copies/reaction (95% confidence interval)] | | | |
| --- | --- | --- | --- | --- |
| Dilution Factor | **Reaction concentration of complex sample** | | | |
|  | **0%** | **1.8%** | **9%** | **18%** |
| 1:100 | 17,294.447  (15,690.717, 18,898.177) | 18,206.288  (14,562.098, 21,850.479) | 14,058.081  (12,156.699, 15,959.463) | 8,117.564  (7,853.340, 8,381.788) |
| 1:1,000 | 2,052.283  (2,004.589, 2,099.978) | 2,012.620  (1,866.261, 2,158.979) | 1,278.706  (1,047.194, 1,510.217) | 860.426  (803.298, 917.553) |
| 1:10,000 | 185.794  (85.922, 285.667) | 182.379  (142.058, 222.700) | 124.249  (92.492, 156.007) | 64.383  (40.029, 88.738) |
| NTC | 2.534 (-5.540, 10.608) | 0.000 (0.000, 0.000) | 0.000 (0.000, 0.000) | 2.577 (-5.768, 10.922) |

Table S 6: Concentration of viruses investigated in this study in the background of collected field samples using dPCR (final sample concentration of 10%). Cells highlighted in vermillion indicate that the background concentration meets or exceeds 2x of the LOD for the selected primer set targeting that virus and is therefore unfit for usage in further testing. Primer sets highlighted in yellow indicate the primer set falls within 1x of the LOD for the selected primer set and therefore should be avoided for further usage in testing, but should not be used for sensitivity and specificity studies. Shaded rows indicate field samples selected for the characterization of selected primer sets in complex samples.

| **Sample ID** | **Viral Background Concentration in 9% Field Sample (cps/µL of 9% Field Sample)** | | | | |
| --- | --- | --- | --- | --- | --- |
|  | **BAV-3** | **BHV-1** | **BPIV-3** | **BRSV** | **BVDV-1** |
| **A406** | 0.00 | 0.00 | 0.00 | 0.00 | 0.00 |
| **C006** | 0.00 | 0.00 | 4.66 | 0.00 | 0.00 |
| **F238** | 3.11 | 1.55 | 0.00 | 0.00 | 0.00 |
| **F280** | 1.50 | 0.00 | 0.00 | 0.00 | 0.00 |
| **G127** | 0.00 | 0.00 | 0.00 | 0.00 | 0.00 |
| **G139** | 1.75 | 0.00 | 1.47 | 0.00 | 0.00 |
| **G143** | 11.55 | 0.00 | 1.57 | 0.00 | 0.00 |
| **H287** | 0.00 | 0.00 | 0.00 | 0.00 | 0.00 |
| **H306** | 3.04 | 3.07 | 1.51 | 1.52 | 0.00 |
| **H323** | 0.00 | 0.00 | 1.51 | 0.00 | 0.00 |
| **H506** | 0.00 | 1.50 | 3.00 | 0.00 | 0.00 |
| **H508** | 7.64 | 0.00 | 0.00 | 0.00 | 0.00 |
| **H510** | 1.53 | 0.00 | 1.53 | 0.00 | 0.00 |
| **J127** | 1.51 | 0.00 | 0.00 | 0.00 | 0.00 |
| **J139** | 0.00 | 0.00 | 1.55 | 0.00 | 0.00 |
| **J228** | 3.11 | 0.00 | 0.00 | 0.00 | 0.00 |
| **J242** | 1.58 | 0.00 | 10.56 | 0.00 | 0.00 |
| **J263** | 1.54 | 0.00 | 0.00 | 0.00 | 0.00 |
| **J276** | 0.00 | 0.00 | 0.00 | 1.54 | 0.00 |
| **J279** | 3.90 | 0.00 | 1.48 | 0.00 | 0.00 |
| **1X LOD (cps/µL)** | 2 | 0.8 | 8.4 | 163 | 47 |
| **LOD (copies/reaction)** | 10 (10, 12) | 4 (3, 6) | 42 (34, 52) | 815 (789, 843) | 235 (194, 276) |

Table S 7: Counts and calculations for Sensitivity and Specificity analyses.

| Primer Set | Multiple of Preliminary LOD | Tt (min) | True Positive | True Negative | False Positive | False Negative | FPR  (%) | FNR (%) | Sensitivity (%) | Specificity (%) | Accuracy (%) | Error (%) |
| --- | --- | --- | --- | --- | --- | --- | --- | --- | --- | --- | --- | --- |
| BAV-3.pV.3 | 2x | 38 | 30 | 30 | 0 | 0 | 0.0 | 0.0 | 100.0 | 100.0 | 100.0 | 0.0 |
|  | 1x | 45 | 29 | 30 | 0 | 1 | 0.0 | 3.3 | 96.7 | 100.0 | 98.3 | 3.3 |
| BHV-1.gD.4 | 2x | 24 | 30 | 30 | 0 | 0 | 0.0 | 0.0 | 100.0 | 100.0 | 100.0 | 0.0 |
|  | 1x | 28 | 30 | 30 | 0 | 0 | 0.0 | 0.0 | 100.0 | 100.0 | 100.0 | 0.0 |
| bPIV-3.L.1 | 2x | 34 | 28 | 30 | 0 | 2 | 0.0 | 6.7 | 93.3 | 100.0 | 96.7 | 6.7 |
|  | 1x | 53 | 25 | 30 | 0 | 5 | 0.0 | 16.7 | 83.3 | 100.0 | 91.7 | 16.7 |
| BRSV.M2.1 | 2x | 40 | 22 | 27 | 3 | 8 | 10.0 | 26.7 | 73.3 | 90.0 | 81.7 | 28.5 |
|  | 1x | 42 | 20 | 25 | 5 | 10 | 16.7 | 33.3 | 66.7 | 83.3 | 75.0 | 37.3 |
| BVDV-1.5UTR.3 | 2x | 29 | 25 | 30 | 0 | 5 | 0.0 | 16.7 | 83.3 | 100.0 | 91.7 | 16.7 |
|  | 1x | 36 | 21 | 30 | 0 | 9 | 0.0 | 30.0 | 70.0 | 100.0 | 85.0 | 30.0 |

Table S 8: Results of primer scoring for candidate primer sets targeting BAV-3 from Stage I of the primer screening and selection process.

| Primer Set | True Positives | Maximum Intensity (RFU) | | Reaction Time (min.) | | False Positives | | | | | Overall Score |
| --- | --- | --- | --- | --- | --- | --- | --- | --- | --- | --- | --- |
|  |  | Average | Std. Dev. | Average | Std. Dev. | Total | Reaction Time (min.) | | | |  |
| BAV-3.pV.2 | 4 | 89680.64 | 11931.96 | 9.00 | 0.00 | 0 | - | - | - | - | 95.22 |
| BAV-3.pV.3 | 4 | 118937.26 | 9845.31 | 11.00 | 0.71 | 1 | 41 | - | - | - | 92.85 |
| BAV-3.Penton.3 | 4 | 90265.55 | 23516.09 | 7.00 | 0.00 | 0 | - | - | - | - | 92.76 |
| BAV-3.E3.2 | 4 | 79054.81 | 7994.47 | 6.25 | 0.43 | 1 | 31 | - | - | - | 91.54 |
| BAV-3.Penton.9 | 4 | 45100.01 | 11589.21 | 6.00 | 0.00 | 1 | 26 | - | - | - | 88.46 |
| BAV-3.E1b.2 | 4 | 46024.79 | 4555.57 | 7.00 | 0.00 | 4 | 43 | 35 | 49 | 52 | 86.92 |
| BAV-3.Hexon.6 | 4 | 50961.24 | 34012.46 | 8.75 | 0.43 | 0 | - | - | - | - | 86.63 |
| BAV-3.Penton.8 | 4 | 44012.99 | 24464.09 | 14.75 | 0.83 | 0 | - | - | - | - | 86.43 |
| BAV-3.Penton.6 | 4 | 59315.96 | 21387.30 | 7.00 | 0.00 | 1 | 23 | - | - | - | 86.17 |
| BAV-3.Penton.2 | 4 | 72567.09 | 36927.75 | 7.50 | 0.50 | 1 | 48 | - | - | - | 86.01 |
| BAV-3.E1a.1 | 4 | 100392.45 | 9506.52 | 7.75 | 0.43 | 4 | 30 | 49 | 18 | 50 | 85.24 |
| BAV-3.Penton.7 | 4 | 77443.44 | 16627.21 | 6.75 | 0.43 | 2 | 25 | 50 | - | - | 85.21 |
| BAV-3.E1b.5 | 4 | 71403.38 | 13240.75 | 7.75 | 0.43 | 2 | 26 | 33 | - | - | 83.69 |
| BAV-3.E3.3 | 4 | 92813.08 | 22823.62 | 7.25 | 0.43 | 4 | 30 | 2 | 47 | 55 | 83.30 |
| BAV-3.pV.1 | 4 | 92833.68 | 9711.46 | 8.00 | 0.00 | 4 | 37 | 3 | 45 | 53 | 83.02 |
| BAV-3.Penton.4 | 4 | 62065.83 | 25322.51 | 6.00 | 0.00 | 2 | 34 | 43 | - | - | 81.66 |
| BAV-3.E1a.2 | 4 | 102472.60 | 24338.56 | 8.00 | 0.00 | 3 | 30 | 35 | 53 | - | 81.14 |
| BAV-3.Penton.1 | 4 | 84677.14 | 8334.66 | 7.00 | 0.00 | 4 | 2 | 2 | 49 | 54 | 80.36 |
| BAV-3.E1a.3 | 4 | 113338.34 | 9935.95 | 8.50 | 0.50 | 3 | 31 | 40 | 45 | - | 78.82 |
| BAV-3.E1b.3 | 4 | 74180.99 | 4498.26 | 7.00 | 0.00 | 4 | 23 | 21 | 36 | 53 | 74.60 |
| BAV-3.Penton.5 | 4 | 60320.78 | 26975.55 | 8.00 | 0.00 | 4 | 5 | 2 | 6 | 6 | 71.66 |
| BAV-3.Penton.10 | 4 | 61220.04 | 30184.04 | 9.00 | 0.71 | 4 | 27 | 48 | 42 | 6 | 70.07 |
| BAV-3.E3.1 | 4 | 95025.33 | 20723.44 | 9.00 | 0.00 | 4 | 23 | 22 | 31 | 53 | 69.59 |
| BAV-3.Hexon.8 | 4 | 30807.53 | 4128.01 | 6.00 | 0.00 | 4 | 20 | 34 | 25 | 23 | 58.98 |
| BAV-3.Hexon.4 | 4 | 36857.97 | 7547.18 | 6.00 | 0.00 | 4 | 26 | 28 | 28 | 35 | 40.47 |
| BAV-3.Hexon.1 | 4 | 99244.55 | 40812.64 | 36.25 | 14.13 | 4 | 25 | 20 | 33 | 50 | 19.13 |
| BAV-3.E1b.4 | 3 | 0.00 | 0.00 | 0.00 | 0.00 | 0 | 0 | 0 | 0 | 0 | 0.00 |
| BAV-3.E1b.1 | 3 | 0.00 | 0.00 | 0.00 | 0.00 | 0 | 0 | 0 | 0 | 0 | 0.00 |
| BAV-3.Hexon.7 | 3 | 0.00 | 0.00 | 0.00 | 0.00 | 0 | 0 | 0 | 0 | 0 | 0.00 |
| BAV-3.Hexon.5 | 1 | 0.00 | 0.00 | 0.00 | 0.00 | 0 | 0 | 0 | 0 | 0 | 0.00 |
| BAV-3.Hexon.3 | 3 | 0.00 | 0.00 | 0.00 | 0.00 | 0 | 0 | 0 | 0 | 0 | 0.00 |
| BAV-3.Hexon.2 | 3 | 0.00 | 0.00 | 0.00 | 0.00 | 0 | 0 | 0 | 0 | 0 | 0.00 |

Table S 9: Results of primer scoring for candidate primer sets targeting BHV-1 from Stage I of the primer screening and selection process.

| Primer Set | True Positives | Maximum Intensity (RFU) | | Reaction Time (min.) | | False Positives | | | | | Overall Score |
| --- | --- | --- | --- | --- | --- | --- | --- | --- | --- | --- | --- |
|  |  | Average | Std. Dev. | Average | Std. Dev. | Total | Reaction Time (min.) | | | |  |
| BHV-1.gD.4 | 4 | 69113.12 | 3870.96 | 6.50 | 0.50 | 0 | - | - | - | - | 92.75 |
| BHV-1.US8.2 | 4 | 117455.05 | 7887.27 | 10.00 | 0.00 | 1 | 27 | - | - | - | 92.63 |
| BHV-1.gC.1 | 4 | 78690.14 | 9194.85 | 14.25 | 0.43 | 1 | 50 | - | - | - | 88.87 |
| BHV-1.UL23.2 | 4 | 106083.72 | 9935.93 | 21.25 | 0.83 | 0 | - | - | - | - | 88.20 |
| BHV-1.US8.4 | 4 | 107528.84 | 7687.88 | 19.50 | 0.50 | 2 | 34 | 35 | - | - | 87.47 |
| BHV-1.bICP22.3 | 4 | 115438.59 | 8979.88 | 19.75 | 1.48 | 1 | 52 | - | - | - | 86.30 |
| BHV-1.gC.2 | 4 | 78781.71 | 4143.96 | 15.75 | 0.83 | 2 | 32 | 45 | - | - | 84.04 |
| BHV-1.gD.2 | 4 | 89837.35 | 9308.00 | 8.75 | 0.43 | 2 | 26 | 26 | - | - | 80.56 |
| BHV-1.US8.1 | 4 | 89984.20 | 22625.90 | 19.50 | 1.12 | 4 | 55 | 55 | 56 | 56 | 78.93 |
| BHV-1.UL23.1 | 4 | 92760.29 | 18133.51 | 27.50 | 1.50 | 0 | - | - | - | - | 78.39 |
| BHV-1.gC.3 | 4 | 110281.50 | 8409.58 | 10.00 | 0.00 | 3 | 26 | 36 | 39 | - | 77.89 |
| BHV-1.bICP0.3 | 4 | 97896.98 | 15586.51 | 13.50 | 0.87 | 4 | 43 | 48 | 52 | 52 | 77.04 |
| BHV-1.US8.3 | 4 | 68721.04 | 25597.81 | 10.75 | 0.83 | 3 | 37 | 51 | 51 | - | 76.87 |
| BHV-1.bICP22.2 | 4 | 104702.33 | 23739.98 | 23.25 | 1.48 | 2 | 46 | 45 | - | - | 73.34 |
| BHV-1.bICP0.5 | 4 | 98657.63 | 9217.24 | 15.25 | 0.43 | 4 | 29 | 29 | 35 | 2 | 71.05 |
| BHV-1.UL23.4 | 4 | 78416.54 | 8332.16 | 42.00 | 2.35 | 2 | 50 | 49 | - | - | 70.73 |
| BHV-1.bICP0.2 | 4 | 76785.77 | 19665.54 | 7.00 | 0.00 | 4 | 24 | 27 | 35 | 53 | 64.94 |
| BHV-1.gD.1 | 4 | 114467.18 | 17281.81 | 11.25 | 0.43 | 3 | 23 | 28 | 27 | - | 63.67 |
| BHV-1.bICP22.1 | 4 | 92596.30 | 3930.78 | 23.25 | 3.27 | 4 | 41 | 40 | 43 | 41 | 57.95 |
| BHV-1.gC.4 | 4 | 88541.60 | 8418.75 | 14.25 | 0.83 | 4 | 19 | 18 | 21 | 33 | 48.93 |
| BHV-1.gD.5 | 4 | 107374.14 | 25237.06 | 11.50 | 0.50 | 4 | 23 | 26 | 25 | 26 | 36.78 |
| BHV-1.bICP0.1 | 4 | 77284.83 | 26426.57 | 7.00 | 0.00 | 4 | 15 | 17 | 16 | 16 | 29.51 |
| BHV-1.gD.3 | 4 | 91512.91 | 26573.75 | 7.50 | 0.50 | 4 | 17 | 17 | 16 | 17 | 24.76 |
| BHV-1.UL23.3 | 0 | 0.00 | 0.00 | 0.00 | 0.00 | 0 | 0 | 0 | 0 | 0 | 0.00 |
| BHV-1.bICP0.4 | 2 | 0.00 | 0.00 | 0.00 | 0.00 | 0 | 0 | 0 | 0 | 0 | 0.00 |

Table S 10: Results of primer scoring for candidate primer sets targeting bPIV-3 from Stage I of the primer screening and selection process.

| Primer Set | True Positives | Maximum Intensity (RFU) | | Reaction Time (min.) | | False Positives | | | | | Overall Score |
| --- | --- | --- | --- | --- | --- | --- | --- | --- | --- | --- | --- |
|  |  | Average | Std. Dev. | Average | Std. Dev. | Total | Reaction Time (min.) | | | |  |
| bPIV-3.L.3 | 4 | 108573.09 | 22114.02 | 12.75 | 0.43 | 0 | - | - | - | - | 86.52 |
| bPIV-3.L.1 | 4 | 104277.41 | 23479.84 | 9.50 | 0.50 | 0 | - | - | - | - | 86.49 |
| bPIV-3.M.3 | 4 | 101689.61 | 8426.01 | 21.75 | 0.83 | 0 | - | - | - | - | 84.96 |
| bPIV-3.F.2 | 4 | 113453.21 | 19152.10 | 15.25 | 1.30 | 0 | - | - | - | - | 81.40 |
| bPIV-3.L.2 | 4 | 81360.09 | 8710.28 | 29.50 | 0.50 | 0 | - | - | - | - | 81.16 |
| bPIV-3.M.2 | 4 | 89830.67 | 23377.59 | 8.00 | 0.00 | 2 | 40 | 41 | - | - | 81.15 |
| bPIV-3.M.1 | 4 | 113034.51 | 26561.32 | 11.50 | 0.87 | 1 | 41 | - | - | - | 79.59 |
| bPIV-3.F.3 | 4 | 125884.94 | 7603.90 | 25.00 | 2.24 | 0 | - | - | - | - | 77.09 |
| bPIV-3.F.1 | 4 | 106217.25 | 30183.07 | 12.75 | 0.83 | 3 | 24 | 29 | 29 | - | 44.02 |

Table S 11: Results of primer scoring for candidate primer sets targeting BRSV from Stage I of the primer screening and selection process.

| Primer Set | True Positives | Maximum Intensity (RFU) | | Reaction Time (min.) | | False Positives | | | | | Overall Score |
| --- | --- | --- | --- | --- | --- | --- | --- | --- | --- | --- | --- |
|  |  | Average | Std. Dev. | Average | Std. Dev. | Total | Reaction Time (min.) | | | |  |
| BRSV.NS2.3 | 4 | 120362.79 | 9133.89 | 8.00 | 0.00 | 0 | - | - | - | - | 98.41 |
| BRSV.M2.1 | 4 | 88978.81 | 7798.80 | 7.00 | 0.00 | 0 | - | - | - | - | 96.04 |
| BRSV.NS1.3 | 4 | 114619.51 | 22977.44 | 27.75 | 0.43 | 0 | - | - | - | - | 83.81 |
| BRSV.NS2.1 | 4 | 91832.36 | 10393.86 | 23.00 | 1.73 | 0 | - | - | - | - | 80.65 |
| BRSV.NS1.2 | 4 | 122604.55 | 6672.15 | 36.00 | 2.55 | 0 | - | - | - | - | 77.11 |
| BRSV.M2.3 | 4 | 119367.86 | 19957.68 | 14.50 | 1.50 | 4 | 38 | 48 | 57 | 57 | 63.80 |
| BRSV.NS1.1 | 4 | 113753.10 | 23204.76 | 25.50 | 0.50 | 3 | 39 | 34 | 32 | - | 47.84 |
| BRSV.M2.2 | 4 | 74081.82 | 29411.26 | 43.75 | 2.28 | 4 | 2 | 2 | 30 | 32 | 24.20 |
| BRSV.NS2.2 | 3 | 0.00 | 0.00 | 0.00 | 0.00 | 0 | 0 | 0 | 0 | 0 | 0.00 |

Table S 12: Results of primer scoring for candidate primer sets targeting BRSV from Stage I of the primer screening and selection process.

| Primer Set | True Positives | Maximum Intensity (RFU) | | Reaction Time (min.) | | False Positives | | | | | Overall Score |
| --- | --- | --- | --- | --- | --- | --- | --- | --- | --- | --- | --- |
|  |  | Average | Std. Dev. | Average | Std. Dev. | Total | Reaction Time (min.) | | | |  |
| BVDV-1.E2.2 | 4 | 126761.51 | 9135.36 | 11.50 | 0.50 | 0 | - | - | - | - | 94.06 |
| BVDV-1.5UTR.3 | 4 | 119311.74 | 9598.67 | 9.50 | 0.50 | 1 | 37 | - | - | - | 91.55 |
| BVDV-1.5UTR.2 | 4 | 80055.69 | 18668.29 | 7.00 | 0.00 | 1 | 29 | - | - | - | 87.45 |
| BVDV-1.E2.1 | 4 | 49902.66 | 19914.84 | 9.25 | 0.43 | 2 | 37 | 51 | - | - | 83.02 |
| BVDV-1.E2.3 | 4 | 113761.33 | 22961.42 | 15.75 | 0.43 | 0 | - | - | - | - | 81.51 |
| BVDV-1.NS3.1 | 4 | 102230.25 | 10745.39 | 7.00 | 0.00 | 3 | 35 | 39 | 39 | - | 63.14 |
| BVDV-1.NS3.3 | 4 | 108281.02 | 26797.18 | 16.75 | 8.81 | 1 | 27 | - | - | - | 57.80 |
| BVDV-1.5UTR.1 | 4 | 74363.51 | 8253.73 | 10.75 | 0.43 | 4 | 36 | 41 | 45 | 43 | 48.96 |
| BVDV-1.NS3.2 | 3 | 0.00 | 0.00 | 0.00 | 0.00 | 0 | 0 | 0 | 0 | 0 | 0.00 |

Table S 13: Average reaction times and LOD of Stage II (“Coarse LOD”) screening of candidate primer sets using extracted whole genome nucleic acids. When an entry states “x of 3” where x is between 0 and 2, it means that not all replicates amplified. Rather, only x replicates out of 3 total replicates amplified. Primer sets that were selected to advance to proceed to Stage III (“Fine LOD”) are highlighted in grey.

| Pathogen | Primer Set | LOD  (copies/reaction) | Template Concentration (copies/reaction) | | | | | | NTC |
| --- | --- | --- | --- | --- | --- | --- | --- | --- | --- |
|  |  |  | **1E5** | **1E4** | **1E3** | **1E2** | **1E1** | **1E0** |  |
|  |  |  | **Average Reaction Time (min) OR number of replicates amplified** | | | | | | |
| BAV-3 | BAV-3.pV.2 | 100 | 9.00 | 10.00 | 12.00 | 15.33 | 2 of 3 | 1 of 3 | 0 of 3 |
|  | BAV-3.pV.3 | 100 | 8.00 | 8.00 | 10.33 | 15.67 | 1 of 3 | 1 of 3 | 0 of 3 |
|  | BAV-3.Penton.3 | 100 | 11.33 | 13.00 | 17.67 | 28.33 | 0 of 3 | 1 of 3 | 0 of 3 |
| BHV-1 | BHV-1.gD.4 | 100 | 9.00 | 10.67 | 13.00 | 15.33 | 2 of 3 | 0 of 3 | 0 of 3 |
|  | BHV-1.US8.2 | - | 10.33 | 12.00 | 14.00 | 16.33 | 19.33 | 23.33 | 2 of 3 |
| BPIV-3 | BPIV-3.L.1 | 100 | 13.00 | 15.00 | 17.67 | 22.33 | 1 of 3 | 1 of 3 | 0 of 3 |
|  | BPIV-3.L.3 | 1000 | 22.00 | 26.00 | 29.33 | 1 of 3 | 0 of 3 | 0 of 3 | 0 of 3 |
|  | BPIV-3.M.3 | 1000 | 21.67 | 26.00 | 40.00 | 0 of 3 | 0 of 3 | 0 of 3 | 0 of 3 |
| BRSV | BRSV.M2.1 | 1000 | 7.00 | 8.00 | 10.00 | 1 of 3 | 1 of 3 | 1 of 3 | 0 of 3 |
|  | BRSV.NS2.3 | 1000 | 7.33 | 9.00 | 11.33 | 2 of 3 | 1 of 3 | 1 of 3 | 0 of 3 |
| BVDV-1 | BVDV-1.5UTR.3 | 1000 | 12.33 | 13.33 | 15.33 | 0 of 3 | 0 of 3 | 0 of 3 | 0 of 3 |
|  | BVDV-1.E2.2 | - | 9.33 | 11.00 | 15.00 | 2 of 3 | 2 of 3 | 1 of 3 | 1 of 3 |

Table S 14: Average reaction times and LOD of Stage III (“Fine LOD”) screening of candidate primer sets using extracted whole genome nucleic acids. When an entry states “x of 3” where x is between 0 and 3, it means that x replicates out of 3 total replicates amplified.

| Primer Set | LOD | Template Concentration (copies/reaction) | | | | | | NTC |
| --- | --- | --- | --- | --- | --- | --- | --- | --- |
|  |  | **1000** | **500** | **250** | **125** | **62.5** | **31.25** |  |
| bPIV-3.L.1 | Indeterminant | 16.67 | 17.33 | 18.67 | 19.67 | 19.67 | 20.33 | 0 of 3 |
| BRSV.M2.1 | 500 | 9.33 | 9.33 | 2 of 3 | 2 of 3 | 1 of 3 | 1 of 3 | 0 of 3 |
| BRSV.NS2.3 | 1000 | 17.00 | 1 of 3 | 1 of 3 | 0 of 3 | 2 of 3 | 0 of 3 | 0 of 3 |
| BVDV-1.5UTR.3 | 250 | 15.33 | 27.00 | 36.00 | 1 of 3 | 0 of 3 | 0 of 3 | 0 of 3 |
|  |  | **100** | **50** | **25** | **12.5** | **6.25** | **3.125** |  |
| BAV-3.pV.2 | Indeterminant | 17.00 | 23.00 | 2 of 3 | 2 of 3 | 0 of 3 | 1 of 3 | 1 of 3 |
| BAV-3.pV.3 | 12.5 | 14.67 | 17.67 | 19.00 | 17.33 | 1 of 3 | 0 of 3 | 0 of 3 |
| BHV-1.gD.4 | Indeterminant | 16.33 | 16.33 | 18 | 20.33 | 23.33 | 18.33 | 0 of 3 |
| bPIV-3.L.1 | 25 | 21.67 | 20.67 | 24.67 | 2 of 3 | 24.67 | 1 of 3 | 0 of 3 |
|  |  | **25** | **12.5** | **6.25** | **3.125** | **1.5625** | **0.78125** |  |
| BHV-1.gD.4 | 6.25 | 18.00 | 18.67 | 21.33 | 2 of 3 | 2 of 3 | 1 of 3 | 0 of 3 |

Table S 15: Expected and measured concentration (using dPCR) of Stage III (“Fine LOD”) screening of candidate primer sets using extracted whole genome nucleic acids for the indicated primer sets. Dilutions and reactions were made using nuclease-free water. Measured concentrations were reported for dPCR that was conducted in parallel with qLAMP/RT-qLAMP reactions. Measured concentrations are rounded down to the nearest whole number; the 95% confidence interval was rounded up for the left-hand tail, and down to the nearest whole number for the right-hand tail.

| Primer Set | Expected Concentration (copies/reaction) | Measured Concentration (copies/reaction)  (95% confidence interval) | Primer Set | Expected Concentration (copies/reaction) | Measured Concentration (copies/reaction)  (95% confidence interval) |
| --- | --- | --- | --- | --- | --- |
| BAV-3.pV.2 /BAV-3.pV.3 | 100 | 76 (49, 104) | **bPIV-3.L.1** | 100 | 199 (182, 216) |
|  | 50 | 39 (28, 51) |  | 50 | 84 (68, 101) |
|  | 25 | 20 (12, 28) |  | 25 | 42 (34, 52) |
|  | 12.5 | 10 (10, 12) |  | 12.5 | 20 (5, 36) |
|  | 6.25 | 5 (1, 10) |  | 6.25 | 7 (7, 8) |
|  | 3.125 | 1 (-2, 4) |  | 3.125 | 4 (3, 6) |
|  | 0 | 0 (0, 1) |  | 0 | 0 (0, 0) |
| BHV-1.gD.4 (Round 1) | 100 | 100 (86, 114) | **BRSV.M2.1** | 1,000 | 1,680 (1,584, 1,777) |
|  | 50 | 61 (50, 72) |  | 500 | 815 (789, 843) |
|  | 25 | 24 (21, 28) |  | 250 | 404 (398, 411) |
|  | 12.5 | 11 (9, 14) |  | 125 | 204 (158, 252) |
|  | 6.25 | 4 (3, 6) |  | 62.5 | 95 (89, 101) |
|  | 3.125 | 5 (-1, 11) |  | 31.25 | 45 (30, 60) |
|  | 0 | 0 (0, 0) |  | 0 | 0 (0, 0) |
| BHV-1.gD.4 (Round 2) | 25 | 20 (10, 31) | **BRSV.NS2.3** | 1,000 | 893 (876, 910) |
|  | 12.5 | 6 (-2, 15) |  | 500 | 294 (236, 353) |
|  | 6.25 | 4 (3, 6) |  | 250 | 135 (85, 185) |
|  | 3.125 | 4 (1, 7) |  | 125 | 59 (44, 75) |
|  | 1.5625 | 0 (0, 1) |  | 62.5 | 22 (12, 32) |
|  | 0.78125 | 0 (0, 1) |  | 31.25 | 9 (5, 15) |
|  | 0 | 0 (0, 0) |  | 0 | 0 (0, 0) |
|  | | | **BVDV-1.5UTR.3** | 1,000 | 1,020 (915, 1,126) |
|  | | |  | 500 | 499 (420, 579) |
|  | | |  | 250 | 235 (194, 276) |
|  | | |  | 125 | 113 (94, 133) |
|  | | |  | 62.5 | 56 (39, 74) |
|  | | |  | 31.25 | 26 (22, 31) |
|  | | |  | 0 | 0 (0, 0) |

Table S 16: *In silico* inclusivity results for primer sets used in this study. Inclusivity percentage is the percentage of viral sequence and primer set alignments with 10 or less mismatches across all primers in the primer sets.

| Primer Set | Number of Sequences | Inclusivity (%) | Primer Set | Number of Sequences | Inclusivity (%) | Primer Set | Number of Sequences | Inclusivity (%) |
| --- | --- | --- | --- | --- | --- | --- | --- | --- |
| BAV-3.E1a.1 | 4 | 100 | **BAV-3.pV.1** | 2 | 100 | **BRSV.NS1.1** | 29 | 100 |
| BAV-3.E1a.2 | 4 | 100 | **BAV-3.pV.2** | 2 | 100 | **BRSV.NS1.2** | 29 | 100 |
| BAV-3.E1a.3 | 4 | 100 | **BAV-3.pV.3** | 2 | 100 | **BRSV.NS1.3** | 29 | 100 |
| BAV-3.E1b.1 | 4 | 75 | **BHV-1.UL23.1** | 64 | 95 | **BRSV.NS2.1** | 30 | 97 |
| BAV-3.E1b.2 | 4 | 100 | **BHV-1.UL23.2** | 64 | 95 | **BRSV.NS2.2** | 30 | 97 |
| BAV-3.E1b.3 | 4 | 75 | **BHV-1.UL23.3** | 64 | 95 | **BRSV.NS2.3** | 30 | 97 |
| BAV-3.E1b.4 | 4 | 75 | **BHV-1.UL23.4** | 64 | 95 | **BVDV-1.5UTR.1** | 5004 | 9 |
| BAV-3.E1b.5 | 4 | 75 | **BHV-1.US8.1** | 73 | 93 | **BVDV-1.5UTR.2** | 5004 | 59 |
| BAV-3.E3.1 | 3 | 100 | **BHV-1.US8.2** | 73 | 92 | **BVDV-1.5UTR.3** | 5004 | 59 |
| BAV-3.E3.2 | 3 | 100 | **BHV-1.US8.3** | 73 | 92 | **BVDV-1.E2.1** | 411 | 7 |
| BAV-3.E3.3 | 3 | 100 | **BHV-1.US8.4** | 73 | 93 | **BVDV-1.E2.2** | 411 | 7 |
| BAV-3.Hexon.1 | 20 | 95 | **BHV-1.bICP0.1** | 70 | 84 | **BVDV-1.E2.3** | 411 | 7 |
| BAV-3.Hexon.2 | 20 | 10 | **BHV-1.bICP0.2** | 70 | 84 | **BVDV-1.NS3.1** | 319 | 10 |
| BAV-3.Hexon.3 | 20 | 10 | **BHV-1.bICP0.3** | 70 | 89 | **BVDV-1.NS3.2** | 319 | 8 |
| BAV-3.Hexon.4 | 20 | 10 | **BHV-1.bICP0.4** | 70 | 84 | **BVDV-1.NS3.3** | 319 | 13 |
| BAV-3.Hexon.5 | 20 | 10 | **BHV-1.bICP0.5** | 70 | 84 | **bPIV-3.F.1** | 107 | 3 |
| BAV-3.Hexon.6 | 20 | 15 | **BHV-1.bICP22.1** | 119 | 95 | **bPIV-3.F.2** | 107 | 4 |
| BAV-3.Hexon.7 | 20 | 15 | **BHV-1.bICP22.2** | 119 | 100 | **bPIV-3.F.3** | 107 | 11 |
| BAV-3.Hexon.8 | 20 | 10 | **BHV-1.bICP22.3** | 119 | 100 | **bPIV-3.L.1** | 57 | 16 |
| BAV-3.Penton.1 | 2 | 100 | **BHV-1.gC.1** | 71 | 96 | **bPIV-3.L.2** | 57 | 14 |
| BAV-3.Penton.2 | 2 | 100 | **BHV-1.gC.2** | 71 | 96 | **bPIV-3.L.3** | 57 | 5 |
| BAV-3.Penton.3 | 2 | 100 | **BHV-1.gC.3** | 71 | 96 | **bPIV-3.M.1** | 70 | 6 |
| BAV-3.Penton.4 | 2 | 100 | **BHV-1.gC.4** | 71 | 96 | **bPIV-3.M.2** | 70 | 6 |
| BAV-3.Penton.5 | 2 | 100 | **BHV-1.gD.1** | 78 | 100 | **bPIV-3.M.3** | 70 | 6 |
| BAV-3.Penton.6 | 2 | 100 | **BHV-1.gD.2** | 78 | 100 |  | | |
| BAV-3.Penton.7 | 2 | 100 | **BHV-1.gD.3** | 78 | 96 |  |  |  |
| BAV-3.Penton.8 | 2 | 100 | **BHV-1.gD.4** | 78 | 100 |  |  |  |
| BAV-3.Penton.9 | 2 | 100 | **BHV-1.gD.5** | 78 | 96 |  |  |  |
| BAV-3.Penton.10 | 2 | 100 | **BRSV.M2.1** | 27 | 93 |  |  |  |
| BAV-3.Penton.11 | 2 | 100 | **BRSV.M2.2** | 27 | 85 |  |  |  |
| BAV-3.Penton.12 | 2 | 100 | **BRSV.M2.3** | 27 | 93 |  |  |  |

### 5. Supplementary References

Boxus M, Letellier C, Kerkhofs P. 2005. Real Time RT-PCR for the detection and quantitation of bovine respiratory syncytial virus. J Virol Methods. 125(2):125–130. https://doi.org/10.1016/j.jviromet.2005.01.008

Chandranaik BM, Rathnamma D, Patil SS, Kovi RC, Dhawan J, Ranganatha S, Isloor S, Renukaprasad C, Prabhudas K. 2013. Development of a Probe Based Real Time PCR Assay for Detection of Bovine Herpes Virus-1 in Semen and Other Clinical Samples. Indian J Virol. 24(1):16–26. https://doi.org/10.1007/s13337-012-0112-1

Kamel M, Davidson JL, Schober JM, Fraley GS, Verma MS. 2025. A paper-based loop-mediated isothermal amplification assay for highly pathogenic avian influenza. Sci Rep. 15(1):12110. https://doi.org/10.1038/s41598-025-95452-6

Kishimoto M, Tsuchiaka S, Rahpaya SS, Hasebe A, Otsu K, Sugimura S, Kobayashi S, Komatsu N, Nagai M, Omatsu T, et al. 2017. Development of a one-run real-time PCR detection system for pathogens associated with bovine respiratory disease complex. J Vet Med Sci. 79(3):517–523. https://doi.org/10.1292/jvms.16-0489

Mari V, Losurdo M, Lucente MS, Lorusso E, Elia G, Martella V, Patruno G, Buonavoglia D, Decaro N. 2016. Multiplex real-time RT-PCR assay for bovine viral diarrhea virus type 1, type 2 and HoBi-like pestivirus. J Virol Methods. 229:1–7. https://doi.org/10.1016/j.jviromet.2015.12.003

Nixon G, Garson JA, Grant P, Nastouli E, Foy CA, Huggett JF. 2014. Comparative Study of Sensitivity, Linearity, and Resistance to Inhibition of Digital and Nondigital Polymerase Chain Reaction and Loop Mediated Isothermal Amplification Assays for Quantification of Human Cytomegalovirus. Anal Chem. 86(9):4387–4394. https://doi.org/10.1021/ac500208w

Pavšič J, Žel J, Milavec M. 2016. Digital PCR for direct quantification of viruses without DNA extraction. Anal Bioanal Chem. 408(1):67–75. https://doi.org/10.1007/s00216-015-9109-0

QIAGEN. 2023. QIAcuity® Application Guide Version 2 [Internet]. [accessed 2025 Mar 31]. https://www.qiagen.com/us/resources/resourcedetail?id=5d19083d-fa10-4ed2-88a0-2953d9947e0c

Thonur L, Maley M, Gilray J, Crook T, Laming E, Turnbull D, Nath M, Willoughby K. 2012. One-step multiplex real time RT-PCR for the detection of bovine respiratory syncytial virus, bovine herpesvirus 1 and bovine parainfluenza virus 3. BMC Vet Res. 8(1):37. https://doi.org/10.1186/1746-6148-8-37
