## Supplementary File 3 for "Detection of five viruses commonly implicated with Bovine Respiratory Disease using loop-mediated isothermal amplification": LICENSE

Creative Commons Legal Code

CC0 1.0 Universal

    CREATIVE COMMONS CORPORATION IS NOT A LAW FIRM AND DOES NOT PROVIDE
    LEGAL SERVICES. DISTRIBUTION OF THIS DOCUMENT DOES NOT CREATE AN
    ATTORNEY-CLIENT RELATIONSHIP. CREATIVE COMMONS PROVIDES THIS
    INFORMATION ON AN "AS-IS" BASIS. CREATIVE COMMONS MAKES NO WARRANTIES
    REGARDING THE USE OF THIS DOCUMENT OR THE INFORMATION OR WORKS
    PROVIDED HEREUNDER, AND DISCLAIMS LIABILITY FOR DAMAGES RESULTING FROM
    THE USE OF THIS DOCUMENT OR THE INFORMATION OR WORKS PROVIDED
    HEREUNDER.

Statement of Purpose

The laws of most jurisdictions throughout the world automatically confer
exclusive Copyright and Related Rights (defined below) upon the creator
and subsequent owner(s) (each and all, an "owner") of an original work of
authorship and/or a database (each, a "Work").

Certain owners wish to permanently relinquish those rights to a Work for
the purpose of contributing to a commons of creative, cultural and
scientific works ("Commons") that the public can reliably and without fear
of later claims of infringement build upon, modify, incorporate in other
works, reuse and redistribute as freely as possible in any form whatsoever
and for any purposes, including without limitation commercial purposes.
These owners may contribute to the Commons to promote the ideal of a free
culture and the further production of creative, cultural and scientific
works, or to gain reputation or greater distribution for their Work in
part through the use and efforts of others.

For these and/or other purposes and motivations, and without any
expectation of additional consideration or compensation, the person
associating CC0 with a Work (the "Affirmer"), to the extent that he or she
is an owner of Copyright and Related Rights in the Work, voluntarily
elects to apply CC0 to the Work and publicly distribute the Work under its
terms, with knowledge of his or her Copyright and Related Rights in the
Work and the meaning and intended legal effect of CC0 on those rights.

1. Copyright and Related Rights. A Work made available under CC0 may be
protected by copyright and related or neighboring rights ("Copyright and
Related Rights"). Copyright and Related Rights include, but are not
limited to, the following:

  i. the right to reproduce, adapt, distribute, perform, display,
     communicate, and translate a Work;
 ii. moral rights retained by the original author(s) and/or performer(s);
iii. publicity and privacy rights pertaining to a person's image or
     likeness depicted in a Work;
 iv. rights protecting against unfair competition in regards to a Work,
     subject to the limitations in paragraph 4(a), below;
  v. rights protecting the extraction, dissemination, use and reuse of data
     in a Work;
 vi. database rights (such as those arising under Directive 96/9/EC of the
     European Parliament and of the Council of 11 March 1996 on the legal
     protection of databases, and under any national implementation
     thereof, including any amended or successor version of such
     directive); and
vii. other similar, equivalent or corresponding rights throughout the
     world based on applicable law or treaty, and any national
     implementations thereof.

2. Waiver. To the greatest extent permitted by, but not in contravention
of, applicable law, Affirmer hereby overtly, fully, permanently,
irrevocably and unconditionally waives, abandons, and surrenders all of
Affirmer's Copyright and Related Rights and associated claims and causes
of action, whether now known or unknown (including existing as well as
future claims and causes of action), in the Work (i) in all territories
worldwide, (ii) for the maximum duration provided by applicable law or
treaty (including future time extensions), (iii) in any current or future
medium and for any number of copies, and (iv) for any purpose whatsoever,
including without limitation commercial, advertising or promotional
purposes (the "Waiver"). Affirmer makes the Waiver for the benefit of each
member of the public at large and to the detriment of Affirmer's heirs and
successors, fully intending that such Waiver shall not be subject to
revocation, rescission, cancellation, termination, or any other legal or
equitable action to disrupt the quiet enjoyment of the Work by the public
as contemplated by Affirmer's express Statement of Purpose.

3. Public License Fallback. Should any part of the Waiver for any reason
be judged legally invalid or ineffective under applicable law, then the
Waiver shall be preserved to the maximum extent permitted taking into
account Affirmer's express Statement of Purpose. In addition, to the
extent the Waiver is so judged Affirmer hereby grants to each affected
person a royalty-free, non transferable, non sublicensable, non exclusive,
irrevocable and unconditional license to exercise Affirmer's Copyright and
Related Rights in the Work (i) in all territories worldwide, (ii) for the
maximum duration provided by applicable law or treaty (including future
time extensions), (iii) in any current or future medium and for any number
of copies, and (iv) for any purpose whatsoever, including without
limitation commercial, advertising or promotional purposes (the
"License"). The License shall be deemed effective as of the date CC0 was
applied by Affirmer to the Work. Should any part of the License for any
reason be judged legally invalid or ineffective under applicable law, such
partial invalidity or ineffectiveness shall not invalidate the remainder
of the License, and in such case Affirmer hereby affirms that he or she
will not (i) exercise any of his or her remaining Copyright and Related
Rights in the Work or (ii) assert any associated claims and causes of
action with respect to the Work, in either case contrary to Affirmer's
express Statement of Purpose.

4. Limitations and Disclaimers.

 a. No trademark or patent rights held by Affirmer are waived, abandoned,
    surrendered, licensed or otherwise affected by this document.
 b. Affirmer offers the Work as-is and makes no representations or
    warranties of any kind concerning the Work, express, implied,
    statutory or otherwise, including without limitation warranties of
    title, merchantability, fitness for a particular purpose, non
    infringement, or the absence of latent or other defects, accuracy, or
    the present or absence of errors, whether or not discoverable, all to
    the greatest extent permissible under applicable law.
 c. Affirmer disclaims responsibility for clearing rights of other persons
    that may apply to the Work or any use thereof, including without
    limitation any person's Copyright and Related Rights in the Work.
    Further, Affirmer disclaims responsibility for obtaining any necessary
    consents, permissions or other rights required for any use of the
    Work.
 d. Affirmer understands and acknowledges that Creative Commons is not a
    party to this document and has no duty or obligation with respect to
    this CC0 or use of the Work.
