## Supplementary File 3 for "Detection of five viruses commonly implicated with Bovine Respiratory Disease using loop-mediated isothermal amplification": PrimerAnalysis.ipynb

```
# %% [markdown]
# Execution of Primer Scoring and data analysis code for BRD publication

This workbook is intended to describe the usage of the `PrimerScore` functionality used during data analysis for the publication of *"Detection of five viruses commonly implicated with Bovine Respiratory Disease using loop-mediated isothermal amplification"*

## Setup and initialization
Import `PrimerScore` and `matplotlib` dependencies for this module

## License

Copyright (C) 2025 Purdue Research Foundation. This work is protected under GNU Affero General Public License Version 3 with Commons Clause. Please see the `LICENSE` file for more information.

## Import

# %% [code]
from matplotlib import pyplot as plt

import PrimerScore

# %% [markdown]
Set file and input/output directory paths
 
*Modify file paths and feel free to use patterns for common data types*

# %% [code]
# Set list of virus names
virusName = ['BAV-3', 'BHV-1', 'bPIV-3', 'BRSV', 'BVDV-1']

# Set path for primer scoring data
screeningPathInputPattern = ""
coarseLODInputPath = ""
fineLODGenomicInputPathPattern = ""
fineLODVirusInputPath = ""
sensSpecInputPath = ""


# Set path for primer scoring output
screeningPathOutputPattern = ""
coarseLODOutputPath = ""
fineLODGenomicOutputPathPattern = ""
fineLODVirusOutputPath = ""
sensSpecOutputPath = ""

# Define number of fine LOD rounds
numFineLODRounds = 2

# Initialize PrimerScore class
PrimerScore.initialize([5,10,15,10,60], [3000, 200, 0.9, 0.2], 4, set_threshold_perc=0.2)

# %% [markdown]
## Stage 1: Primer Scoring

Execute the `scorePrimers` function of the `PrimerScore` module to score and rank primers for each virus.

# %% [code]
# Loop through each virus in our list of virus names
for virus in virusName:
    # Score primers for each virus and output results to csv file using output filename pattern
    PrimerScore.scorePrimers(screeningPathInputPattern.format(virus), screeningPathOutputPattern.format(virus))

# %% [markdown]
## Stage 2: Coarse LOD Analysis

Execute the `scoreLOD` function of the `PrimerScore` module to determine and rank the LOD for each primer set that has progressed to Stage 2

# %% [code]
# Run coarse LOD primer screening. Result is a dataframe that is additionally written to an output csv as defined by the output path
coarseLOD_df = PrimerScore.score_LOD(coarseLODInputPath, coarseLODOutputPath)

# %% [markdown]
# Stage 3: Fine LOD Analysis

Execute the `scoreLOD` function of the `PrimerScore` module to determine and rank the fine LOD for each primer set that has progressed to Stage 3. The fine LOD will be broken into successive rounds depending on if the fine LOD is below the minimum dilution level based on the coarse LOD of primer sets progressing to stage 3.

# %% [code]
fineLODGenomicResults_dfList = []
for fineLODRound in range(1, numFineLODRounds+1):
    fineLODGenomicResults_dfList.append(PrimerScore.score_LOD(fineLODGenomicInputPathPattern.format(fineLODRound), fineLODGenomicOutputPathPattern.format(fineLODRound)))

fineLODVirusResutls_df = PrimerScore.score_LOD(fineLODVirusInputPath, fineLODVirusOutputPath)

# %% [markdown]
## Sensitivity and Specificity Analysis

Carry out sensitivity and specificity analysis for all primer sets tested.

# %% [code]
[optimalResults, allResults] = PrimerScore.executeSensSpecAnalysis(sensSpecInputPath, sensSpecOutputPath)
optimalResults

# %% [code]
# Generate list of color blind colors
color_list = [
    ###########################################################################################
    # Colors from Points of View: Color Blindness (Nature Methods, 8, 441 (2011)) by B. Wong  #
    # https://www.nature.com/articles/nmeth.1618                                              #
    # RGB values extracted using ChatGPT v4o (https://chat.openai.com/) and verified by eye   #
    ###########################################################################################
    (230, 159, 0),      # Orange
    (86, 180, 233),     # Sky blue
    (0, 158, 115),      # Bluish green
    (240, 228, 66),     # Yellow
    (0, 114, 178),      # Blue
    (213, 94, 0),       # Vermillion
    (204, 121, 167),    # Reddish purple
    (0, 0, 0),          # Black
]

# Create figure counter to start a new figure for each primer set
figCount = 1

# Loop through each primer set and plot ROC curves for each concentration
for pSet in allResults['Primer Set'].unique():
    # Initialize new figure for this primer set
    plt.figure(figCount)

    # Create color index counter for each concentration level being tested
    colorIdx = 0

    # Loop through each concentration level and plot ROC curve
    for conc in allResults[allResults['Primer Set'] == pSet]['Concentration'].unique():

        # Plot ROC curve using the a color blind list. If number of concentrations levels exceeds 
        # number of colors in list, color will repeat
        plt.plot(allResults[(allResults['Primer Set'] == pSet) & (allResults['Concentration'] == conc)]['FPR'], 
                 allResults[(allResults['Primer Set'] == pSet) & (allResults['Concentration'] == conc)]['Sensitivity'], 
                linestyle='-', marker='o', label=conc, color=tuple(c / 255 for c in color_list[colorIdx % len(color_list)]))
        colorIdx += 1

    # Format plot and add cutoff line of 95% sensitivity
    plt.legend()
    plt.xlabel('FPR')
    plt.ylabel('Sensitivity')
    plt.title(f"ROC Curve for {pSet}")
    plt.plot([0,1], [0.95, 0.95], linestyle='dashed')
    plt.xlim(0,1)
    plt.ylim(0,1)

    # Increment figure counter for next primer set figure
    figCount += 1

# %% [code]
```
