## Supplementary File 3 for "Detection of five viruses commonly implicated with Bovine Respiratory Disease using loop-mediated isothermal amplification": PrimerScore.py

```
#!/usr/bin/env python

'''PrimerAnalysis.py is a python module for analyzing LAMP primer amplification data'''

__author__ = 'Josiah Levi Davidson'
__contact__ = 'Mohit S. Verma'
__email__ = ''
__version__ = '1.1.0'
__status__ = 'Development'
__copyright__ = "Copyright (C) 2024 Purdue Research Foundation"
__license__ = "GNU Affero General Public License Version 3 with Commons Clause"

#### BEGIN IMPORTS
import math
import pandas as pd
import numpy as np
from os import path
from matplotlib import pyplot as plt

# BEGIN GLOBAL VARIABLES
weights = {}
thresholds = []
replicates = 0
primerSetName = ""
rxnTypeDesignation = ""

# BEGIN CONSTANTS
MAX_INST_SIGNAL = 140000
threshold_perc = 0.1


# Initialization method to set weights and thresholds.
# Aruments
#   set_weights := Array of length 5 which contains weights in the following order:
#       Average Maximum Intensity
#       Standard Deviation of Maximum Intensity
#       Standard Deviation of Reaction Time
#       Average Reaction Time
#       False Positive
#   set_thresholds := Array of threshold values to determine exponential phase beginning,
#                       plateau phase ending, positive reaction calculation threshold,
#                       and false positive threshold, respectively
#   set_replicates := Number of replicates. Must be the same for all primer sets.
#   set_instrument_signal := Maximum fluorescent intensity of instrument. Default is 140000
#   set_threshold_perc := Threshold percentage of maximum fluorescent intensity of instrument. Default is 0.1
def initialize(set_weights = [5, 5, 10, 20, 60], set_thresholds = [3000, 200, 0.9, 0.2], set_replicates = 4, set_instrument_signal = 140000, set_threshold_perc = 0.1):
    if not (len(set_weights) == 5):
        raise ValueError('Weights do not contain 5 values. Refer to readme for more information.')
    if not (len(set_thresholds) == 4):
        raise ValueError('thresholds do not contain 2 values. Refer to readme for moe information.')
    if not (set_replicates >= 3):
        raise ValueError('Number of replicates is not greater than or equal to 3.')
    
    global weights, thresholds, replicates, primerSetName, rxnTypeDesignation, MAX_INST_SIGNAL, threshold_perc
    weights = {'Intensity_Avg': set_weights[0], 'Intensity_StdDev': set_weights[1], 'RxnTime_Std':set_weights[2],
               'RxnTime_Avg': set_weights[3], 'False_Positives':set_weights[4]}
    thresholds = set_thresholds
    replicates = set_replicates
    MAX_INST_SIGNAL = set_instrument_signal
    threshold_perc = set_threshold_perc

# Main primer scoring method. Call to generate output
# Arguments:
#   primerData := Path to input file containing primer data formatted correctly. See readme for more details.
#   output := Output file name containing .xlsx
def scorePrimers(primerData, output):
    if not (len(thresholds) == 4):
        raise BaseException('Not initialized. Initialize scoring')
    if not path.exists(primerData):
        raise ValueError('Primer Data file not found.')

    if not (len(output) > 0):
        raise ValueError('Ouput file name not properly defined.')

    getPrimerSummary(primerData).to_csv(output, encoding='utf-8')


# Function to calculate average maximum intensity across the positive replicates of a single primer set
# Arguments:
#   name := String, name of primer set to calculate maximum intensity for
#   df := DataFrame, dataframe containing all primer set amplification data
def calc_max_avg(name, df):
    df = df[df['Primer']==name]
    pos_rxns = df[df['RxnType']=='+'].drop(['Primer', 'RxnType','ObservedType', 'RxnTime', 'RxnPenalty'], axis=1)
    return pos_rxns.max(axis=1).mean()

# Function to calculate the standard deviation of the maximum intensity across the positive replicates of a single primer set
# Arguments:
#   name := String, name of primer set to calculate standard deviation for
#   df := DataFrame, dataframe containing all primer set amplification data
def calc_max_std(name, df):
    df = df[df['Primer']==name]
    pos_rxns = df[df['RxnType']=='+'].drop(['Primer', 'RxnType','ObservedType', 'RxnTime', 'RxnPenalty'], axis=1)
    return pos_rxns.max(axis=1).std()

# Function to calculate average reaction time for a single primer set.
# Arguments:
#   name := String, name of primer set to calculate average reaction time
#   df := DataFrame, dataframe containing all primer set amplification data
def calc_rxn_time_avg(name, df):
    df = df[df['Primer']==name]
    pos_rxns = df[df['RxnType']=='+']
    return pos_rxns['RxnTime'].mean()

# Function to calculate standard deviation of the reaction time across the positive replicates of a single primer set
# Arguments:
#   name := String, name of primer set to calculate maximum intensity for
#   df := DataFrame, dataframe containing all primer set amplification data
def calc_rxn_time_std(name, df):
    curPrimer = df[df['Primer']==name]
    pos_rxns = curPrimer[curPrimer['RxnType']=='+']
    return pos_rxns['RxnTime'].to_numpy().std()

# Function to determine if a reaction is positive or not. Function deteremines a positive according to the following algorithm:
#   1. Determine the intersection point of the reaction and then the reaction reversed.
#   2. Determine the angle from the interxection to time point 0 of both the forward and reverse season.
#   3. If the angle is sufficiently close to 0 (0.95% error in the approximation of the cosine of theta), then mark as negative
#   4. Otherwise, take the derivative and if the lower quantile is greter than 0 (i.e. the signal is not noisy), return as positive
#   5. Otherwise, return negative.
# Arguments:
#   rxnTimeSeries := DataSeries, time series of reaction fluoroescent data.
def determine_pos(rxnTimeSeries):
    errorRateTan = 0.95

    data = rxnTimeSeries
    revData = rxnTimeSeries[::-1]

    idxIntersect = np.where(np.greater(revData.values, data.values)==False)[0][0]

    dataVec = np.array([idxIntersect, data.iloc[idxIntersect] - data.iloc[0]])
    revDataVec = np.array([idxIntersect, revData.iloc[idxIntersect] - revData.iloc[0]])

    magnitude = (np.linalg.norm(dataVec)*np.linalg.norm(revDataVec))
    
    if magnitude == 0:
        return 0
    else:
        theta = math.acos(np.inner(dataVec, revDataVec)/magnitude)

    if((math.cos(theta) > errorRateTan) or math.isnan(theta)):
        return 0
    else:
        if(data.max() > (threshold_perc * MAX_INST_SIGNAL)):
            return 1
        else:
            return 0

# Function to determine true positives based on predicted amplification
# Arguments:
#   name := String, name of primer set to calculate maximum intensity for
#   df := DataFrame, dataframe containing all primer set amplification data
def calc_true_pos(name, df):
    df = df[df['Primer']==name]
    pos_rxns = df[df['RxnType']=='+']
    true_pos_rxns = pos_rxns['ObservedType'].sum()

    return true_pos_rxns

# Function to determine false positives based on predicted amplification
# Arguments:
#   name := String, name of primer set to calculate maximum intensity for
#   df := DataFrame, dataframe containing all primer set amplification data
def calc_false_pos(name, df):
    df = df[df['Primer']==name]
    neg_rxns = df[df['RxnType']=='-']
    true_pos_rxns = neg_rxns['ObservedType'].sum()

    return true_pos_rxns

# Function to calculate reaction time of a single reaction.
# Reaction time is determined as the maximum of the second derivative of the time series.
# Arguments:
#   rxnTimeSeries := DataSeries, time series of reaction fluoroescent data.
def calc_rxn_time(rxnTimeSeries):
    mat = rxnTimeSeries.to_numpy()
    return np.argmax(np.gradient(np.gradient(mat)))

# Function to calculate scored value for a single item
# Arguments:
#   row := DataFrame, results for a single primer set
#   df_results := DataFrame, dataframe for results of all primer sets
#   item := The reaction characteristic that is being scored
def calc_item_score(row, df_results, item):
    item_min = df_results[item].min()
    item_max = df_results[item].max()
    item_range = item_max - item_min
    value = row[item]
    if item_range == 0:
        score = 1
    else:
        if(item == "Intensity_Avg"):
            score = 1 - ((item_max - value)/item_range)
        else:
            score = 1 - ((value - item_min)/item_range)
    weighted_score = score * weights[item]
    return weighted_score

# Function to calculate the overall score for a given primer set in water.
#   row := DataFrame, results for a single primer set
#   df_results := DataFrame, dataframe for results of all primer sets
def calc_overall_score(row, df_results):
    overall_score = 0
    for item in weights.keys():
        if not item=="False_Positives":
            overall_score += calc_item_score(row, df_results, item)
    for i in range(0, replicates):
        overall_score += row['FP_{}'.format(i+1)]
    return overall_score

# Function to calculate the overall score for a given primer set in saliva.
#   row := DataFrame, results for a single primer set
#   df_results := DataFrame, dataframe for results of all primer sets
def calc_overall_score_saliva(row, df_results):
    overall_score = 0
    weights_sal = {'Intensity_Avg': 20, 'Intensity_StdDev': 10, 'RxnTime_Std':40, 'RxnTime_Avg': 30}
    for item in weights_sal.keys():
        overall_score += calc_item_score(row, df_results, item)
    return overall_score

def calc_FP_cardinal(name, df, df_results, n, retValue = 'Score'):

    # Calculate the total number of divisions given a number of replicates. This is just the sum of 1...num of replicates.
    totalDivisions = 0
    for i in range(0, replicates):
        totalDivisions = totalDivisions + (i + 1)

    # Allocate the total number of points for this order of false positives (i.e. 1x, 2x, etc.)
    totalPoints = (weights['False_Positives'] / totalDivisions) * n

    # Select only primer sets that have at least n false positives for consideration in ranking
    subjectPrimerSets = df_results[df_results['False_Positives'] >= n]['PrimerSet']

    if(name not in subjectPrimerSets.values):
        if (retValue == 'Score'):
            return totalPoints
        elif (retValue == 'RxnTime'):
            return np.nan
    else:
        subjectRxnPenalty = df[(df['RxnType']=='-') & (df['ObservedType']==1) & (df['Primer'].isin(subjectPrimerSets))].groupby('Primer')[['RxnPenalty', 'RxnTime']].apply(lambda x: x.sort_values(by='RxnPenalty', ascending=False).iloc[n-1])
        maxSubjectRxnPenalty = subjectRxnPenalty['RxnPenalty'].max()

        if (retValue == 'Score'):
            return (1 - (subjectRxnPenalty['RxnPenalty'][name]/maxSubjectRxnPenalty)) * totalPoints
        elif (retValue == 'RxnTime'):
            return subjectRxnPenalty['RxnTime'][name]

def calc_rxn_penalty(row):
    rxnTimePenalty = 60 - row['RxnTime']
    rxnIntensity = row.drop(['Primer', 'RxnType','ObservedType','RxnTime']).max()
    return rxnIntensity*rxnTimePenalty

# Function to contstruct and return the results table for a set of reactions for a given file.
# Arguments:
#   fileName: String, relative path to the file of interest containing time series data for primer sets.
def getPrimerSummary(fileName):
    [df, primerNames] = format_df(fileName)
    results = pd.DataFrame(data=primerNames, columns=['PrimerSet'])
    results['TruePos'] = results.apply(lambda row: calc_true_pos(row['PrimerSet'], df), axis=1)
    results['Intensity_Avg'] = results.apply(lambda row: 0 if(row['TruePos'] < replicates) else calc_max_avg(row['PrimerSet'], df), axis=1)
    results['Intensity_StdDev'] = results.apply(lambda row:  0 if(row['TruePos'] < replicates) else calc_max_std(row['PrimerSet'], df), axis=1)
    results['RxnTime_Avg'] = results.apply(lambda row:  0 if(row['TruePos'] < replicates) else calc_rxn_time_avg(row['PrimerSet'], df), axis=1)
    results['RxnTime_Std'] = results.apply(lambda row:  0 if(row['TruePos'] < replicates) else calc_rxn_time_std(row['PrimerSet'], df), axis=1)
    results['False_Positives'] = results.apply(lambda row:  0 if(row['TruePos'] < replicates) else calc_false_pos(row['PrimerSet'], df), axis=1)

    for i in range(0, replicates):
        results['FP_{}'.format(i+1)] = results.apply(lambda row: 0 if(row['TruePos'] < replicates) else calc_FP_cardinal(row['PrimerSet'], df, results, (i+1)), axis=1)

    results['Overall_Score'] = results.apply(lambda row:  0 if(row['TruePos'] < replicates) else calc_overall_score(row, results[results['TruePos']==replicates]), axis=1)
    for i in range(0, replicates):
        results['FP_{}_time'.format(i+1)] = results.apply(lambda row: 0 if(row['TruePos'] < replicates) else calc_FP_cardinal(row['PrimerSet'], df, results, (i+1), retValue='RxnTime'), axis=1)
    return results

# Function to return primer names and DataFrame in a dictionary
# Arguments:
#   fileName: String, relative path to the file of interest containing time series data for primer sets.
def get_primer_data(fileName):
    [df, primerNames] = format_df(fileName)
    return {"DataFrame": df, "primerNames": primerNames}

# Function to format time series data into DataFrame
# Arguments:
#   fileName: String, relative path to the file of interest containing time series data for primer sets.
def format_df(filename):
    df = pd.read_excel(filename, header=None)
    primerNames = df.T[0].unique()
    row_names = ['Primer', 'RxnType']
    for i in range(1, 61):
        row_names.append(str(i))
    df.index = row_names
    df = df.T
    df['ObservedType'] = df.apply(lambda row: determine_pos(row.drop(['Primer', 'RxnType'])), axis=1)
    df['RxnTime'] = df.apply(lambda row:  calc_rxn_time(row.drop(['Primer', 'RxnType','ObservedType'])), axis=1)
    df['RxnPenalty'] = df.apply(lambda row: calc_rxn_penalty(row), axis=1)

    return df, primerNames

# Function to score LOD data and return average and standard deviation at each concentration
# Arguments:
#   fileName: String, relative or absolute path to the file of interest containing time series data for primer sets.
# TODO: Update variable names to be more descriptive
def score_LOD(fileName, outputFile):
    # Read in excel file and read sheet name
    file = pd.ExcelFile(fileName)
    sheets = file.sheet_names

    # Concatenate sheets together in DataFrame by assigning different sheets a unique name corresponding to the sheet name.
    df = pd.concat([pd.read_excel(fileName, header=None, sheet_name=sheet).T.assign(sheet_name=sheet) for sheet in sheets])

    # Rearrange columns and assign column headers to DataFrame
    cols = df.columns.tolist()
    cols = cols[-1:] + cols[:-1]
    df = df[cols]
    col_header = ['Primer Set', 'Conc', 'Units', 'Template']
    col_header.extend([str(x) for x in range(1, 61)])
    df.columns = col_header

    # Determine whether reactions are positive or negative and calculate reaction time
    df['ObservedType'] = df.apply(lambda row: determine_pos(row.drop(['Primer Set', 'Conc', 'Units', 'Template'])), axis=1)
    df['RxnTime'] = df.apply(lambda row: calc_rxn_time(row.drop(['Primer Set', 'Conc', 'Units', 'Template'])), axis=1)

    # Create list of each unique concentration and primer set
    totalConcs = df['Conc'].unique()
    totalPrimerSets = df['Primer Set'].unique()

    # Ensure concentration list is in descending order for assignment of LOD
    totalConcs[totalConcs == "NTC"] = -1
    totalConcs = totalConcs[totalConcs[:].astype(float).argsort()[::-1]]
    totalConcs[-1] = "NTC"

    # Insert columns corresponding to LOD and Pathogen
    cols = np.insert(totalConcs, 0, "LOD", axis=0)
    cols = np.insert(cols, 0, "Pathogen", axis=0)

    # Create new dataframe for with all NaN for all concentrations and primer sets
    results_df = pd.DataFrame(np.nan, index=totalPrimerSets, columns=cols, dtype=str)

    # Loop trhough unique primer sets
    for primerSet in totalPrimerSets:
        # Split pathogen from primer set name
        results_df.loc[primerSet, "Pathogen"] = primerSet.split(".")[0]

        #Loop through unique concentrations and initialize flag that we have not yet found LOD
        primerSet_df = df[df['Primer Set'] == primerSet]
        lodFound = False

        # Loop through unique concentrations
        for conc in totalConcs:
            # Get number of replicates for current concentration based on shape of df
            # NOTE: if number of replicates is not consistent across the experiment, this could change the ability to compare across concentrations
            numRep = primerSet_df[primerSet_df['Conc'] == conc].shape[0]

            # Check whether sum of Observed type is greater than num reps for conc and if so, add average reaction rate to new df
            numPos = primerSet_df[primerSet_df['Conc'] == conc]['ObservedType'].sum()

            # Check if the concentration is a No Template Control (NTC) and assign number of positives. 
            # NOTE: STRICT RULE - If false positives are present, the LOD is indeterminant.
            if conc == 'NTC':
                results_df.loc[primerSet, conc] = f"{numPos} of {numRep}"
                if numPos > 0:
                    results_df.loc[primerSet, 'LOD'] = np.nan
            else:
                # If number of positives is greater than number of replicates, add average reaction time to report dataframe along with standard deviation
                if numPos == numRep:
                    results_df.loc[primerSet, conc] = "{:0.2f}".format(primerSet_df[(primerSet_df['Conc'] == conc) & (primerSet_df['ObservedType'] == 1)]['RxnTime'].mean())
                    results_df.loc[primerSet, f"{conc}_STD"] = primerSet_df[(primerSet_df['Conc'] == conc) & (primerSet_df['ObservedType'] == 1)]['RxnTime'].std()
                else:
                    # If number of positives is less than number of replicates, we have found our LOD. Assign the LOD as the last concentration up and 
                    # flag to true
                    if not lodFound:
                        results_df.loc[primerSet, "LOD"] = totalConcs[[i for i, x in enumerate(totalConcs) if x == conc][0] - 1]
                        lodFound = True
                    
                    # Add number of positives out of total replicates to report dataframe
                    results_df.loc[primerSet, conc] = f"{numPos} of {numRep}"

    results_df.to_csv(outputFile, encoding='utf-8')
    # Return report dataframe
    return results_df.sort_values(by=['Pathogen', 'LOD'])

def formatSensSpecData(fileName):
    # Open and read excel sheet names (which are named after the primer sets) before reading in data
    dataFile = pd.ExcelFile(fileName)
    sheetNames = dataFile.sheet_names

    # Read in data, Transpose, and assign column names based on primer set information stored in the sheet name.
    # Assign read and formatted data to a new DataFrame after concatenating all primer sets
    df = pd.concat([pd.read_excel(fileName, header=None, sheet_name=sheet).T.assign(sheet_name=sheet) for sheet in sheetNames])

    # Rearrange columns such that primer set name and concentraiton are first followed by data 
    # and assign column headers to DataFrame
    cols = df.columns.tolist()
    cols = cols[-1:] + cols[:-1]
    df = df[cols]

    # Assign column header names
    col_header = ['Primer Set', 'Conc']
    col_header.extend([str(x) for x in range(1, 61)])
    df.columns = col_header

    # Determine if each reaction is positive or negative and calculate reaction time
    df['ObservedType'] = df.apply(lambda row: determine_pos(row.drop(['Primer Set', 'Conc'])), axis=1)
    df['RxnTime'] = df.apply(lambda row: calc_rxn_time(row.drop(['Primer Set', 'Conc'])), axis=1)

    # Return formatted DataFrame
    return df

def calc_sens_spec(df: pd.DataFrame, concentration: str, primerSet: str) -> pd.DataFrame:
    numTimePoints = 60
    # Set No Template Control (NTC) reactions as predicted negative and all other reactions as predicted positive
    predPos = df[(df['Primer Set'] == primerSet) & (df['Conc'] == concentration)][['Conc', 'ObservedType', 'RxnTime']]
    predNeg = df[(df['Primer Set'] == primerSet) & (df['Conc'] == 'NTC')][['Conc', 'ObservedType', 'RxnTime']]

    # Initialize DataFrame to store sensitivity, specificity, false positive rate, false negative rate, accuracy, and error
    SensSpecCounts = pd.DataFrame(np.nan, index=[i for i in range(1, numTimePoints + 1)], columns=['TP', 'TN', 'FP', 'FN', 'FPR', 'FNR', 'Sensitivity', 'Specificity', 'Accuracy', 'Error'])

    # Initialize all reactions to the length of time points (or full length of reactions) as this will be the reaction time
    # if a given reaction fails to amplify
    predPos.loc[predPos['ObservedType']==0, 'RxnTime']= numTimePoints
    predNeg.loc[predNeg['ObservedType']==0, 'RxnTime']= numTimePoints

    # Loop through each time point and calculate the sensitivity, specificity, false positive rate, 
    # false negative rate, accuracy, and error at that time
    for cutoff_time in range(1, numTimePoints + 1):
        # Count true/false positives/negatives
        true_positives = predPos[(predPos['RxnTime'] <= cutoff_time)].shape[0]
        true_negatives = predNeg[predNeg['RxnTime'] > cutoff_time].shape[0]
        false_positives = predNeg[predNeg['RxnTime'] <= cutoff_time].shape[0]
        false_negatives = predPos[(predPos['ObservedType'] == 0) | (predPos['RxnTime'] > cutoff_time)].shape[0]

        
        # Calculate sensitivity, specificity, false positive rate, false negative rate, accuracy
        sens = true_positives / (true_positives + false_negatives)
        spec = true_negatives / (true_negatives + false_positives)
        false_positive_rate = false_positives / (false_positives + true_negatives)
        false_negative_rate = false_negatives / (false_negatives + true_positives)
        accuracy = (true_positives + true_negatives) / (true_positives + true_negatives + false_positives + false_negatives)
        classificationError = math.sqrt((1-sens)**2 + (1-spec)**2)
        
        # Add to dataframe
        SensSpecCounts.loc[cutoff_time] = [true_positives, true_negatives, false_positives, false_negatives, false_positive_rate, false_negative_rate, sens, spec, accuracy, classificationError]
        
    return SensSpecCounts

def find_optimum_cutoff(df: pd.DataFrame):
    # Find the minimum value of the error and return the corresponding time (located by the index). If multiple maximums exist, return the first one.
    return df[df['Error'] == df['Error'].min()].index[0]

def executeSensSpecAnalysis(fileName, outputFile) -> tuple[pd.DataFrame, pd.DataFrame]:
    # Open and format Sensitivity and Specificity data
    sensSpecData_df = formatSensSpecData(fileName)
    
    # Determine unique concentrations and primer sets
    lodConcs = sensSpecData_df[sensSpecData_df['Conc'] != 'NTC']['Conc'].unique()
    primerSets = sensSpecData_df['Primer Set'].unique()

    # Initialize output DataFrame and set dtype of descriptor columns to string to avoid future warnings
    optimalResults = pd.DataFrame(np.nan, index=[i for i in range(0, len(lodConcs) * len(primerSets))], columns=['Primer Set', 'Concentration', 'Tt', 'TP', 'TN', 'FP', 'FN', 'FPR', 'FNR', 'Sensitivity', 'Specificity', 'Accuracy', 'Error'])
    optimalResults[['Primer Set', 'Concentration', 'Tt']] = optimalResults[['Primer Set', 'Concentration', 'Tt']].astype(str)

    # Initialize output DataFrame for full sensitivity and specificity analysis results
    sensSpecAnalysisResults = pd.DataFrame(columns=['Primer Set', 'Concentration', 'Tt', 'TP', 'TN', 'FP', 'FN', 'FPR', 'FNR', 'Sensitivity', 'Specificity', 'Accuracy', 'Error'])

    # Initialize level counter to track row index for insertion into output DataFrame using iloc
    levelCounter = 0

    # Loop through each concentration and primer set
    for concentration in lodConcs:
        for primerSet in primerSets:

            # Calculate sensitivity and specificity at each time point for the current concentration and primer set (level)
            levelDf = calc_sens_spec(sensSpecData_df, concentration, primerSet)

            # Calculate the optimum cutoff for the current level and the corresponding sensitivity and specificity data
            run_optimumCutoff = find_optimum_cutoff(levelDf)
            optimalSensSpec = levelDf.loc[run_optimumCutoff]

            # Add Primer Set, template concentration, and the optimum cutoff to the optimal sensitivity and specificity data
            rowDescriptors = pd.Series(index=['Primer Set', 'Concentration', 'Tt'], data=[primerSet, concentration, run_optimumCutoff])
            optimalSensSpec = pd.concat([rowDescriptors, optimalSensSpec])

            # Write optimal sensitivity and specificity data to output DataFrame for current level
            optimalResults.iloc[levelCounter] = optimalSensSpec

            # Add Primer Set and template concentration to sensitivity and specificity data
            levelDf[['Primer Set', 'Concentration']] = [primerSet, concentration]

            # Rearrange columns such that primer set name and concentraiton are first followed by data 
            # and assign column headers to DataFrame
            cols = levelDf.columns.tolist()
            cols = cols[-2:] + cols[:-2]
            levelDf = levelDf[cols]

            # Write sensitivity and specificity data for current level to output DataFrame
            # If sensSpecAnalysisResults is not empty, concatenate current level data to it
            if sensSpecAnalysisResults.empty:
                sensSpecAnalysisResults = levelDf
            else:
                sensSpecAnalysisResults = pd.concat([sensSpecAnalysisResults, levelDf])

            # Increment level counter
            levelCounter = levelCounter + 1

    # Write optimal data to "optimalResults" sheet in output and write full data to "data" sheet in output excel file
    with pd.ExcelWriter(outputFile) as writer:
        optimalResults.to_excel(writer, sheet_name='optimalResults')
        sensSpecAnalysisResults.to_excel(writer, sheet_name='data')
    
    # Return optimal sensitivity and specificity data and full sensitivity and specificity data
    return (optimalResults, sensSpecAnalysisResults)
```
