## Supplementary File 3 for "Detection of five viruses commonly implicated with Bovine Respiratory Disease using loop-mediated isothermal amplification": README.md

# PrimerScoring
Contains scripts used to score primer sets.

# Setup
## Dependencies
  - Pandas
  - Numpy
## Installation
Copy the PrimerScore.py file to either a valid location in the search path or to the current director.

# Usage

## Data formatting

### Primer screening data format
The input data file must be in .xls format **NOT .XLSX**. Additionally, the file **MUST** be formmatted in the following manner, with each column representing the data for a single reaction for a specific primer set:
  - First row should contain primer set IDs in the form of *{Target}*.*{SetID}*. For example, orf1ab.2.
  - Second row should contain reaction type annotation. 
    - For negative reaction, put **'-'** in the cell. 
    - For positive reactions, put **'+'** in the cell.
  - Rows three and onward should contain data.
 
**NOTE:**
1. The data should be rectangular; this means that all data is of the same length. This has not been tested for data of different lengths.
2. There should be an equal amount of positive and negatives for a speific primer set. 
3. For proper scoring, each primer set should have the same number of replicates 

### LOD data format
Data for LOD should should be in a `.xls` file. Each tab should correspond to the fluorescent data for one primer set. So, if uniqe primer sets are to be analzyed, there should be four tabs with four unique names.

Each tab should contain a rectangular layout were each column corresponds to the time series of RFU values for a single replicate. The first row should annotate the concentration number, while the second row should annotate the concentration units. For instance, if the concentration for a replicate was 25 copies per reaction (cps/rxn), the first row would be 25 and the second row would be cps/rxn. The third row should contain the template for that replicate. For NTC reactions, the concentration units row should be empty. The following $n$ rows should be the fluorescent RFU values at each time point for that replicate.

### Sens/Spec data format
Similar to Sens/Spec data, data for sensitivity and specificity analysis should be contained in one `.xls` file. Each tab should correspond to the fluorescent data for one primer set. So, if uniqe primer sets are to be analzyed, there should be four tabs with four unique names.

Each tab should contain a rectangular layout where each column corresponds to the time series of RFU values for a single replicate. The first row should contain the concentration designation. Do not indicate replicates, as the code will automatically determine the number of replicates. Then the following $n$ rows should be the flourescent RFU value at each time point for that replicate. 

Data should be rectangular. As before, each concentration and NTC reactions should have the same number of replicates.


 ## Execution
 ### Importing
 If the PrimerScore.py file is located in a valid path or the current directory, simply execute the following line to import the module.
 ```
 import PrimerScore
 ```
 
 ### Initialization
 To use the primer scoring method, the initialization method must first be called to set weights and tolerances.
 
 To initialize with default weights, simply execute the following line of code:
 ```
 PrimerScore.intialize()
 ```
 This will initialize weights to the following value:
 
 |  Metric  | Weight  |
 |:---:|:---:|
 |  Average Maximum Intensity | 5 |
 |  Max. Intensity Std. Dev.  | 5 |
 |  Reaction Time Std. Dev.   | 10|
 |  Average Reaction Time     | 20|
 |  Number of False Positives | 60|
 
 Additionally, the thresholds will be set at:
 |  Threshold | Value |
 |:---:|:---:|
 |  Exponential Phase  |  3000  |
 |  Plateau Phase      |  200   |
 |  Positive Reaction Threshold | 0.9 |
 |  False Positive Threshold    | 0.2 |
 
Finally, the number of replicates will be set at 4:
 |  Parameter | Value |
 |:---:|:---:|
 |  Number of Replicates  |  4  |
 |  Instrument Saturation/Maximum Intensity      |  140000   |
 |  Positive Threshold Percentage | 0.1 |
 
The thresholds above are presently non-functional, but are retained for potential future use. Reaction time is determined as the maximum of the second derivative of the intensity over time.  

 
 To use custom weightings, execute the following line of code with the given arguments.
 
 ```
 PrimerScoring.initialize(set_weights, set_thresholds, set_replicates, set_instrumentMax, set_threshold_perc)
 ```
The arguments in the above expression are defined as:
- set_weights: An array containing weights for the following metrics in the following order:
  - Avg. Max. Intensity
  - Max. Intensity Std. Dev.
  - Reaction time Std. Dev.
  - Avg. Reaction Time
  - Number of False Positives
  - **THE ARRAY MUST CONTAIN WEIGHTS FOR ALL OF THE ABOVE METRICS.**
- set_thresholds: An array contianing weights for the threshold tolerances of the exponential and plateau phases, positive reaction threshold, and false positive threshold, respectively. Whereas this value can be set, it is currently not in use.
- set_replicates: An integer value greater than 3 for the number of replicates. **Must be the same for all primer sets scored.**

An example to initialize to default settings as above would be as follows:
```
PrimerScoring.intialize([5,5,10,20,60], [3000, 200, 0.9, 0.2], 4, 140000, 0.1)
```

### Scoring
To calculate the scores for each primer, execute the following line of code with the given arguments:

```
PrimerScore.scorePrimers(primerData, output)
```
The arguments in the above line of code are defined as follow:
  - primerData: Path (relative or absolute) or file name of excel (.xls) file containing primer data formated properly as outlined above.
  - output: Output excel file (.xlsx) name or path containing primer scores and metrics. Please include .xslx extension.
 
 An example to take an input file "WaterScore.xls" and write values to "PrimerScores_Water.xlsx" is as follows:
 ```
 PrimerScore.scorePrimer('WaterScore.xls', 'PrimerScores_Water.xlsx')
 ```
 
 ### Output
 After scoring primers as outlined above, the output file will have the following column headers:
  - Primer Set: Column contains Primer Set ID (copied from input)
  - TruePos: Column contains the number of true positives (completed reaction)
  - Intensity_Avg: Calculated average maximum intensity. 
  - Intensity_StdDev: Calculated maximum intensity standard deviation.
  - RxnTime_Avg:  Calculated average reaction time.
  - RxnTime_StdDev: Calculated reaction time standard deviation.
  - False_Positives:  Number of total false positives.
  - FP_...: There will be a false positive column for each replicate containing the contribution of that false positive to the overall score.
  - Overall Score: Calculated overall score for each primer set.
 
# Methodology

## Nomenclature
The weights input during the initialization procedue are inidcated by $\omega_{x}$ where $x$ is a given feature and can be $\bar{I}$ for average maximum intensity, $\sigma_{I}$ for maximum intensity standard deviation, $\sigma_{t_{rxn}}$ for reaction time standard deviation, $\bar{t_{rxn}}$ for average reaction time, FP for false positives.

## Positive amplification detection
LAMP amplification reactions typically produce a sigmoidal amplification; however, given fluorometric methods typically have some background auto-flourescence or variable response over time, it is not sufficient to simply check for an increase in signal over time. To this end, the following methodology was used to determine a "positive amplification", regardless of designation (true positive or No Template Control (NTC)):

1. The series containing the fluorometric reads over time was duplicated and reversed.
2. The intersection of the series and the reversed series was determined by the time point at which the forward time-series first exceeded the reverse time-series.
3. Two vectors were created, one for the forward time series and one for the reverse, using the following definition:
    - $\overrightarrow{\text{Forward/Reverse Data Vector}} = <\text{Time of Intersection }, y_{Intersection} - y_0>$
4. The cosine between to two vectors was calculated
    - $\theta = \arccos(\frac{\overrightarrow{\text{Forward}}\cdot\overrightarrow{\text{Reverse}}}{\|\overrightarrow{\text{Forward}}\| \| \overrightarrow{\text{Reverse}}\|})$
5. Using the understanding that $\cos(x) \approx 1$ if $x \approx 0 $, we check that the $\cos{\theta} > 0.95$ assuming a 95% error in the approximatation. Essentially, these steps are checking to see if our data is "flat" or relatively constant within error. 
    - If it is, we will return that this reaction is not a positive amplification
    - If it is not, continue to step 6.
6. Check if the maximum of the time series is above some threshold percentage of the maximum fluorescent intensity of the instrument, and if it does return a positive amplification.
    - Default parameters for maximum fluorescent intensity taken on an Analytik-Jena qTower 3G is approximately 140000 Relative Fluorescence Units.
    - Default threshold percentage is 10%. 

## False negatives
If any reaction labelled "+" is detected as a negative amplification, it is labelled as a false negative. Given that there are no checks on the number of replicates a user inputs in this script to ensure that the statistical "power" for averages and standard deviations is comparable across all compared primer sets for scoring, any false negative reaction automatically results in a score of 0 for that primer set and it is removed from consideration for scoring metrics.

## Reaction time
Reaction time is determined as the maximum of the 2nd derivative of the fluorescent time series data. This is implemented using [`numpy.gradient`](https://numpy.org/devdocs/reference/generated/numpy.gradient.html).

Penalties are incurred for later reaction times, thus the feature that is used for scoring is the value $60 - t_{rxn}$

## Average and standard deviation of reaction features
All averages and standard deviations are calculated from individual reaction metrics over all replicates that are labelled as positive reactions in input data and detected as positives.


## Weighting of False Positives
False positives are undesireable in the context of the developed diagnostics and hence are weighted very strongly to filter out primer sets that produce false positives. Additionally, it is possible to have a one-off or rare occurrence false positive due to operator error or contamination, rather than an inherent interaction of the primers in the primer set, which should be strongly discouraged. When a reaction is labelled as "-" in the input data, but is detected as a positive during the positive amplification detection, it is labelled as a false positive.  

To this end, a "progressive" penalty for increasing occurrence of false positives was implemented to select for primer sets with less "persistent" false positives. This is accomplished by dividing the total weight allocated to false positives during initialization by a factor, $\alpha = \omega_{FP} / \sum_{i=1}^{n} (i)$ where $n$ is the number of replicates. This factor is then increased linearly for increasing numbers of False Positives in a given reaction by multiplying the false positive order by $\alpha$ (i.e. the first positive receives a penalty of $\alpha$, the second false positive receives a penalty of $2 \cdot \alpha$, etc.). 

Furthermore, an overall "reaction penalty" $\left( \Omega \right)$ is calculated by multiplying the maximum intensity of a replicate by the reaction time of a replicate. This penalty is on a per reaction basis, *not* averaged across all replicates. In this manner, if a reaction is detected as a false positive, but only amplifies a small amount compared to other reactions, it is not penalized as heavily. Likewise, late stage false postives are penalized less. 

Lastly, for all false positive calculations, primer sets being compared must all have at least $i$ false positives. Therefore, a primer set with 3 primer sets will only be compared against all primer sets also with at least 3 false positives. This analysis is conducted for all numbers between 1 and the number of replicates. The value that is weighted is the reaction penalty for the $i$ th false positive when all reaction penalties are sorted in ascending order for each primer set. The resulting value from each primer set is then compared and weighted in a manner similar to other reaction features.

## Scoring

Once all primer sets have had primer set performance features calculated, an overall score is calculated by ordering primer sets and weighting a primer sets individual score according to its placement in the resulting order amongst all primer sets. This is achieve using the following formulation:

$S_k = \omega_{\bar{I}} \cdot \left( 1 - \frac{max \left( \bar{I} \right) - \bar{I}_k}{Range \left( \bar(I) \right)} \right) + \sum _x \omega _x \cdot \left( 1 - \frac{\text{min}(x) - x_k}{\text{Range}(x))} \right) + \sum _{i=0} ^n \left( i  \cdot \alpha \cdot \left( 1 - \frac{\phi \left( \Omega \right)_i}{\text{max}(\phi (\Omega))_i} \right) \right)$

where $k$ is a given primer set, $x$ indicates a given feature, $\omega_x$ is the weight allocated to feature $x$, $x_k$ is the feature value for primer set $k$, $\alpha$ is the false positive weighting factor, $n$ is the number of replicates, $\Omega_i$ is the reaction penalty for reaction $i$, and $\phi$ is the set of reaction penalities for each *false positive* reaction for a specific primer set ordered from smallest to largest such that an element $\Omega \in \phi$ if a given primer set has at least $i$ false positives, and $\text{max}(\phi (\Omega))_i $ is the maximum value of the $i$th reaction penalty of each primer set containing at least $i$ false positives.

## LOD Determination and Scoring

The Limit of Detection for a primer set was determined by observing the lowest concentration, $C$, at which all replicates, $n$, at that concentration level amplified and all replicates of all concentrations higher than that level. All reactions are determined to be positive using the same positive detection methodology as above.

If any false positives (i.e. deteremined positive amplifications in designated NTC reactions) are detected, then the LOD is indeterminant for that primer set.

## Sensitivity and Specificity Analysis

Sensitivity and specificity analyses were typically conducted at 2x and 1x the concentration of LOD (but this is not all the case). In any event, the current process identifies the number of "distinct" concentrations along with a common negative control set of reactions and determines analytical sensitivity and specificity on each of those distinct concentrations.

For all input reactions, the algorithm determines any reaction that is *not* an NTC reaction to be a predicted positive reaction. All NTC reactions are predicted negatives. Positive reaction detection is then applied to all reactions to determine observed positive and observed negative reactions along with reaction times. Observed positves were further designated at a given time point, $t_{set}$, only if $t_{set} < t_{rxn}$. Otherwise, the reaction was counted as an observed negative at that time point.

For each time point, $t_set$ in the LAMP reaction, the following were calculated.
- True Positive  (TP): Predicted Positive and Observed Positive
- True Negative  (TN): Predicted Negative and Observed Negative
- False Positive (FP): Predicted Negative and Observed Positive
- False Negative (FN): Predicted Positive and Observed Negative

From these counts, the following were then calculated for each tiem point:
- $\text{Sens} = \frac{\text{TP}}{\text{TP} + \text{TN}}$
- $\text{Spec} = \frac{\text{TN}}{\text{TN} + \text{FP}}$
- $\text{FPR}  = \frac{\text{FP}}{\text{FP} + \text{TN}}$
- $\text{FNR}  = \frac{\text{FN}}{\text{FN} + \text{TP}}$
- $\text{Acc}  = \frac{\text{TP} + \text{TN}}{\text{TP} + \text{TN} + \text{FP} + \text{FN}}$
- $\text{Classification Error} = \sqrt{\left( 1 - \text{Sens}\right)^2 + \left( 1 - \text{Spec}\right)^2}$

The reaction time which minimized the classification error was then chosen as the optimum point for reporting final primer set Sensitivity and Specificity metrics.
