## Supplementary File 3 for "Detection of five viruses commonly implicated with Bovine Respiratory Disease using loop-mediated isothermal amplification": DigitalAnalysis.ipynb

```
# %% [markdown]
# Digital Assay Analysis

This folder is meant to serve as an example of how to call and interact with the `DigitalAssayAnalysis` module. 

## License
Copyright (C) 2025 Purdue Research Foundation. This work is protected under GNU Affero General Public License Version 3 with Commons Clause. Please see the `LICENSE` file for more information.

# %% [markdown]
## Setup

# %% [code]
import DigitalAssayAnalysis
from os import walk
from os.path import join

# %% [markdown]
## Calculation of concentration for fine LODs using dPCR

# %% [code]
inputFilePath = ""
outputFilePath = ""
summary_df, results_df = DigitalAssayAnalysis.calc_dPCRStats(inputFilePath, outputFilePath, plate_type='26k')
display(summary_df, results_df)

# %% [markdown]
## Calculation of background sample concentration using dPCR

# %% [code]
fileDir = ""

# For background data, we will simply loop through each file, one per virus and run the analysis
fileNames = [(dirpath,f) for (dirpath, dirnames, filenames) in walk(fileDir) for f in filenames]

# For each file we found in our directory for the input background data files, see if it is in the input folder, and run the analysis
for file in fileNames:
    # Split directory and file name to get directory path and file name
    dir = file[0]
    fn = file[1]

    # Check if the file is in the input folder
    if(dir.split("/")[-1] == 'input'):
        # Join the directory and file name to get the full file path
        fInPath = join(dir, fn)

        # Determine the output directory and file name
        outputDirIdx = dir.rfind("/")
        fileNameIdx = fn.rfind(".")
        fOutPath = f"{dir[:outputDirIdx]}/output/{fn[:fileNameIdx]}.xlsx"

        # Run analysis. For this specific setup, the background analysis was conducted on a 96-well, 8.5k plate with hyperwelling every E3 - H3 and E6 - H6.
        # Additionally, since we are interested in the copies per microliter as opposed to copies per reaction, we will set the template_rxn_vol to 1
        # and interpret the concentration as copies per microliter in the output file where it indicates copies per reaction.
        DigitalAssayAnalysis.calc_dPCRStats(fInPath, fOutPath, plate_type='8.5k', makeSummary = False, rxn_dilution_factor = 4 , template_rxn_vol = 1, hyperwellGroups=[['E3','F3','G3','H3'], ['E6', 'F6', 'G6', 'H6']])

# %% [code]
```
