## Supplementary File 3 for "Detection of five viruses commonly implicated with Bovine Respiratory Disease using loop-mediated isothermal amplification": DigitalAssayAnalysis.py

```
#!/usr/bin/env python3
# -*- coding: utf-8 -*-
'''Digital Assay Analysis.py is a python module for analyzing digital assay data acquired with Qiagen QIAcuity Software'''

__author__ = 'Josiah Levi Davidson'
__contact__ = 'Mohit S. Verma'
__email__ = ''
__version__ = '1.0.0'
__status__ = 'Development'
__copyright__ = "Copyright (C) 2025 Purdue Research Foundation"
__license__ = "GNU Affero General Public License Version 3 with Commons Clause"

#### BEGIN IMPORTS

import pandas as pd
import numpy as np
from scipy.stats import t

# Set partition counts based on manufacturer specifications
# (Page 102 of the QIAcuity App. Guide, 
# https://www.qiagen.com/us/resources/resourcedetail?id=5d19083d-fa10-4ed2-88a0-2953d9947e0c)
partition_count = {'8.5k': 8510, '26k': 26384}

def calc_absoluteQuant(row, rxn_dilution_factor, template_rxn_vol, plate_type):
    # If there is no positive partitions, return 0 for everything
    if row['Positive'] == 0:
        return [0,0,0,0,0,0,0,0]
    else:
        # Calculate the number of hyperwells based on the approximate devision for a full well
        # NOTE: This will fail if the imaging step fails to identify well based on the reference dye
        numHyperwells = round(row['Total']/partition_count[plate_type])

        # Calculate average partition volume        
        partitionVol = row['Volume']/(numHyperwells * partition_count[plate_type])

        # Calculate percent positive partitions
        p = row['Positive']/row['Total']

        # Calculate lambda and confidence around lambda
        lambda_center = -np.log(row['Negative']/row['Total'])
        lambda_low = -np.log((1 - p + 1.96*np.sqrt(p*(1-p)/row['Total'])))
        lambda_high = -np.log((1 - p - 1.96*np.sqrt(p*(1-p)/row['Total'])))

        # Determine lambda range
        lambda_range = lambda_high - lambda_low

        # If lambda range is greater than lambda center, use lambda high for percentage, else use lambda range
        # NOTE: I was not able to determine why this substitution was made, but it was nevertheless necessary
        # to produce results identical to that of the QIAcuity software suite
        if lambda_range > lambda_center:
            lambda_perc = lambda_high/(2*lambda_center)*100
        else:
            lambda_perc = lambda_range/(2*lambda_center) * 100
        
        # Calculate concentration per µL
        conc_dPCRRxn_uL = (lambda_center / partitionVol)

        # Calculate concentration per reaction and confidence bounds
        conc_avg = (lambda_center / partitionVol) * rxn_dilution_factor * template_rxn_vol
        conc_low = (lambda_low / partitionVol) * rxn_dilution_factor * template_rxn_vol
        conc_high = (lambda_high / partitionVol) * rxn_dilution_factor * template_rxn_vol

        # Return all concentration values
        return [conc_dPCRRxn_uL,conc_avg, conc_low, conc_high, lambda_perc, lambda_center, lambda_low, lambda_high]

def calc_avgQuant(rows, rxn_dilution_factor, template_rxn_vol, plate_type, alpha_crit = 0.05):
        # Calculate the number of hyperwells based on the approximate devision for a full well
    # NOTE: This will fail if the imaging step fails to identify well based on the reference dye
    numHyperwells = (rows['Total']/partition_count[plate_type]).round()

    # Calculate average partition volume
    partVol = rows['Volume']/partition_count[plate_type]

    # Calculate ideal partition count for plate type
    wellPartCount = numHyperwells * partition_count[plate_type]

    # Calculate the average concentration per microliter
    a = rows['Volume'].mul(rows['Conc(cps/uL)'], axis=0).div(wellPartCount)
    conc_avg_uL = a.sum()/(partVol).sum()

    # Calculate the average concentration per reaction
    a = rows['Volume'].mul(rows['Conc(cps/rxn)'], axis=0).div(wellPartCount)
    conc_avg_rxn = a.sum()/(partVol).sum()

    # Calculate the t-critical value based on number of replicates
    t_crit = t.ppf(1-(alpha_crit/2), df=rows.shape[0])

    # Calculate the standard deviation of the concentration per microliter based on the t-critical value
    std_conc_uL = (((rows['Conc(cps/uL)'] - conc_avg_uL)**2).sum()/(rows.shape[0]-1))**0.5

    # Calculate high and low concentration per microliter bounds for a (1-alpha)% confidence interval
    conc_avg_uL_low = conc_avg_uL - t_crit*std_conc_uL/(rows.shape[0]**0.5)
    conc_avg_uL_high = conc_avg_uL + t_crit*std_conc_uL/(rows.shape[0]**0.5)

    # Calculate the standard deviation of the concentration per reaction based on the t-critical value
    std_conc_rxn = (((rows['Conc(cps/rxn)'] - conc_avg_rxn)**2).sum()/(rows.shape[0]-1))**0.5

    # Calculate high and low concentration per reaction bounds for a (1-alpha)% confidence interval
    conc_avg_rxn_low = conc_avg_rxn - t_crit*std_conc_rxn/(rows.shape[0]**0.5)
    conc_avg_rxn_high = conc_avg_rxn + t_crit*std_conc_rxn/(rows.shape[0]**0.5)    

    return [conc_avg_uL, conc_avg_uL_low, conc_avg_uL_high, conc_avg_rxn, conc_avg_rxn_low, conc_avg_rxn_high ]

def calc_dPCRStats(fileInputPath, fileOutputPath, plate_type = '26k', rxn_dilution_factor = 4, template_rxn_vol = 5, makeSummary = True, hyperwellGroups = []) -> tuple[pd.DataFrame, pd.DataFrame]:
    # Read in data file without limiting memory usage, as file is large. 
    """
    Reads in a Digital PCR file from QIAcuity and calculates the concentrations based on the sample name.
    The calculations are done on a per well basis, and the results are then averaged over the wells within a sample
    to generate a summary dataframe. The results are then written out to an excel file with the sample name and
    calculated concentrations.

    Parameters:
    fileInputPath (str): The path to the Digital PCR file from QIAcuity.
    fileOutputPath (str): The path to the output excel file.
    plate_type (str): The type of plate used. Default is '26k'.
    rxn_dilution_factor (int): The dilution factor for the reaction. Default is 4.
    template_rxn_vol (int): The volume of the reaction. Default is 5.
    makeSummary (bool): Whether or not to make a summary dataframe. Default is True.
    hyperwellGroups (list): A list of lists, where each sublist is a group of wells to be hyperwelled together.

    Returns:
    tuple[pd.DataFrame, pd.DataFrame]: A tuple containing the summary dataframe and the full dataframe.
    """
    df = pd.read_csv(fileInputPath, low_memory=False, header=1)

    # Loop through each group that we will hyperwell together
    for hGroup in hyperwellGroups:
        # Merge wells in group together based on Well identifier
        groupVols = df[df['Well'].isin(hGroup)][['Well', 'Cycled volume']].drop_duplicates()
        # Total yperwell volume is the sum of each well in group
        totalVol = groupVols['Cycled volume'].sum()

        # Assign total volume and name to each well in hyperwell to complete "merge"
        df.loc[df['Well'].isin(hGroup), 'Cycled volume'] = totalVol
        df.loc[df['Well'].isin(hGroup), 'Well'] = hGroup[0]

    # Set threshold as the first unique value. Common threshold should be used.
    threshold = df['Threshold'].unique()[0]

    # Create summary table based on unique sample names
    sampleList = df['Sample'].unique()

    # Create rows for resulting concentration in each row.
    rows = df['Well'].unique()
    cols = ['Sample', 'Total', 'Positive', 'Negative', 'Conc', 'LowConc', 'HighConc', 'Volume',
            'Conc(cps/uL)', 'Conc(cps/rxn)', 'lambda_perc', 'lambda_center', 
            'lambda_low', 'lambda_high']
    
    # Initialize summary dataframe
    resultsDf = pd.DataFrame(0, index=rows, columns=cols)

    resultsDf = resultsDf.astype({cols[0] : str})
    resultsDf = resultsDf.astype({key: int for key in cols[1:4]})
    resultsDf = resultsDf.astype({key: float for key in cols[4:]})

    #sumDf.loc[:, ['Conc(cps/uL)', 'Conc(cps/rxn)', 'LowConc', 'HighConc', 'lambda_perc', 'lambda_center', 'lambda_low', 'lambda_high']].astype(float)
    # Loop through each well and calculate stats
    for well in rows:
        # Select data only for current well
        currentWellData = df[(df['Well'] == well) & (df['Is invalid'] == "0")]

        # Set sample name for current well. Whereas unique() should only return one value, we index in case
        # any typo or edits occured so as not to introduce errors.
        resultsDf.loc[well, 'Sample'] = currentWellData['Sample'].unique()[0]

        # Determine invalid well counts based on QIAcuity invalid designation, and then positive and negative annotations
        # based on amplification being located above or below threshold.
        resultsDf.loc[well, 'Total'] = currentWellData[currentWellData['Is invalid'] == "0"]['Is invalid'].count()
        resultsDf.loc[well, 'Positive'] = currentWellData[currentWellData['RFU'] >= threshold]['RFU'].count()
        resultsDf.loc[well, 'Negative'] = currentWellData[currentWellData['RFU'] < threshold]['RFU'].count()

        # Set current well volume, which should be unique but in the event mutation of data occurred, select the first value
        resultsDf.loc[well, 'Volume'] = currentWellData['Cycled volume'].unique()[0]
        
        # Calculate concentrations and concentration range
        resultsDf.loc[well, ['Conc(cps/uL)', 'Conc(cps/rxn)', 'LowConc', 'HighConc', 'lambda_perc', 'lambda_center', 
                             'lambda_low', 'lambda_high']] = calc_absoluteQuant(resultsDf.loc[well, :], rxn_dilution_factor,  template_rxn_vol, plate_type)

    for sample in sampleList:
        #currentSampleData = df[df['Sample'] == sample]

        resultsDf.loc[resultsDf[resultsDf['Sample'] == sample].index, 
                  ['AvgConc(uL)', 'AvgConc(uL) Low', 'AvgConc(uL) High', 'AvgConc(rxn)', 'AvgConc(rxn) Low', 'AvgConc(rxn) High']] = \
                    calc_avgQuant(resultsDf[resultsDf['Sample'] == sample], 4, 5, plate_type)

    resultsDf = resultsDf[['Sample', 'Conc(cps/rxn)', 'LowConc', 'HighConc', 'AvgConc(rxn)', 'AvgConc(rxn) Low', 
                  'AvgConc(rxn) High', 'Total', 'Positive', 'Negative', 'Conc(cps/uL)', 'lambda_perc', 
                  'lambda_center', 'lambda_low', 'lambda_high', 'AvgConc(uL)', 'AvgConc(uL) Low', 
                  'AvgConc(uL) High', 'Volume']]
    
    summaryDf = pd.DataFrame()

    if makeSummary:
        # Create a summary dataframe that only shows whole number copies per reaction for each level
        summaryDf = resultsDf[['Sample', 'AvgConc(rxn)', 'AvgConc(rxn) Low', 'AvgConc(rxn) High']].drop_duplicates()

        # For average values, round down if positive and round up if negative
        summaryDf.loc[summaryDf[summaryDf['AvgConc(rxn)'] < 0].index, 'AvgConc(rxn)'] = np.ceil(summaryDf[summaryDf['AvgConc(rxn)'] < 0]['AvgConc(rxn)'])
        summaryDf.loc[summaryDf[summaryDf['AvgConc(rxn)'] > 0].index, 'AvgConc(rxn)'] = np.floor(summaryDf[summaryDf['AvgConc(rxn)'] > 0]['AvgConc(rxn)'])

        # Round the high and low confidence intervals such that the intervals are being contracted not expanded
        # by the rounding operation\
        summaryDf['AvgConc(rxn) Low'] = np.ceil(summaryDf['AvgConc(rxn) Low'])
        summaryDf['AvgConc(rxn) High'] = np.floor(summaryDf['AvgConc(rxn) High'])

        summaryDf = summaryDf.astype({key: int for key in summaryDf.columns[1:]})

        # Write summary data to "optimalResults" sheet in output and write full data to "data" sheet in output excel file
        with pd.ExcelWriter(fileOutputPath) as writer:
            summaryDf.to_excel(writer, sheet_name='Summary')
            resultsDf.to_excel(writer, sheet_name='data')
    else:
        # Write full data to "data" sheet in output 
        with pd.ExcelWriter(fileOutputPath) as writer:
            resultsDf.to_excel(writer, sheet_name='data')

    # Return dataframes
    return summaryDf, resultsDf
```
