## Supplementary File 3 for "Detection of five viruses commonly implicated with Bovine Respiratory Disease using loop-mediated isothermal amplification": README.md

 # Digital Assay Analysis

This folder contains the library used to analyze dPCR data from the QIAcuity software suite.

## Dependencies

This file requires the following libraries to run:
- Numpy
- Pandas
- Scipy

## Input data

This script requires raw dPCR data from the QIAcuity control software in the form of `.csv` files. See the software manual for the proper wasy to export raw data from the QIAcuity software suite. Additionally, for every partition, the name and well must be defined. Therefore, it may be necessary to copy sample names and well locations down the rows of the raw data file before running the script.

## Usage

The script is intended to be used in a jupyter notebook by calling 

`import DigitalAssayAnalysis`.


Then, to run the script simply call `calc_dPCRStat(...)` with the following input arguments:
- (mandatory) fileInputPath (`string`): Path to the raw data `.csv` file exported from the QIAcuity software.
- (mandatory) fileOutputPath (`string`):  Desired output path for `.xls` file.
- (optional) plate_type (`string`): Either of `{"8.5k", "26k"}`. Default: `"26k"`
- (optional) rxn_dilution_factor (`int`): Dilution factor for given by $V_{rxn}/V_{Template}$. Default:  `4`
- (optional) template_rxn_vol (`int`): Reaction volume of template, in $\mu L$. Default: `5`
- (optional) makeSummary (`boolean`): Return summary table of all samples. Default: `True`
- (optional) hyperwellGroups (array of array of strings): Wells which should be hyperwelled together in an array. Default: `[]` Example: If well A1, A2, and A3 are hyperwelled together, the input is `[["A1", "A2", "A3"]]`
## Methodology

This script largely uses the methodology from page 102 of the [QIAcuity Software manual](
https://www.qiagen.com/us/resources/resourcedetail?id=5d19083d-fa10-4ed2-88a0-2953d9947e0c). A few modifications concerning the calculation of averages and hyperwelling have been used and are described below.

### Absolute Quantification

Quantification of template in digital assays is conducted accoring to a Poisson distribution where the average concentration is given by: 
$C_{avg, well} = \frac{\lambda_{avg}}{V_{part}}\cdot F_{Rxn} \cdot V_{Temp}$

where 

$\lambda_{avg} = -\ln{\left( \frac{N_{neg}}{N_{Total}}\right)}$

and $N_{Total}$, $N_{Neg}$, and $N_{Pos}$ are the number of Total partitions, number of negative partitions, and the number of positive partitions, respectively. $F_{rxn}$ is the reaction dilution factor. $V_{Temp}$ is the volume of template added to the reaction. $V_{part}$ is the partition volume and is given by the: $V_{loaded} / N_{part, ideal}$. $N_{Part, ideal}$ is the number of partitions in a "perfect" well array as reported by Qiagen and is 8510 for 8.5k plates and 26,384 for 26k plates.

### Confidence Intervals on Absolute Quantification
The 95% confidence interval around this is given by:

$C_{95\\%} = \left( \frac{\lambda_{low}\cdot F_{Rxn} \cdot V_{rxn}}{V_{part}}, \frac{\lambda_{high}\cdot F_{Rxn} \cdot V_{rxn}}{V_{part}}    \right) $

where 

$\lambda_{low} =-\ln \left( (1-p) + 1.96 \cdot \sqrt{\frac{p\cdot(1-p)}{N_{Total}}} \right) $

and 

$\lambda_{high} =-\ln \left( (1-p) - 1.96 \cdot \sqrt{\frac{p\cdot(1-p)}{N_{Total}}} \right) $.

All other values remain the same as before.

### Average per sample

Average are calculated by combining samples with the same name in the raw data `csv`. Therefore, to count $n$ replicates, each replicate must have the same name. If wells are hyperwelled, ensure that you keep the names the same, but be sure you input the correct wells in the list format for the arguments. 

Once samples have been determined, average partition volume is calculated by:

$V_{part} = \frac{V_{partArr}}{N_{part,Ideal}}$

Then, the ideal partition count is calculated by multiplying $N_{part,Ideal}$ by the number of hyperwells.

For each replicate, the concentration per $\mu L$ is calculated as:
$C_{cps/\mu L, part} = V_{partArr}\cdot C_{cps/\mu L, well}$

Then, the average is calculated by 

$C_{avg, sample} = \sum{C_{cps/\mu L, part}}/ \sum{V_{part}}$

The average concentration per reaction is calculated in the same way, but replacing $C_{cps/\mu L, well}$ with $C_{cps/rxn,well}$. 

### Confidence Intervals on Sample Averages

Confidence intervals around sample averages are calculated using the critical value for a 2-tailed t-test for the number of replicates, $n$. 

The standard deviation of the average concentration, $\sigma (C)$ is given by:

$\sigma (C) = \sum \left ({C_{cps/\mu L, well}} - C_{avg, sample} \right)^2 / \sqrt{n - 1} $

Then, upper and lower concentrations are determed using the $t_crit$ value, such that 95% confidence interval on the average quantification is given by:

$C_{avg, 95\\%} = \left( \frac{C_{avg, sample} - t_{crit}\cdot\sigma (C)}{\sqrt{n}}\, \frac{C_{avg, sample} + t_{crit}\cdot\sigma (C)}{\sqrt{n}}   \right)$

Averages for copies per reaction are calcualted the same way, except replacing $C_{cps/\mu L}$ with $C_{cps/rxn}$.

<p xmlns:cc="http://creativecommons.org/ns#" xmlns:dct="http://purl.org/dc/terms/"><span property="dct:title">&copy; 2025 - Purdue Research Foundation. Reaction Analysis Library Documentation</span> by <span property="cc:attributionName">Josiah Davidson</span> is licensed under <a href="https://creativecommons.org/licenses/by-nc/4.0/?ref=chooser-v1" target="_blank" rel="license noopener noreferrer" style="display:inline-block;">CC BY-NC 4.0<img style="height:22px!important;margin-left:3px;vertical-align:text-bottom;" src="https://mirrors.creativecommons.org/presskit/icons/cc.svg?ref=chooser-v1" alt=""><img style="height:22px!important;margin-left:3px;vertical-align:text-bottom;" src="https://mirrors.creativecommons.org/presskit/icons/by.svg?ref=chooser-v1" alt=""><img style="height:22px!important;margin-left:3px;vertical-align:text-bottom;" src="https://mirrors.creativecommons.org/presskit/icons/nc.svg?ref=chooser-v1" alt=""></a></p>
