## Supplementary File 3 for "Detection of five viruses commonly implicated with Bovine Respiratory Disease using loop-mediated isothermal amplification": README.md

# Reaction Analysis Library

This repo contains a variety of scripts used by the Verma lab for evaluation and analysis of reaction data. At present, the following scripts are included:

- [DigitalAssays](https://github.itap.purdue.edu/VermaLab/ReactionAnalysisLibrary/tree/main/DigitalAssays)

<p xmlns:cc="http://creativecommons.org/ns#" xmlns:dct="http://purl.org/dc/terms/"><span property="dct:title">&copy; 2025 - Purdue Research Foundation. Primer Analysis Documentation</span> by <span property="cc:attributionName">Josiah Davidson</span> is licensed under <a href="https://creativecommons.org/licenses/by-nc/4.0/?ref=chooser-v1" target="_blank" rel="license noopener noreferrer" style="display:inline-block;">CC BY-NC 4.0<img style="height:22px!important;margin-left:3px;vertical-align:text-bottom;" src="https://mirrors.creativecommons.org/presskit/icons/cc.svg?ref=chooser-v1" alt=""><img style="height:22px!important;margin-left:3px;vertical-align:text-bottom;" src="https://mirrors.creativecommons.org/presskit/icons/by.svg?ref=chooser-v1" alt=""><img style="height:22px!important;margin-left:3px;vertical-align:text-bottom;" src="https://mirrors.creativecommons.org/presskit/icons/nc.svg?ref=chooser-v1" alt=""></a></p>
